# Macroevolutionary shifts in post-hatching ontogeny and the origin of craniofacial disparity in fowl (Aves: Galloanserae)

**DOI:** 10.64898/2026.06.12.731877

**Authors:** Bassel Arnaout, Guillermo Navalón, Olivia Plateau, Stephan Lautenschlager, Benjamin Steventon, Daniel J. Field

## Abstract

Anseriformes (waterfowl) and Galliformes (landfowl) are among the world’s most recognisable groups of birds, together comprising the clade Galloanserae. Despite their close evolutionary relationship, the skulls of adult anseriforms and galliforms exhibit strikingly distinct morphologies, the developmental basis and evolutionary history of which is poorly understood. To illuminate the developmental and evolutionary underpinnings of cranial disparity between and within these major extant bird clades, we quantitatively investigated ontogenetic changes in cranial morphology across galloanseran phylogenetic diversity, focusing on the previously unexplored post-hatching interval during which adult morphology takes shape. Our results reveal the combined effects of multiple heterochronic shifts early in galloanseran evolutionary history including anseriform hypermorphosis, along with influential non-heterochronic changes leading to substantially more disparate ontogenetic trajectories—and greater cranial variability—in anseriforms than galliforms. Key galloanseran fossils help clarify the polarity of evolutionary shifts in cranial development through galloanseran phylogenetic history and demonstrate that extant galliform cranial morphology is more constrained and retains a more plesiomorphic morphology than that of anseriforms. Our work helps illuminate the developmental basis of the iconic differences in cranial form between waterfowl and landfowl and illustrates the importance of broad phylogenetic and ontogenetic sampling for clarifying patterns of post-hatching developmental divergence among major vertebrate clades.

## Introduction

Galloanserae, the clade uniting Galliformes (landfowl) and Anseriformes (waterfowl), comprises nearly 500 extant species and includes some of the most widely recognisable and economically important birds on Earth such as chickens and ducks^1^. As one of the most deeply diverging clades of extant birds, unravelling macroevolutionary patterns in Galloanserae may have general implications for understanding the early evolutionary history of the avian crown clade, Neornithes. Indeed, the oldest convincing crown bird fossils are frequently interpreted as total-clade representatives of Galloanserae (e.g., *Vegavis*^2–4^ and *Asteriornis*^5^), and even some toothed stem birds exhibit cranial features distinctly reminiscent of extant galliforms and anseriforms, suggesting that galloanserans may provide useful models for interrogating aspects of early neornithine morphology and function^6,7^.

Despite their close phylogenetic affinities^8–12^, many aspects of galliform and anseriform morphology are highly evolutionarily divergent with respect to one-another^13^. These morphological dissimilarities are particularly evident in cranial form: for instance, most Anseriformes are characterised by elongate, flattened bills, deep braincases, and relatively small orbits, whereas Galliformes generally exhibit short, pointed beaks with comparatively shallow braincases and proportionally larger orbits. Partially underlying these differences in craniofacial morphology are evolutionarily divergent aspects of early morphogenesis that have been discerned in chickens, quails, and mallards^14–19^. Such investigations have generally focused on specific aspects of cranial form such as bill shape^17,20,21^, whereas the developmental origins of interclade differences in other aspects of galloanseran cranial morphology, such as the palatal roof and braincase, remain largely unexplored.

To date, explorations of embryological differences in galloanseran cranial development have primarily focused on developmental stages preceding skull ossification^22^, but determining the extent to which these pre-hatching differences account for the evolutionarily divergent cranial forms of adult Anseriformes and Galliformes depends on assessing the degree of similarity in the post-hatching developmental trajectories of these clades. This question has yet to be examined, obscuring our understanding of the potential influence of post-hatching development on macroevolutionary patterns of adult cranial disparity in galloanserans specifically, and among birds more broadly.

Many organisms undergo profound changes in size, shape, physiology, and metabolism during post-embryonic development^23^. For instance, interspecies differences in post-hatching ontogenetic trajectories may result in increases (i.e. ontogenetic divergence^24^), or decreases (ontogenetic convergence^25–26^) in morphological disparity among clades^27^. Furthermore, heterochronic changes (that is, shifts in the rate and timing of developmental events) frequently underlie differences in ontogenetic trajectories and are known to have played key roles in macroevolutionary transitions in a wide range of vertebrate taxa^25,26,28–32^. Examples of the influence of shifts in post-hatching ontogenetic trajectories on vertebrate cranial form include conserved shifts in rostrum length and braincase depth across tetrapods^33^, while investigations in extant birds have usually focused on single species such as chickens^34^, and gulls^35^. With a few exceptions^24^, broad phylogenetic investigations of shifts in post-hatching avian ontogeny have been more limited, and have yet to be undertaken in Galloanserae, limiting our understanding of the mechanistic factors underlying previously discerned ecology-linked differences in cranial morphology^36–38^. To unravel the influence of post-hatching developmental processes on adult cranial disparity in Galloanserae, we used three-dimensional geometric morphometrics and phylogenetic comparative methods to assess ontogenetic patterns of shape and size variation in hatchlings and adults across an unprecedently broad dataset of Anseriformes and Galliformes, extending our data well beyond the three or four species^39^ often used as models in this clade to encompass most major extant galloanseran lineages. We also incorporated cranial reconstructions of pivotal galloanseran fossils into ancestral state estimates and developmental model-informed predictions of fossil hatchling shape to illuminate the timing and polarity of evolutionary changes in craniofacial development across the evolutionary history of Galloanserae.

## Results

### General patterns of shape variation

The morphospace defined by the first two Principal Components of galloanseran skull shape (PCs) accounts for approximately 66% of the total cranial shape variation (Fig.1). Positive scores along PC1 are associated with skulls exhibiting a short and rostrally pointed beak, reduced lacrimals, proportionally large orbits, ventrally facing postorbital processes, and globular, proportionally small braincases. In contrast, negative scores along PC1 are associated with more elongate skulls with proportionally larger, wider, and flatter beaks, more prominent lacrimals, proportionally smaller orbits, cranially facing postorbital processes, and dorsoventrally deeper and laterally narrower braincases. PC2 is associated with more subtle cranial shape differences; negative scores are associated with more pointed and elongate beaks, proportionally wider interorbital regions, proportionally longer postorbital processes, and shallower, longer, and narrower braincases (Fig.1 and Extended Data Figures 1-3). PC1 accounts for over half the total shape variance in our dataset, largely segregating galliforms from anseriforms. PC2 accounts for ∼10% of total shape variance and roughly describes morphological variation between hatchling and adult individuals, although significant ontogenetic change is also evident along PC1, predominantly in Anseriformes.

We found significant differences in mean skull shape between Anseriformes and Galliformes and between hatchlings and adults of the same species (Table S1, see Supplementary Materials and Extended Results for detailed morphological differences). Skulls of adult anseriforms exhibit significantly greater disparity than all other groups considered (Supplementary Figure 1; Tables S2-3), with adults of *Malacorhynchus* and *Spatula* being the most morphologically distinct taxa while the screamer *Chauna* is closer to Galliformes than to other Anseriformes, both at the hatchling and adult stages (Figs.1 & 2a-b, Extended Data Figure 1-3). Conversely, hatchling anseriforms show the lowest degree of morphological disparity—significantly lower than hatchling or adult galliforms (Supplementary Fig.1; Table S2). In contrast to anseriforms, adult and hatchling galliforms exhibit similar levels of disparity, indicating evolutionarily divergent ontogenetic patterns between Galliformes and Anseriformes (Fig.2a; Table S2).

### Ontogenetic convergence/divergence

To compare ontogenetic variation within and among Anseriformes and Galliformes, we calculated angles between the ontogenetic shape trajectories of each species and estimated the degree of ontogenetic divergence^27^ (i.e. when individuals become less similar during ontogeny) versus convergence (i.e. when individuals become more similar during ontogeny) of each pair of species based on pairwise Procrustes distances between immature and adult individuals (see Methods).

Angular comparisons and ontogenetic convergence-divergence results yielded similar overall patterns (Fig. 2c & Extended Data Figure 4). Most ontogenetic trajectories between anseriform and galliform species are ontogenetically divergent with respect to one-another, with the trajectories of the galliforms *Callipepla, Rollulus, Lagopus, Perdix,* and *Coturnix* being the most ontogenetically divergent with respect to anseriforms, and the anseriforms *Cygnus, Callonetta, Anas,* and *Mergus* being the most ontogenetically divergent with respect to galliforms (Fig. 2c & Extended Data Figure 4). Similarly, the greatest angular comparison values were observed between species of Galliformes and Anseriformes (i.e., > 70°), primarily highlighting ontogenetically divergent trajectories (Extended Data Figure 4a).

Within clades, anseriform ontogenetic trajectories tend to converge in shape during ontogeny more strongly than galliforms (Fig. 2c). Relatedly, the greatest angles associated with ontogenetic divergence (i.e., > 70°) are primarily observed in Galliformes, such as in *Callipepla* and *Rollulus*, whereas the greatest angles associated with convergence (i.e., 50° < θ < 60° degrees) are primarily found in Anseriformes, such as in *Oxyura* and *Anas* (Extended Figure 4b). However, most ontogenetic trajectories indicate only slight divergence or convergence within subclades, with many values near zero reflecting primarily parallel slopes (Fig. 2c).

Notably, *Chauna* (Anseriformes: Anhimidae) exhibits an ontogenetic trajectory that is more ontogenetically divergent with respect to other Anseriformes than it is to Galliformes, illustrating that the striking similarities between screamers and galliforms extend beyond morphological similarities to patterns of ontogenetic change.

### Ontogenetic allometry

We found that skull size (i.e. centroid size) variation accounts for ∼31% of shape variation in the total sample (*F*=26.6, p=0.0001). Moreover, regression scores showed some phylogenetic clustering between species-level allometric trajectories in Galliformes and Anseriformes, except in *Chauna* whose trajectory clearly falls within the range of variation of Galliformes (Fig. 3, Supplementary Figure 4). Within Galliformes, allometric trajectories exhibited similar slopes and y-intercepts (shape), but variable lengths broadly associated with body size: *Coturnix* exhibited the shortest trajectory and *Afropavo* exhibited the longest. On the other hand, allometric trajectories in Anseriformes are more variable, with a greater range of y-intercepts, allometric slopes and lengths. The shortest anseriform trajectory was exhibited by *Mergellus* and the longest by *Cygnus* (Fig. 3). Combining the two main axes of shape variation with size showed slight differences between the trajectories of galliforms and anseriforms (Supplementary Figure 5), although these differences are not statistically significant (see below).

Overall, Galliformes and Anseriformes exhibit similar slopes and y-intercepts (Table S4); this shared pattern of allometric ontogenetic change reflects the shift from a skull shape characterised by a relatively short, pointed beak with large orbits and a globular braincase, to a skull shape characterised by a longer bill, proportionally smaller orbits and a less rounded braincase (Fig. 3). Despite these similarities, we found significant shape differences between galliforms and anseriforms at the largest body sizes within our sample (see peramorphosis test in Methods), suggesting that allometric cranial morphology in Anseriformes is peramorphic relative to Galliformes in light of our ancestral reconstructions of galloanseran ontogeny (d=0.29, p =0.000) (Table S4). Furthermore, mean centroid sizes were significantly greater in anseriform hatchlings (d=0.037, p=0.0000) and adults (d=0.15, p=0.0001) relative to Galliformes (Tables S5-6), and the extent of ontogenetic shape (d= 0.08, p=0.002) and size change (d=0.08, p=0.0026) (Tables S5-6) were also significantly greater in Anseriformes than Galliformes. Collectively, these results suggest that hypermorphosis plays a key role in the evolutionary ontogenetic divergence between Galliformes and Anseriformes.

### Detailed variation in ontogenetic variables

The major clades under investigation can be distinguished based on variation in ontogenetic variables (see Methods), with Anseriformes exhibiting considerably greater ontogenetic variability than Galliformes (Fig. 4 & Extended Data Figure 5). Although there is some overlap, interclade separation is largely evident along PC1, with Anseriformes exhibiting greater variance than Galliformes in ontogenetic size and shape change, as well as adult and hatchling skull sizes (Fig.4, Extended Data Figure 5). Within Galliformes, evolutionarily divergent taxa include *Meleagris,* which exhibits particularly large hatchling and adult skull sizes, and a greater degree of ontogenetic shape change than other galliforms (Fig.4). Within Anseriformes, evolutionarily divergent taxa include *Cygnus* and *Anser* whose adult sizes and extent of ontogenetic size change are greater than other anseriforms investigated, and *Anas*, whose hatchling skull size and degree of ontogenetic shape change are greater than other anseriforms.

Ancestral state reconstructions of the five ontogenetic traits investigated affirmed general patterns arising from our ontogenetic morphospace. For instance, crown galliforms tend to be smaller at hatchling and adult sizes, and to exhibit lower degrees of ontogenetic shape and size change than anseriforms (Supplementary Figures 2-3). However, allometric slopes appear similar in both galliforms and anseriforms, with outliers in both clades (Supplementary Figure 4).

Our pairwise ontogenetic morphospace shows that interclade (i.e. galliform-anseriform) differences are more variable than intraclade differences (i.e. comparisons within Galliformes or Anseriformes; Fig.5 & Extended Data Figure 6). Notably, intraclade comparisons in both Anseriformes and Galliformes are dominated by similar ontogenetic traits, as revealed by strong overlap along the major axes of ontogenetic variation in morphospace (Fig. 5 & Extended Data Figure 6). In both groups, major intraclade differences are associated with variation in allometric slopes, and hatchling and adult skull size. Nevertheless, variation among anseriforms is greater than among galliforms.

### Ancestral ontogeny reconstruction

To understand the polarity of evolutionary developmental changes across galloanseran evolutionary history, we estimated the cranial morphology of ancestral hatchlings and adults at key nodes across Galloanserae. First, we estimated skull morphology at the base of the crown clades Galloanserae, Galliformes, Anseriformes and Anatidae using ancestral state reconstruction under the lambda model of evolution, which was identified as the best-fitting model for hatchling-to-adult shape according to the Generalized Information Criterion (GIC) (Table S7). At each node, our results show that estimated hatchling and adult skull shapes are generally similar to one another (Supplementary Figure 7). Notably, ancestral reconstructions for Galloanserae and Galliformes exhibit numerous similarities, as do ancestral reconstructions of Anseriformes and Anatidae (Supplementary Figure 7). Predicted hatchling skull shapes for the fossil taxa *Presbyornis* and *Asteriornis* were similar to their adult counterparts regardless of the specific estimated ancestral ontogenetic trajectory applied to the fossil skull (that is, whether the estimated ancestral ontogenetic trajectories for Galloanserae, Galliformes, Anseriformes or Anatidae were used) (Supplementary Figure 8).

Projections of the reconstructed and predicted skull shapes on the PCA plot of the phylogenetically inclusive extant skull sample (Fig. 6 & Supplementary Figure 7) show that *Asteriornis*, Galliformes, and palaeognaths show relatively similar ontogenetic trajectories separate from representatives of Neoaves and most anseriforms, including the fossil total-clade anseriform *Presbyornis*. Notably, the screamer *Chauna* represents the only anseriform in our sample clustering with Galliformes. Hatchling and adult ancestral reconstructions for each investigated clade plotted between Anseriformes and the *Asteriornis*-Galliformes-Palaeognathae cluster, with ancestral Anseriformes and ancestral Anatidae plotting closer to extant Anseriformes, and ancestral Galloanserae and ancestral Galliformes plotting closer to extant Galliformes (Fig. 6 & Supplementary Figure 7).

The shape of adult *Presbyornis* plots near extant Anatidae, and adult *Asteriornis* plots near adult *Chauna.* The same pattern is observed for predicted fossil hatchling shapes, with all predicted *Presbyornis* hatchling shapes plotting within extant Anatidae, and all predicted *Asteriornis* hatchling shapes plotting closer to extant Galliformes. Altogether, these results are congruent with a galliform-like ontogeny having been ancestral for Galloanserae, with morphologically influential evolutionary changes in ontogenetic trajectories evolving within total-clade Anseriformes (Fig. 6 & Supplementary Figure 7).

## Discussion

### The evolution of cranial form in Galloanserae

Our results illustrate strong phylogenetic clustering of hatchling skull shapes, with galliforms and non-anhimid anseriforms generally occupying distinct regions of morphospace, illustrating the influence of pre-hatching development on generating evolutionarily divergent cranial forms in these clades at an early point in ontogeny^39^ (Fig. 1; Extended Data Figure 8). However, our results also reveal the important extent to which interclade shape differences increase from hatching to the adult stage, illustrating the previously unexplored role of divergent post-hatching ontogenetic trajectories in driving the development of adult cranial form disparity between Galliformes and Anseriformes (Fig. 2; Extended Data Figure 4; Table S1). Collectively, our results confirm that cranial ontogenetic trajectories and resultant adult morphologies are substantially more constrained in Galliformes than in Anseriformes^40^, which may contribute to previous observations highlighting the notable evolvability of the anseriform skull in response to shifts in diet^41,42^.

The comparatively low cranial disparity of galliforms suggests limited within-clade variance in ontogenetic trajectories, apart from weak phylogenetic clustering of phasianid hatchlings relative to other Galliformes (Fig. 2a), which requires further investigation. By contrast, the markedly greater cranial disparity in Anseriformes relative to Galliformes (Fig. 1, Supplementary figure 1) emphasises the comparative plasticity of anseriform post-hatching ontogenetic patterns. This enhanced morphological variability is exemplified by anseriform subclades with highly distinct morphologies, ranging from the comparatively galliform-like skulls of screamers (Anhimidae) to the narrow, elongate skulls of mergansers (Mergini), to taxa with strongly spatulate bills such as *Spatula* and *Malacorhynchus.* Distinctive aspects of the cranial morphologies of these species seem to first arise during embryonic development, indicated by the notable disparity observed among anseriform hatchlings (Figs. 1b & 2a), and is further exaggerated during post-hatching development (Figs. 1b & 2b).

The relative proximity of the inferred ancestral skull shapes for Galliformes and Galloanserae to extant Galliformes, as opposed to extant Anseriformes, suggest that extant galliform morphology is more reflective of the plesiomorphic galloanseran condition than that of Anseriformes (Fig. 6 & Supplementary Figure 7). Moreover, the proximity of *Asteriornis* (the earliest known clear representative of total-group Galloanserae) and *Chauna* (representing Anhimidae, the sister group to all other Anseriformes) to crown-galliform cranial morphospace further supports the interpretation of galliform skull morphology as being generally plesiomorphically retentive (Fig. 6 & Supplementary Figure 7). As such, post-hatching ontogenetic trends in galliform cranial morphonology are likely less derived than those of anseriforms; indeed, most crown galliforms are similar to the fossil galloanserans included in our dataset in estimated post-hatching size and shape changes (Supplementary Figures 2-3).

In most discernible respects, the cranial morphology of screamers (Anseriformes: Anhimidae, represented in our dataset by *Chauna*) appears to be effectively intermediate between that of Galliformes and Anseriformes, congruent with previous hypotheses that the galliform-like beak of screamers reflects the retention of a plesiomorphic galloanseran feature^43^. By contrast, the relative size of the screamer skull, and its degree of ontogenetic size and shape change throughout ontogeny exceed those of Galliformes and are instead more comparable to those of other Anseriformes (Supplementary Figures 2-3). Some palaeontological investigations have interpreted the comparatively galliform-like skulls of screamers to reflect a secondary reversal from a more typical anseriform ancestral condition (for instance, one resembling the Paleocene total-clade anseriform *Conflicto*^44^). This interpretation is supported by some phylogenetic investigations supporting stem-anseriform positions for the fossil taxa *Presbyornis* and *Conflicto*, which both exhibit skulls comparable to those of extant anatoids, although conflicting phylogenetic results emphasise the lability of this hypothesis^3^, and the importance of ongoing work to clarify phylogenetic relationships among fossil total-clade anseriforms^45,46^. We favour the interpretation of *Conflicto* and *Presbyornis* as stem-group anatoids^3^, supporting a comparatively simple macroevolutionary scenario in which the evolutionary divergence of the distinctive anatoid skull from galliform-like precursors may have arose in two plausible steps: the skulls of the earliest crown anseriforms would have been more similar to those of extant galliforms than anatoids (supported by the proximity of *Chauna* to crown Galliformes in morphospace; Fig. 1), although they may have exhibited a greater degree of post-hatching ontogenetic shape change than Galliformes (Supplementary Figure 2). Subsequently, increased embryonic and post-hatching developmental divergence occurred along the anatoid stem lineage characterised by the acquisition of a long and comparatively flat bill, smaller orbits, and a shorter and narrower braincase. The relative order by which these two heterochronic changes evolved is uncertain, but this scenario would imply that the Eocene fossil anseriform *Presbyornis* (and by inference, the Paleocene fossil anseriform *Conflicto*—unsampled here due to incomplete preservation), capture a relatively early stage in total-clade anatoid history subsequent to the evolution of these heterochronic shifts, as evidenced by the recovery of *Presbyornis* within the Anatidae cluster (Fig. 6 & Supplementary Figure 7).

### Heterochronic drivers of shifts in galloanseran cranial development

Though our ability to assess intraclade variability in ontogenetic trajectories is limited by our sample size per species, our results document variable ontogenetic shape trajectories (Fig. 1), degrees of ontogenetic shape and size change (Supplementary Figure 2), and hatchling and adult skull size (Fig. 4,5, and Supplementary Figure 3; Table S6) between galliforms and anseriforms, providing insight into the factors contributing to the evolution and development of galloanseran cranial form. Our statistical tests enabled us to evaluate the influence of alternative heterochronic mechanisms as drivers of differences between galliform and anseriform cranial ontogeny (Table S4). Both ontogenetic shape and size change are significantly greater in Anseriformes than Galliformes (Δsize = 0.08, Δshape = 0.03) (Table S5), though their ontogenetic allometric vectors overlap (Table S4). These results suggest that differences in cranial ontogenetic trajectories in Anseriformes and Galliformes are at least partially ascribable to peramorphic hypermorphosis in anseriforms relative to the ancestral condition for Galloanserae (Table S4, Extended Data Figure 7). Additionally, the skull shapes of anseriform and galliform hatchlings are already statistically different at hatching (Extended Data Figure 7), suggesting a mechanism of pre-hatching ontogenetic divergence that we dub an ‘ontogenetic shift’, characterised by shifted onsets and offsets of allometric vectors that otherwise overlap in their slope, direction, and intercept (Extended Data Figure 7). Our tests illustrate that alternative heterochronic mechanisms have limited explanatory power applied to our data (Extended Data Figure 7; Table S4), enabling us to conclude that that interclade differences in anseriform and galliform ontogenetic size and shape variation are predominantly the result of the combined effects of anseriform hypermorphosis and an ontogenetic shift (Extended Data Figure 9). These heterochronic differences may underpin the acquisition of evolutionarily divergent cranial forms in anseriforms relative to the comparatively galliform-like plesiomorphic condition for Galloanserae (Extended Data Figure 9).

The ontogenetic shift we identify may be related to differences in embryonic cranial morphogenesis and incubation period between anseriforms and galliforms. Investigations of interspecific embryonic morphogenetic differences in the bills of mallards, chickens and quails have identified numerous morphogenetic differences that might account for the comparatively long bills of mallards. For instance, ducks exhibit prolonged proliferation and slower maturation of cranial neural crest cells (CNCCs)—the embryonic precursors to the beak—relative to chickens and quails^17,18,47^. Moreover, duck embryos exhibit a greater number and faster rate of proliferation of mandibular CNCCs relative to quail embryos, resulting in the comparatively long mandibles of ducks^18^, while the quail mandible exhibits a greater amount of bone resorption at later stages of craniofacial formation, further contributing to the distinctively shorter mandibles of galliforms relative to anseriforms^48^. In addition to these factors, Anseriformes tend to exhibit longer incubation periods, larger hatchling sizes^49^, and longer lifespans^50^ than Galliformes, which may contribute both to their relatively larger skulls and their comparatively hypermorphic growth.

To the extent enabled by our sampling, our results also reveal some insight into plausible heterochronic shifts driving cranial divergence within various galloanseran subclades. For instance, representatives of the anseriform subclade Mergini are characterised by elongate, narrow bills associated with their piscivorous diet^51^. The amount of ontogenetic shape and size change is greater in *Mergus* than in *Mergellus* (Supplementary Figures 2-3), but their hatchling shapes are similar. The similarity in the origins and angles of the shape vectors of these two taxa, and the shorter shape change for *Mergellus* (Fig. 1), indicate that the skull of *Mergellus* is likely paedomorphic relative to *Mergus*, further supported by the small size of *Mergellus* relative to other members of Mergini^51^ (see Supplementary Results for further discussion of intraclade heterochronic patterns) although this hypothesis remains to be tested by future research.

At various phylogenetic scales, the present study thus provides insight into the important extent to which shifts in post-hatching ontogenetic trajectories have influenced macroevolutionary changes in cranial form across Galloanserae—a clade providing an ideal case study for unravelling the evolution of divergent cranial morphologies among avian sister taxa.

## Online Materials & Methods

### Specimen acquisition and shape quantification

Two datasets were assembled for this study. The first comprised sixty-one hatchling-stage and adult specimens with intact skulls spanning all major extant galloanseran subclades, except Anseranatidae (see Extended Data Figures 1-3; Table S8 for the full list). The second comprised eleven hatchling-stage and adult specimens including incomplete specimens of the Magpie Goose (*Anseranas semipalmata;* Anseranatidae), two palaeognaths (the Common Ostrich *Struthio camelus* and the Tataupa Tinamou *Crypturellus tataupa*), and two neoavians (the Feral Rock Pigeon, *Columba livia* and the Ruff, *Calidris pugnax*). We also included the reconstructed fossil skull of the probable stem galloanseran *Asteriornis maastrichtensis*^5^ (see below), along with the stem anseriform *Presbyornis pervetus*^13^ (Table S8). Most specimens were scanned with a Nikon XTEK H 225 ST µCT scanner at the Cambridge Biotomography Centre using scanning parameters maximising bone contrast (voltage: 90–170 kV, exposure time: 354–500 ms, current: 50–225 μA), while others were scanned at the following facilities: The Imaging and Analysis Centre at the Natural History Museum (London), the Centre for Advanced Imaging (The University of Queensland), the School of Earth Sciences X-ray Tomography Facility (University of Bristol), The University of Michigan Museum of Zoology MicroCT Scanning Laboratory, Ohio University MicroCT Facility (Ohio University), and the Cold Spring Harbor Laboratory. CT scan data were digitally segmented using Dragonfly 2024.1 (Object Research Systems, Inc, Montreal, Quebec, Canada). Segmented three-dimensional skull meshes from the first dataset were used for most of our analyses, while the second dataset was used for our ancestral shape reconstructions to ascertain the phylogenetic polarity of developmental changes.

Subsequently, we used three-dimensional geometric morphometrics to quantify the shapes of all the skull 3D meshes. The shapes of the skull meshes from the first dataset, made up exclusively of Galloanserae specimens, were landmarked with forty-nine fixed landmarks and 272 semi-landmarks, covering the shape of the beak, palate, orbit, and the occipital region (Tables S9-11, Extended Data Figure 10). The landmarking scheme was applied to adults and near-hatching skull meshes using 3D Slicer^52^ and applying established homology criteria (Tables S11). Subsequently, the landmarking scheme was eroded to a reduced configuration of fourteen fixed landmarks and 86 semi-landmarks covering the beak and orbit regions (Tables S9-10) to enable the inclusion of incomplete fossil and some extant specimens in our analyses. This reduced configuration was applied to both datasets. Raw landmarks from both datasets were exported to the R statistical environment v.4.3.3^53^, where all subsequent analyses were conducted.

### Statistical analyses

#### Shape disparity

Shape data (Procrustes residuals) were extracted using a Generalised Procrustes Analysis using the *gpagen* function from the geomorph package (4.0.8)^53–57^. To explore evolutionary and ontogenetic variation within our taxon sample we conducted a Principal Components Analysis (PCA) using all shape data and the *gm.prcomp* function from geomorph, package and plotted the first three Principal Components. Subsequently, all specimens were divided into four groups: galliform hatchlings, galliform adults, anseriform hatchlings, and anseriform adults. Differences in shape variation among the four groups was quantified using Procrustes ANOVA and pairwise tests using the RRPP package (2.1.0)^55,57^. Differences between the mean shape for each group were determined using the *procD.lm* and the *pairwise* functions from geomorph (4.0.8). Morphological disparity in each group was subsequently quantified by calculating the Procrustes variance using the *morphol.disparity* function from geomorph (4.0.8). To visualise shape differences among hatchlings and among adults we calculated the Procrustes distance between each pair of hatchlings and each pair of adults using the *dist* function and plotted them as heatmaps using the *heatmap* function in the stats package (4.4.2).

#### Ontogenetic convergence/divergence analysis

We quantified the pattern and strength of ontogenetic convergence, or divergence, across all pairs of species. Convergence/divergence of ontogenetic trajectories was assessed in two ways. First, we used a distance-based method^27^, that represents the ratio of the calculated Procrustes distances between the hatchlings of two species and the adults of those same species. Convergence is indexed as a decrease in Procrustes distance from hatchlings to adults, and *vice versa* for ontogenetic divergence. We also assessed the angular distance between ontogenetic trajectory vectors for each species pair^28,58^. To distinguish between large absolute angular values due to convergence or divergence, we used the results of the distance-based method to assign a positive sign to the angle of convergent trajectories and a negative sign to the angle of divergent ones.

#### Allometric ontogenetic analysis

We used the *procD.lm* function to explore the relationship between shape and size in our sample and plotted the regression scores of shape against cranial centroid size to visualize allometric ontogenetic changes in the full sample. We further visualised the relationship between shape and size change by plotting shape using principal components (PC1 and PC2) and centroid sizes in a 3D plot.

We evaluated evolutionary changes in ontogenetic trajectories based on^59^, which has been widely used in many phylogenetic groups including crocodilians^60^, squamates^26,30^, birds^24^, non-avian dinosaurs^61^ and mammals^62^. This approach relies on evaluating the relationship between shape and size changes throughout ontogeny across different groups. Differences in these relationships across related taxa therefore provide evidence for, and allow us to distinguish among, different heterochronic and non-heterochronic mechanisms underlying ontogenetic differences throughout the evolutionary history of a clade. Here, we applied this approach to the differences in ontogenetic change between Galliformes and Anseriformes, and Phasianidae and Anatidae^59^ (Extended Data Figure 7). First, we applied the homogeneity of slopes test (HOS)^30,63,64^, comparing differences in the direction of shape change relative to size (that is, slope vector angle), and the amount of shape change per unit of size change (that is, slope vector length). This is equivalent to assessing the rate of shape/size change, and enables testing for the acceleration of ontogenetic change between two clades^65^. Second, for the comparisons where allometric slopes were statistically indistinguishable, we used the intercept test^30,60^ to determine whether or not allometric trajectories overlap or not, indicating a common allometric vector, and therefore, a common allometric pattern among groups. Third, for the comparisons where intercepts were statistically indistinguishable, we applied the peramorphosis test^26,30,60^ to determine whether overlapping allometric trajectories have a similar endpoint, by determining whether shapes at maximum size are statistically different between groups. Finally, for the comparisons where shape differences were found at the adult stage, we compared statistical differences in hatchling shape and sizes among clades using *procD.lm* and *pairwise.* This final step was aimed to distinguish between two different but interrelated heterochronic mechanisms^65^. Namely hypermorphosis, in which two ontogenetic trajectories begin at similar points, but the ontogeny of one extends further in shape and size than the other^65^; and ‘ontogenetic shift’, in which the total change in both ontogenetic trajectories is the same, but the onset and end of ontogenetic growth is shifted. Statistical significance in all our tests was evaluated using 10,000 iterations.

#### Variation in ontogenetic traits

Despite our datasets being unprecedently comprehensive in encompassing all major lineages of Galloanserae, each species is only represented by one hatchling and one adult individual, precluding species-level statistical testing. To overcome this limitation and gain further insight into species-level ontogenetic evolution in Galloanserae, we extracted five ontogenetic variables from each species-level ontogeny: i) ontogenetic shape change, ii) ontogenetic size change, iii) adult skull size, iv) hatchling skull size, and v) the rate of shape to size change. We then reconstructed the evolutionary history of each of these traits using maximum likelihood with the R function fastAnc and plotted the estimated values across the phylogeny of Galloanserae using contMap from the R package phytools v.2.3.0^66^. We summarised these values in an ontogenetic-phylomorphospace by standardising each of the aforementioned variables to values between 0 and 1 to avoid over-weighting of some variables over others, and summarised the species-values dispersion using a Principal Component Analyses over which we projected our phylogeny. We also included biplots showing the weights of original variables in each Principal Component axis; doing so allowed us to visually inspect whether clades and species differ in general ontogenetic characteristics and which ontogenetic traits determine these differences^67^.

To further explore how inter- and intraclade ontogenetic differences vary, we compared pairwise differences among all pairs of species-level ontogenies in eight ontogenetic variables : i) angle of shape change, ii) degree of shape convergence/divergence (Procrustes differences between hatchling shapes – Procrustes differences between adult shapes), iii) differences in hatchling shape, iv) differences in hatchling size, v) differences in adult shape, vi) differences in adult size, vii) differences in rate of shape to size change (allometry), and viii) angle of allometric change. We summarised these pairwise comparisons by means of heatmaps for shape convergence/divergence and ontogenetic shape angles. We then summarised all eight variables in a Pairwise-Ontogenetic-Morphospace using a Principal Component Analysis using the function *prcomp* from the R package stats v. 4.3.3 including biplots showing the weights of individual variables in each Principal Component axis. This allowed us to visually inspect whether ontogenetic comparisons among species differ between and within Galliformes and Anseriformes, and which ontogenetic traits are driving these differences^66,67^.

#### Cranial reconstruction of the early galloanseran Asteriornis maastrichtensis

For the restoration of *Asteriornis* the segmented skull elements were imported into Blender 2.81 to remove taphonomic artefacts, articulate individual elements, and perform retrodeformation steps. This process generally followed the guidelines outlined in^68^ and are described in detail below.

Individual elements were repositioned using the translation and rotation tools in Blender. Smaller breaks and fractures were removed using the Sculpt mode in Blender using the Clay, Inflate, and Smooth tools. The left and right premaxilla were selected as the starting point for the restoration as these elements are largely complete and provided a clear reference for the skull anatomy. The left premaxilla is more completely preserved, including larger portions of the premaxillary body and the maxillary process. The latter was moved dorsally to allow an articulation with the maxilla. Next, the left nasal was articulated. For this, the premaxillary process had to be rotated as the part had been deformed. Further distortion in the main body of the nasal was corrected so that it formed a flush surface with the premaxilla. The right nasal showed more deformation and was only used to inform the general outline of the element. The frontals and partially preserved postorbitals are largely articulated and retain a clear articulation facet with the nasals which allowed them to be articulated with the latter. The mesethmoid is relatively complete but the ventral portion had been distorted towards the left side. This was straightened using a Lattice Deform modifier in Blender. These elements provided a general framework for the restoration of the other elements for which the preservation or position were more ambiguous.

Of the two maxillae, only the right was preserved well enough to be used in the restoration. It was mirrored to the opposing side and articulated with the premaxilla. The laterosphenoid region is incompletely preserved on both sides but a more complete element could be produced by mirroring and superimposing the right onto the left element. Together with the postorbital, the posterior curvature of the orbital region could be restored. Although only fragmentarily preserved, the basisphenoid retains general details about the element’s size and outline and could be fitted to the laterosphenoid. The left quadrate is among the best-preserved elements, but its placement was less clear. It was tentatively articulated first ventral to the postorbital/laterosphenoid region with its final position subsequently being informed by the spatial arrangement of the mandible. The jugals are only fragmentarily preserved and large parts of the element had to be reconstructed by extending the existing morphology using the Lattice Deform modifier.

As with the skull, the mandibular elements are better preserved on the left side, particularly the symphyseal region of the dentary. The left element was therefore used for the main part of the reconstruction with the right counterpart filling in gaps. As the dentaries had been deformed considerably, they were straightened using the Lattice Deform modifier. The position and orientation were informed by the overall dimensions (i.e. skull width) of the other elements. The left splenial is incompletely preserved but is largely undeformed and further provided guidance for the restoration of the dentary. A further mandibular element originally identified as retroarticular process instead represents the medial process of the mandible^69^.

Once all elements had been re-articulated for the better-preserved left side, they were mirrored to generate a completely symmetrical skull model. Missing elements were created to fill in the gaps by using a box-modelling approach^70^. These included parts of the laterosphenoid-mesethmoid-basisphenoid region, the distal part of the postorbital, parts of the palate, the postdentary region of the mandible, and the entire back of the skull. While most of these regions could be inferred by extending the morphology between the preserved elements, the palate had to be reconstructed in its entirety and was modelled as two simple rod-like structures seen in the early anseriform *Presbyornis pervetus*, but also Galliformes in general. Similarly, the back of the skull is unknown, and the preserved elements provide no information about its size and dimensions. As a stand-in, the respective skull region of *Meleagris gallopavo* was scaled and fitted to the remaining skull.

In a final step, minor gaps between the individual elements were closed using the Grab tool in Sculpt mode. Subsequently, all elements were remeshed using the Remesh modifier to create solid surface models. High resolution settings were used to preserve all necessary detail while removing minor artefacts.

#### Ancestral ontogeny reconstruction

Although our previous clade-level and species-level analyses allowed us to characterise evolutionary ontogenetic changes between and within crown Galliformes and crown Anseriformes, they could not directly inform on the polarity of these ontogenetic changes. That is, whether ontogenetic trajectories in crown Anseriformes are derived and ontogenetic trajectories in crown Galliformes are more similar to the ancestral condition for Galloanserae, or vice versa. This determination is important to shed light on the underlying mechanisms behind key differences in ontogenetic trajectories between the two clades (e.g., distinguishing between postdisplacement or predisplacement).

To that end, we used the reduced landmark configurations of the specimens from the two datasets and extracted shape data using generalised GPA with the *gpagen* function from geomorph (4.0.8). A Principal Component Analysis was carried out, and PC1-3 plots were generated to visualise the main patterns of cranial disparity across the more phylogenetically inclusive sample. Then, re reconstructed the ancestral adult and hatchling skull shapes for the major galloanseraen subclades. First, we fit four evolutionary models (Brownian motion, Ornstein-Uhlenbeck, Early Burst and the lambda model) to the shape data across galloanseran phylogeny using the function *mvgls* from the mvMORPH package (1.2.1)^71^. We then used the Generalised Information Criterion (GIC) to determine the relative fit of each model to the shape data, using the *gic* function. The model with the best fit was then used to predict ancestral adult and hatchling skull shapes for four clades (Galloanserae, Galliformes, Anseriformes, and Anatidae) using the *ancestral* function from the mvMORPH package (1.2.1)^71^, and these reconstructed shapes were then projected onto the aforementioned PCA plots.

Subsequently, we calculated shape differences between reconstructions of the ancestral adult and hatchling shapes. These differences were applied to the landmark configurations of adult *Asteriornis* and *Presbyornis* skulls to predict four multiple alternative shapes for their hatchling skulls. Estimated ontogenetic trajectories at the Galloanserae and Galliformes nodes were applied to *Asteriornis* to produce two possible reconstructions of its hatchling skull geometry. For *Presbyornis*, reconstructed ontogenetic trajectories at the Galloanserae, Anseriformes, and Anatidae nodes were used to produce three possible reconstructions of hatchling skull shape. We then projected all five predicted hatchling shapes onto the more phylogenetically inclusive PCA plot.

## Supporting information

Code used for the analysis

## Data Availability

All meshes were uploaded to morphosource database ( https://www.morphosource.org/projects/000719441/temporary_link/Q8BkFHFjDhM9S7QypkSacnGf?locale=en)

## Code Availability

The code used for this manuscript is provided in the supplementary materials.

## Acknowledgments

We thank Drs. Guillermo Serrano Nájera, Dillan Saunders, Alejandra Guzman Herrera, Apolline Delahaye, other members of the Steventon group, and Dr. Albert Chen for help in result interpretation and visualisations. We also thank Keturah Smithson for CT scanning at the Cambridge Biotomography Centre. We thank Nick Willis (Anglia Wildfowl farm) and Judith White (Natural History Museum, Tring) for providing access to adult and hatchling specimens. We thank Eloise Hunt (Natural History Museum, London), Heather Janetzki (Queensland Museum), Ekaterina Strounina (University of Queensland), Patrick O’Connor and Joseph Groenke (Ohio University) for providing CT scans of several specimens. This work was funded by UKRI grant MR/X015130/1 to DJF. For the purpose of open access, the authors have applied a Creative Commons Attribution (CC BY) licence to any Author Accepted Manuscript version arising.

## References

1 Billerman, S. M., Keeney, B. K., Rodewald, P. G. & Schulenberg, T. S. (Cornell Laboratory of Ornithology, Ithaca, NY, USA, 2022).

2 Clarke, J. A., Tambussi, C. P., Noriega, J. I., Erickson, G. M. & Ketcham, R. A. Definitive fossil evidence for the extant avian radiation in the Cretaceous. Nature 433, 305–308 (2005). 10.1038/nature03150

3 Torres, C. R. et al. Cretaceous Antarctic bird skull elucidates early avian ecological diversity. Nature 638, 146–151 (2025). 10.1038/s41586-024-08390-0

4 Irazoqui, F., Acosta Hospitaleche, C., Paulina-Carabajal, A., Bona, P. & Vega, N. New Species of Vegavis (Neornithes) from Antarctica Highlights Unexpected Cretaceous Antarctic Diversity. Diversity 18, 82 (2026).

5 Field, D. J., Benito, J., Chen, A., Jagt, J. W. M. & Ksepka, D. T. Late Cretaceous neornithine from Europe illuminates the origins of crown birds. Nature 579, 397–401 (2020). 10.1038/s41586-020-2096-0

6 Benito, J., Kuo, P.-C., Widrig, K. E., Jagt, J. W. M. & Field, D. J. Cretaceous ornithurine supports a neognathous crown bird ancestor. Nature 612, 100–105 (2022). 10.1038/s41586-022-05445-y

7 Benito, J. et al. Shouldering the challenge of deciphering avian palate evolution. Proc Natl Acad Sci U S A 123, e2514111123 (2026). 10.1073/pnas.2514111123

8 Prum, R. O. et al. A comprehensive phylogeny of birds (Aves) using targeted next-generation DNA sequencing. Nature 526, 569–573 (2015). 10.1038/nature15697

9 Hackett, S. J. et al. A phylogenomic study of birds reveals their evolutionary history. Science 320, 1763–1768 (2008). 10.1126/science.1157704

10 Jarvis, E. D. et al. Whole-genome analyses resolve early branches in the tree of life of modern birds. Science 346, 1320–1331 (2014). 10.1126/science.1253451

11 Stiller, J. et al. Complexity of avian evolution revealed by family-level genomes. Nature 629, 851–860 (2024). 10.1038/s41586-024-07323-1

12 Chen, A., Benito, J. & Field, D. J. Phylogenetic nomenclature of major clades in total-group Galloanserae (Aves), with a name for the clade uniting Cracidae and Phasianoidea. Bulletin of Phylogenetic Nomenclature (In review).

13 Olson, S. L. & Feduccia, A. Presbyornis and the Origin of the Anseriformes (Aves: Charadriomorphae). Smithsonian Contributions to Zoology, 1–24 (1980). 10.5479/si.00810282.323

14 Brugmann, S. A. et al. Comparative gene expression analysis of avian embryonic facial structures reveals new candidates for human craniofacial disorders. Hum Mol Genet 19, 920–930 (2010). 10.1093/hmg/ddp559

15 Smith, F. J. et al. Divergence of craniofacial developmental trajectories among avian embryos. Dev Dyn 244, 1158–1167 (2015). 10.1002/dvdy.24262

16 Schneider, R. A. Neural crest and the origin of species-specific pattern. Genesis 56, e23219 (2018). 10.1002/dvg.23219

17 Wu, P., Jiang, T. X., Shen, J. Y., Widelitz, R. B. & Chuong, C. M. Morphoregulation of avian beaks: comparative mapping of growth zone activities and morphological evolution. Dev Dyn 235, 1400–1412 (2006). 10.1002/dvdy.20825

18 Fish, J. L., Sklar, R. S., Woronowicz, K. C. & Schneider, R. A. Multiple developmental mechanisms regulate species-specific jaw size. Development 141, 674–684 (2014). 10.1242/dev.100107

19 Young, N. M. et al. Embryonic bauplans and the developmental origins of facial diversity and constraint. Development 141, 1059–1063 (2014). 10.1242/dev.099994

20 Schneider, R. A. Regulation of Jaw Length During Development, Disease, and Evolution. Curr Top Dev Biol 115, 271–298 (2015). 10.1016/bs.ctdb.2015.08.002

21 Jheon, A. H. & Schneider, R. A. The cells that fill the bill: neural crest and the evolution of craniofacial development. J Dent Res 88, 12–21 (2009). 10.1177/0022034508327757

22 . (!!! INVALID CITATION !!! 21,22).

23 Ricklefs, R. E. & Starck, J. M. in Avian Growth and Development (eds R.E. Ricklefs & J. M. Starck) 31–58 (Oxford University Press, 1998).

24 Navalon, G. et al. Craniofacial development illuminates the evolution of nightbirds (Strisores). Proc Biol Sci 288, 20210181 (2021). 10.1098/rspb.2021.0181

25 Morris, Z. S., Vliet, K. A., Abzhanov, A. & Pierce, S. E. Heterochronic shifts and conserved embryonic shape underlie crocodylian craniofacial disparity and convergence. Proceedings of the Royal Society B: Biological Sciences 286, 20182389 (2019). 10.1098/rspb.2018.2389

26 Esquerre, D., Sherratt, E. & Keogh, J. S. Evolution of extreme ontogenetic allometric diversity and heterochrony in pythons, a clade of giant and dwarf snakes. Evolution 71, 2829–2844 (2017). 10.1111/evo.13382

27 Plateau, O., Navalon, G., Benito, J. & Field, D. J. Developmental underpinnings of morphological disparity in the avian bony palate. Nat Commun 17, 3806 (2026). 10.1038/s41467-026-69576-w

28 Foth, C., Hedrick, B. P. & Ezcurra, M. D. Cranial ontogenetic variation in early saurischians and the role of heterochrony in the diversification of predatory dinosaurs. PeerJ 4, e1589 (2016). 10.7717/peerj.1589

29 Bhullar, B. A. et al. Birds have paedomorphic dinosaur skulls. Nature 487, 223–226 (2012). 10.1038/nature11146

30 Ollonen, J. et al. Dynamic evolutionary interplay between ontogenetic skull patterning and whole-head integration. Nat Ecol Evol (2024). 10.1038/s41559-023-02295-3

31 Wilson, L. A. B. et al. Patterns of ontogenetic evolution across extant marsupials reflect different allometric pathways to ecomorphological diversity. Nat Commun 14, 2689 (2023). 10.1038/s41467-023-38365-0

32 Senevirathne, G. et al. The evolution of hominin bipedalism in two steps. Nature 645, 952–963 (2025). 10.1038/s41586-025-09399-9

33 Emerson, S. B. & Bramble, D. M. in The skull: functional and evolutionary mechanisms Vol. 3 (eds J. Hanken & B.K. Hall) 384–421 (1993).

34 Marugán-Lobón, J., Blanco Miranda, D., Chamero, B. & Martín Abad, H. On the importance of examining the relationship between shape data and biologically meaningful variables. An example studying allometry with geometric morphometrics. Spanish Journal of Palaeontology 28, 139–148 (2020). 10.7203/sjp.28.2.17848

35 Hanai, T., Iwami, Y., Tomita, N. & Tsuihiji, T. Postnatal cranial ontogeny and growth strategies in the black-tailed gull Larus crassirostris breeding on Kabu Island, Aomori, Japan. Journal of Zoology 315, 183–198 (2021). 10.1111/jzo.12907

36 Felice, R. N., Tobias, J. A., Pigot, A. L. & Goswami, A. Dietary niche and the evolution of cranial morphology in birds. Proc Biol Sci 286, 20182677 (2019). 10.1098/rspb.2018.2677

37 Hunt, E. S. E., Felice, R. N., Tobias, J. A. & Goswami, A. Ecological and life-history drivers of avian skull evolution. Evolution 77, 1720–1729 (2023). 10.1093/evolut/qpad079

38 Chatterji, R. M., Williams, S. & Buckner, J. C. Geometric morphometrics enables accurate predictions of paleoecology and reveals unique adaptations to an expanded niche space in extinct waterfowl. bioRxiv, 2025.2005.2029.656821 (2025). 10.1101/2025.05.29.656821

39 Arnaout, B., Brzezinski, K., Chen, A., Steventon, B. & Field, D. J. Galloanseran cranial development highlights exceptions to von Baer’s laws. EvoDevo 16, 17 (2025). 10.1186/s13227-025-00253-7

40 Linde-Medina, M. Testing the cranial evolutionary allometric ‘rule’ in Galliformes. J Evol Biol 29, 1873–1878 (2016). 10.1111/jeb.12918

41 Chatterji, R. et al. Dietary specialization drives adaptation, convergence, and integration across the cranial and appendicular skeleton in Waterfowl (Anseriformes). bioRxiv, 2024.2009.2021.614171 (2024). 10.1101/2024.09.21.614171

42 Olsen, A. M. Feeding ecology is the primary driver of beak shape diversification in waterfowl. Functional Ecology 31, 1985–1995 (2017). 10.1111/1365-2435.12890

43 Zelenkov, N. V. Stable morphological types and mosaicism in the macroevolution of birds (Neornithes). Biology Bulletin Reviews 6, 208–218 (2016). 10.1134/s2079086416030087

44 Tambussi, C. P., Degrange, F. J., De Mendoza, R. S., Sferco, E. & Santillana, S. A stem anseriform from the early Palaeocene of Antarctica provides new key evidence in the early evolution of waterfowl. Zoological Journal of the Linnean Society 186, 673–700 (2019). 10.1093/zoolinnean/zly085

45 Mayr, G., Carrió, V. & Kitchener, A. C. On the "screamer-like" birds from the British London Clay: An archaic anseriform-galliform mosaic and a non-galloanserine "barb-necked" species of. Palaeontol Electron (2023). 10.26879/1301

46 Field, D. J. Paleontology: Ducks all the way down? Current Biology 35, R409–R412 (2025). 10.1016/j.cub.2025.03.051

47 Wu, P., Jiang, T. X., Suksaweang, S., Widelitz, R. B. & Chuong, C. M. Molecular shaping of the beak. Science 305, 1465–1466 (2004). 10.1126/science.1098109

48 Ealba, E. L. et al. Neural crest-mediated bone resorption is a determinant of species-specific jaw length. Dev Biol 408, 151–163 (2015). 10.1016/j.ydbio.2015.10.001

49 Laird, A. K. Dynamics of embryonic growth. Growth 30, 263–275 (1966).

50 Prinzinger, R. Life-Span in Birds and the Aging Theory of Absolute Metabolic Scope. Comp Biochem Phys A 105, 609–615 (1993). Doi 10.1016/0300-9629(93)90260-B

51 Todd, F. S. Natural History of Waterfowl. (Hancock House Publishers, 1996).

52 Fedorov, A. et al. 3D Slicer as an image computing platform for the Quantitative Imaging Network. Magn Reson Imaging 30, 1323–1341 (2012). 10.1016/j.mri.2012.05.001

53 R: A Language and Environment for Statistical Computing (R Foundation for Statistical Computing, Vienna, Austria, 2025).

54 Adams, D. C., Otárola-Castillo, E. & Paradis, E. geomorph: anrpackage for the collection and analysis of geometric morphometric shape data. Methods in Ecology and Evolution 4, 393–399 (2013). 10.1111/2041-210x.12035

55 Collyer, M. L. & Adams, D. C. RRPP: linear model evaluation with randomized residuals in a permutation procedure. R package version 0.4. 0 (2019).

56 Baken, E. K., Collyer, M. L., Kaliontzopoulou, A. & Adams, D. C. geomorph v4.0 and gmShiny: Enhanced analytics and a new graphical interface for a comprehensive morphometric experience. Methods in Ecology and Evolution 12, 2355–2363 (2021). 10.1111/2041-210X.13723

57 Collyer, M. L. & Adams, D. C. RRPP: An r package for fitting linear models to high-dimensional data using residual randomization. Methods in Ecology and Evolution 9, 1772–1779 (2018). 10.1111/2041-210X.13029

58 Zelditch, M. L., Sheets, H. D. & Fink, W. L. The ontogenetic dynamics of shape disparity. Paleobiology 29, 139–156 (2003). Doi 10.1666/0094-8373(2003)029<0139:Todosd>2.0.Co;2

59 Alberch, P., Gould, S. J., Oster, G. F. & Wake, D. B. Size and Shape in Ontogeny and Phylogeny. Paleobiology 5, 296–317 (1979). Doi 10.1017/S0094837300006588

60 Piras, P. et al. The role of post-natal ontogeny in the evolution of phenotypic diversity in Podarcis lizards. J Evol Biol 24, 2705–2720 (2011). 10.1111/j.1420-9101.2011.02396.x

61 Fabbri, M. et al. A shift in ontogenetic timing produced the unique sauropod skull. Evolution 75, 819–831 (2021). 10.1111/evo.14190

62 White, H. E. et al. Pedomorphosis in the ancestry of marsupial mammals. Curr Biol 33, 2136–2150 e2134 (2023). 10.1016/j.cub.2023.04.009

63 Collyer, M. L. & Adams, D. C. Phenotypic trajectory analysis: comparison of shape change patterns in evolution and ecology. Hystrix 24, 75–83 (2013). 10.4404/hystrix-24.1-6298

64 Collyer, M. L., Sekora, D. J. & Adams, D. C. A method for analysis of phenotypic change for phenotypes described by high-dimensional data. Heredity (Edinb) 115, 357–365 (2015). 10.1038/hdy.2014.75

65 McNamara, K. J. Heterochrony: the Evolution of Development. Evolution: Education and Outreach 5, 203–218 (2012). 10.1007/s12052-012-0420-3

66 Revell, L. J. phytools 2.0: an updated R ecosystem for phylogenetic comparative methods (and other things). PeerJ 12, e16505 (2024). 10.7717/peerj.16505

67 Cooney, C. R. et al. Mega-evolutionary dynamics of the adaptive radiation of birds. Nature 542, 344–347 (2017). 10.1038/nature21074

68 Lautenschlager, S. From bone to pixel—fossil restoration and reconstruction with digital techniques. Geology Today 33, 155–159 (2017). 10.1111/gto.12194

69 Crane, A. et al. Taphonomic damage obfuscates interpretation of the retroarticular region of the Asteriornis mandible. Geobios (2024). 10.1016/j.geobios.2024.03.003

70 Rahman, I. A. & Lautenschlager, S. Applications of Three-Dimensional Box Modeling to Paleontological Functional Analysis. The Paleontological Society Papers 22, 119–132 (2017). 10.1017/scs.2017.11

71 Clavel, J., Escarguel, G. & Merceron, G. mvmorph: an r package for fitting multivariate evolutionary models to morphometric data. Methods in Ecology and Evolution 6, 1311–1319 (2015). 10.1111/2041-210X.12420

72 Ducatez, S. & Field, D. J. Disentangling the avian altricial-precocial spectrum: Quantitative assessment of developmental mode, phylogenetic signal, and dimensionality. Evolution 75, 2717–2735 (2021). 10.1111/evo.14365

73 Orkney, A., Rothier, P. S. & Hedrick, B. P. Great expectations: altricial developmental strategies are associated with more flexible evolution of limb skeleton proportions in birds. Proceedings of the Royal Society B: Biological Sciences 292 (2025). 10.1098/rspb.2025.1647

74 Berv, J. S. et al. Genome and life-history evolution link bird diversification to the end-Cretaceous mass extinction. Science Advances 10, eadp0114 (2024). doi:10.1126/sciadv.adp0114

