## Supplementary material for "Macroevolutionary shifts in post-hatching ontogeny and the origin of craniofacial disparity in fowl (Aves: Galloanserae)": Figures and Supp. Materials

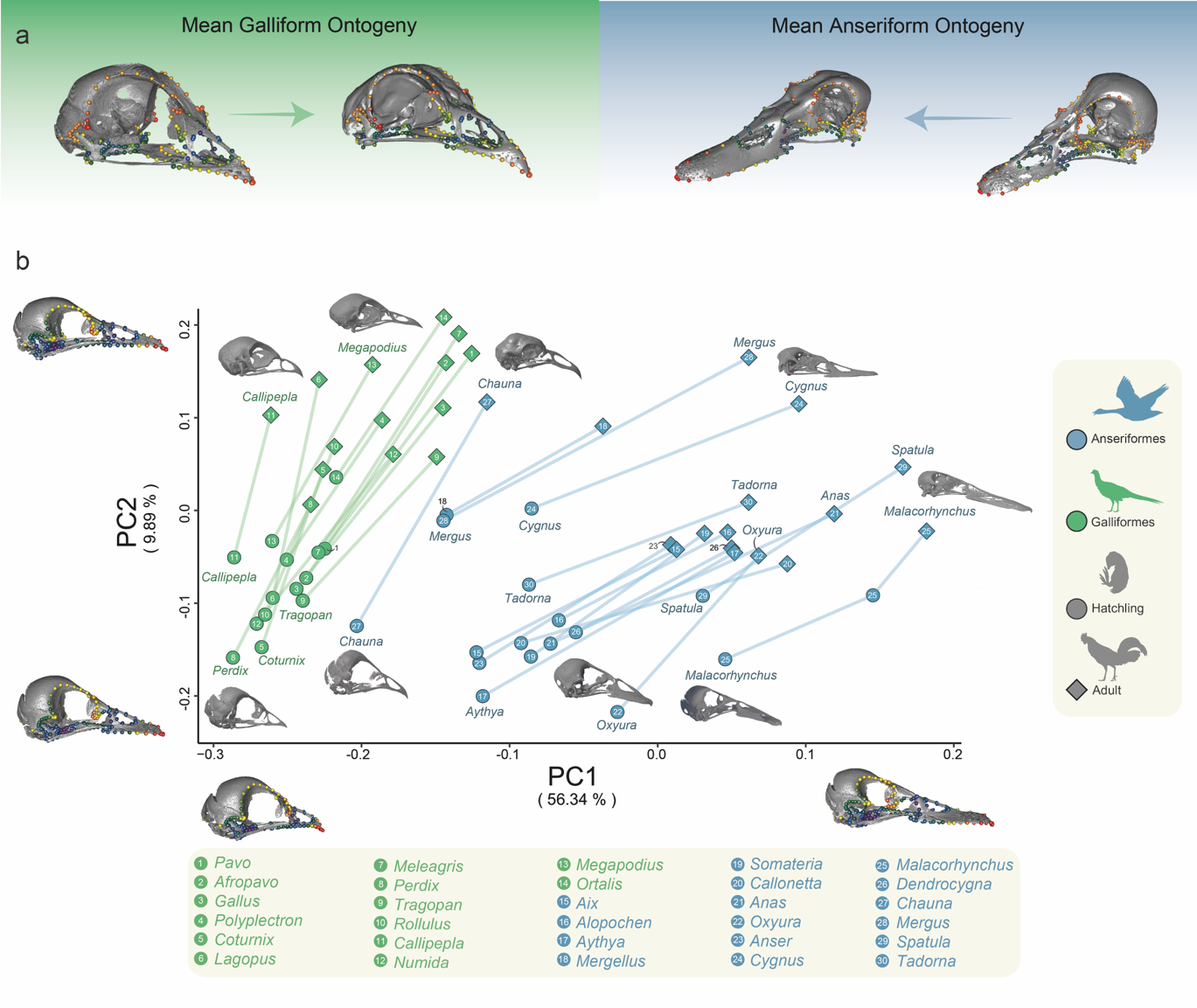

### Figures

**Figure 1: Cranial ontogenetic morphospace of crown Galloanserae.** a) Illustration of mean post-hatching ontogenetic shape changes from hatchling to adults in crown Anseriformes and Galliformes. b) PCA plot of cranial disparity at hatchling and adult stages in crown Galloanserae, showing intra- and interclade ontogenetic variance in skull shape. Galliformes and Anseriformes occupy distinct regions of morphospace, with the screamer *Chauna* (Anseriformes: Anhimidae) occupying an intermediate position between the morphospace circumscribed by other Anseriformes, and Galliformes morphospace.

**Figure 2: Heatmaps of ontogenetic shape variance and shape convergence/divergence in the skulls of crown-group Galloanserae.** a) Heatmap showing variance in pairwise Procrustes distance between hatchlings of Galloanserae species. b) Heatmap showing the variance in pairwise Procrustes distance between adults of Galloanserae species. c) Heatmap showing the interspecific degree of pairwise ontogenetic shape convergence or divergence between Galloanserae skulls. Portions of phylogenetic topologies in blue are Anseriformes, portions of phylogenetic topologies in green are Galliformes.

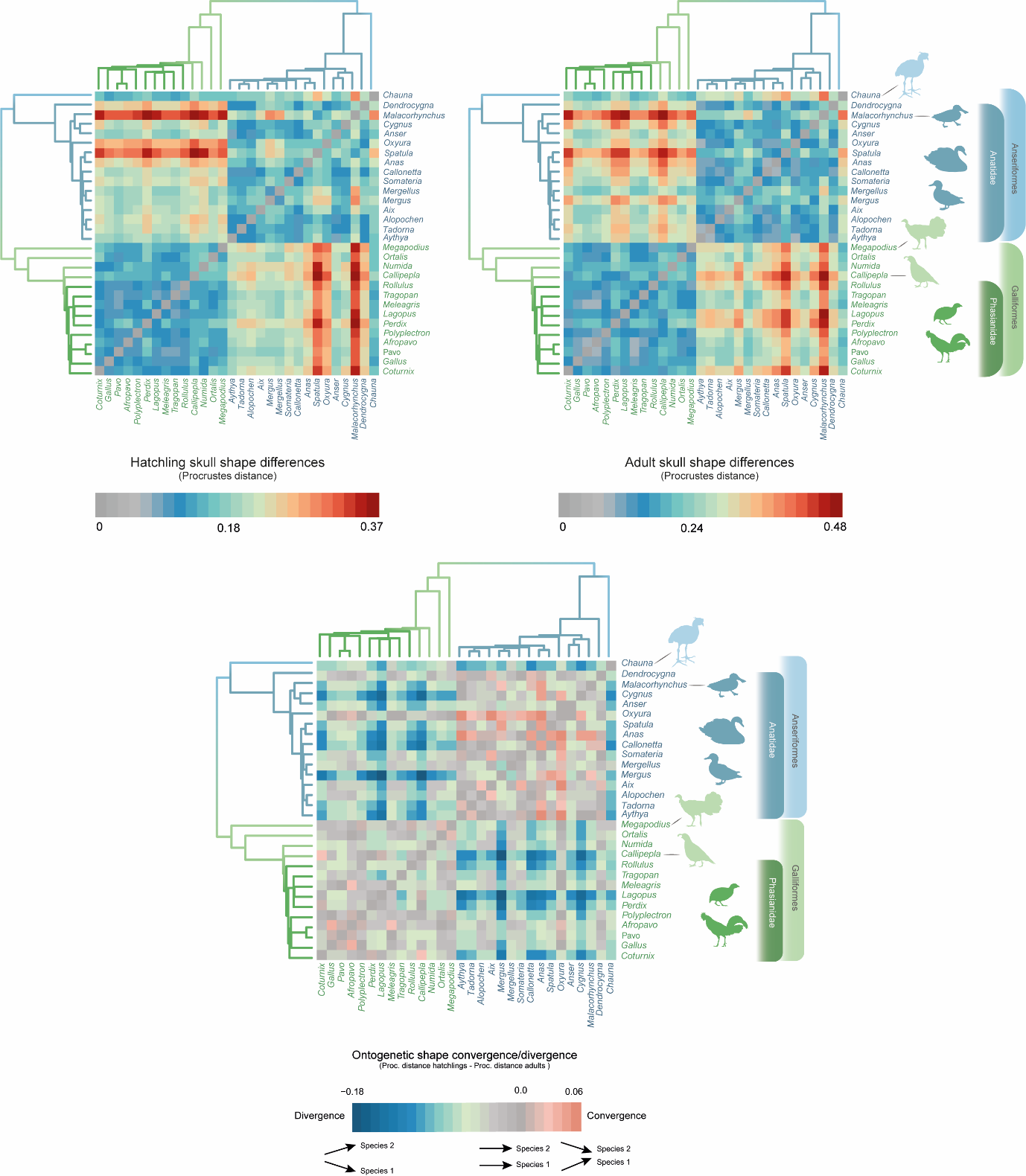

a

b

c

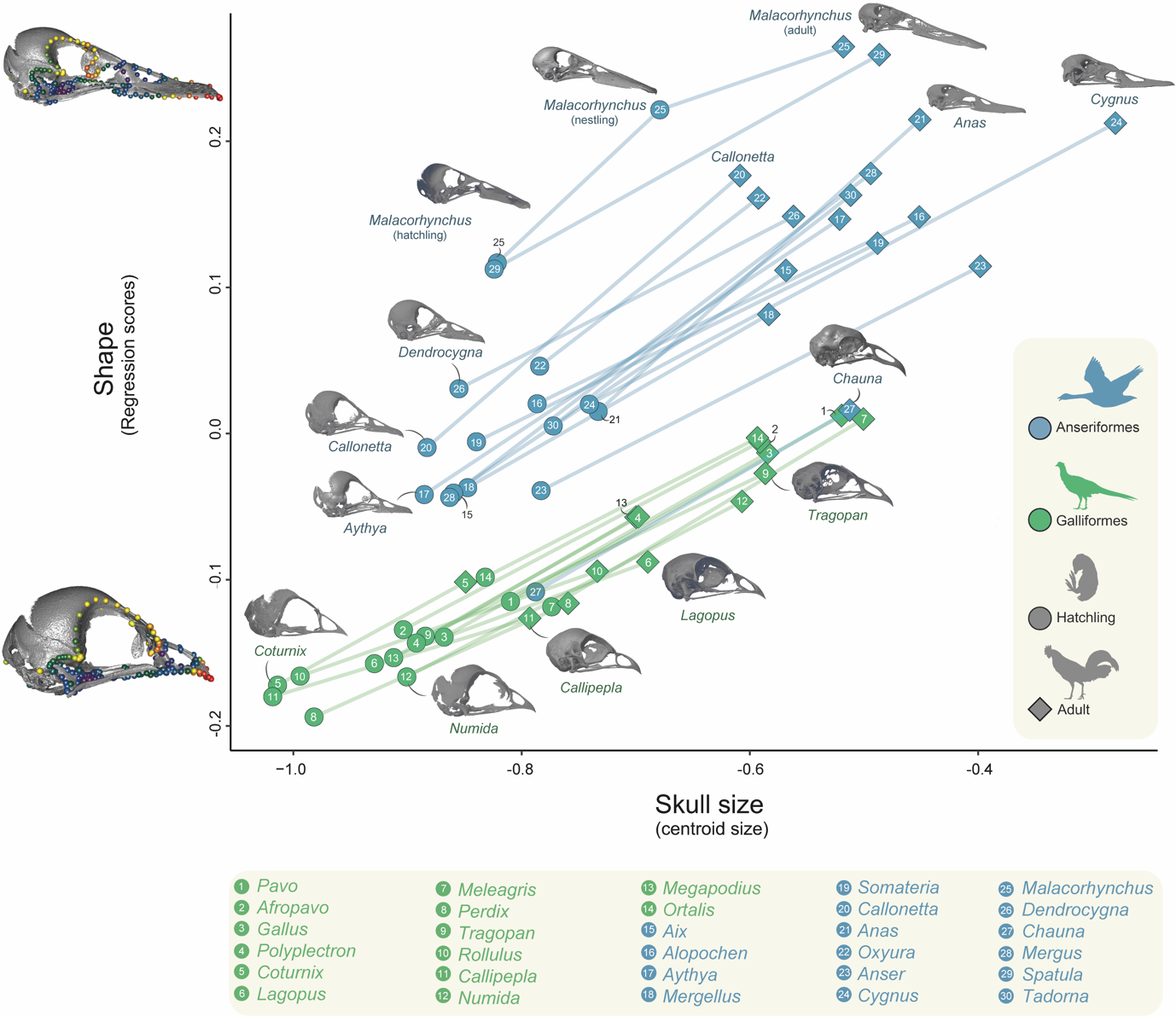
**Figure 3: Allometric trajectories of post-hatching cranial ontogeny across crown Galloanserae.** Trajectories were calculated by regressing shape and size for all hatchling and adult skull pairs. Anseriformes and Galliformes occupy different regions of morphospace, with the exception of *Chauna* (Anseriformes: Anhimidae), which overlaps with galliforms. Skull shape changes associated with the common allometric regression vector are shown on the shape axis.

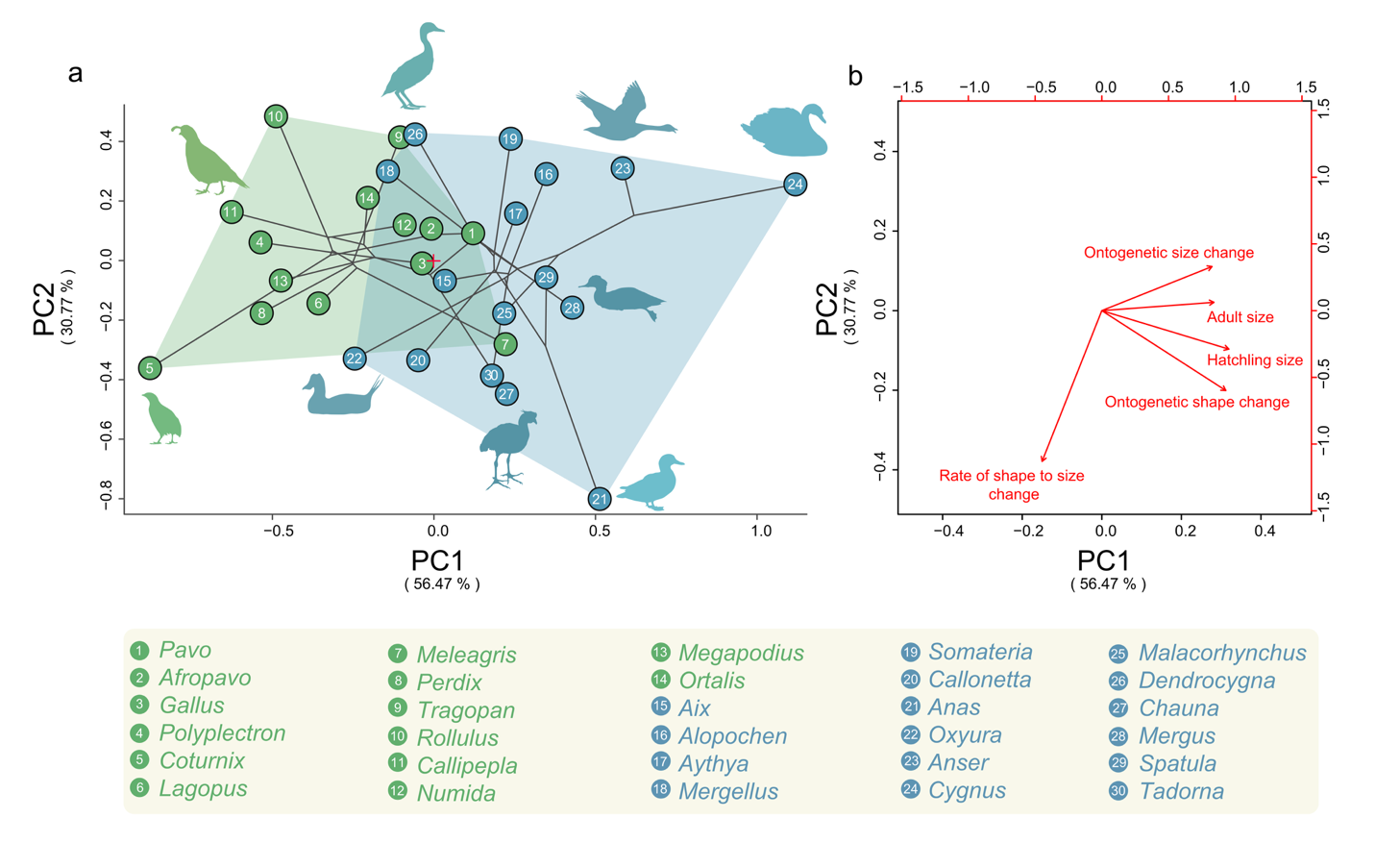
**Figure 4: Galloanseran skull ontogenetic phylomorphospace.** a) Phylogenetic clustering of anseriform and galliform cranial ontogenies and their intraclade ontogenetic variance. The origin (0,0) of the projected axes is shown with a red cross. b) biplot showing the ontogenetic traits accounting for ontogenetic variance within Galloanserae. Anseriform ontogenetic trajectores are more variable than those of galliforms, with high variance in the amount of ontogenetic shape and size change, while galliform ontogenetic variability is mostly limited to intraspecific differences in ontogenetic size change.

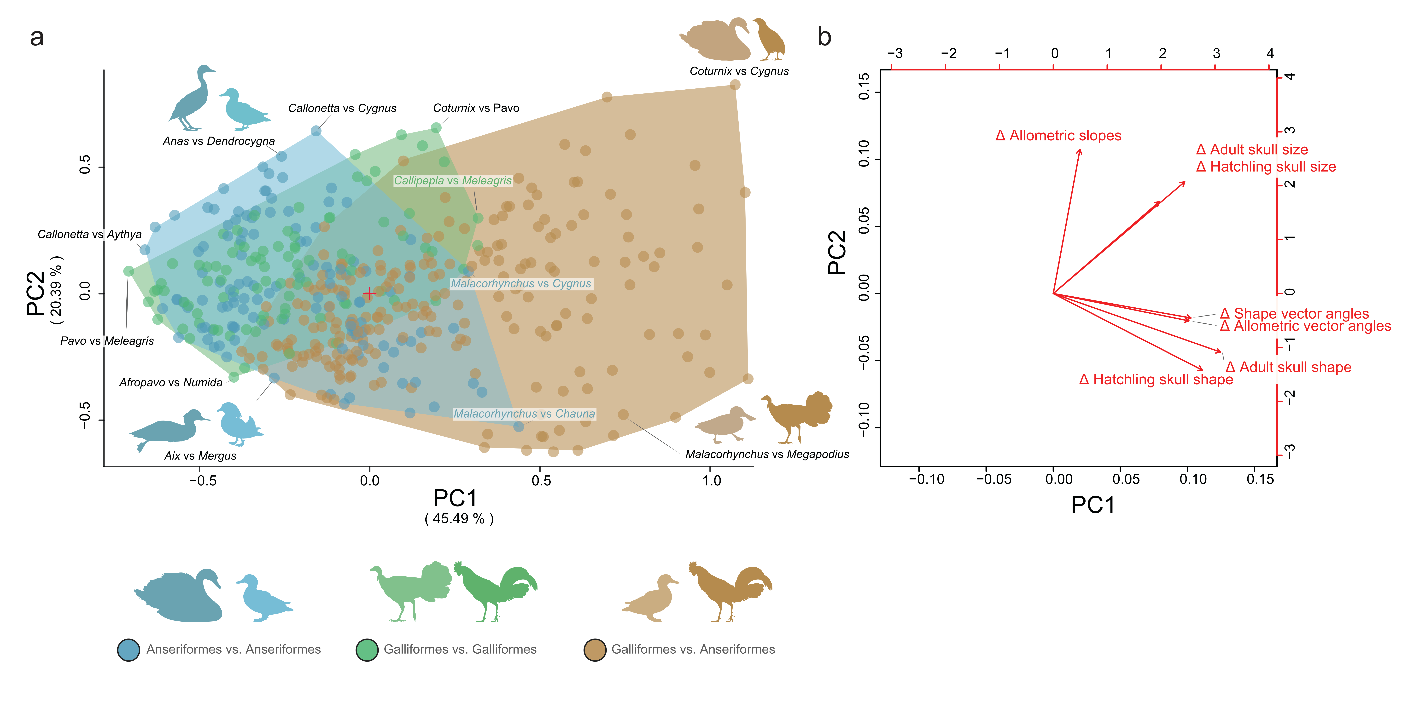
**Figure 5: Pairwise ontogenetic morphospace of crown Galloanserae skulls.** a) Distribution of ontogenetic variance in inter- and intraclade comparisons. Interclade (i.e. galliform vs. anseriform; in brown) comparisons exhibit markedly greater variance than intraclade comparisons (i.e. pairwise comparisons within Galliformes, in green, or Anseriformes, in blue). Intraclade comparisons largely overlap, indicating that the traits underpinning differences among anseriforms are generally the same traits underpinning differences among galliforms. anseriforms differ from other anseriforms in similar traits than galliforms differ from other galliforms. By contrast, interclade comparisons do not overlap in to the same degree with as intraclade comparisons, revealing that anseriforms differ from galliforms in other a broader set of ontogenetic traits than distinguish within-clade comparisons: galliform-galliform or anseriform-anseriform pairs. The origin (0,0) of the projected axes is shown with a red cross. b) biplot showing ontogenetic traits accounting for pairwise ontogenetic differences within Galloanserae. Most ontogenetic traits, especially skull shape and size, vary between Galliformes and Anseriformes, while skull shape variance is greater in Anseriformes and skull size variance is greater in Galliformes.

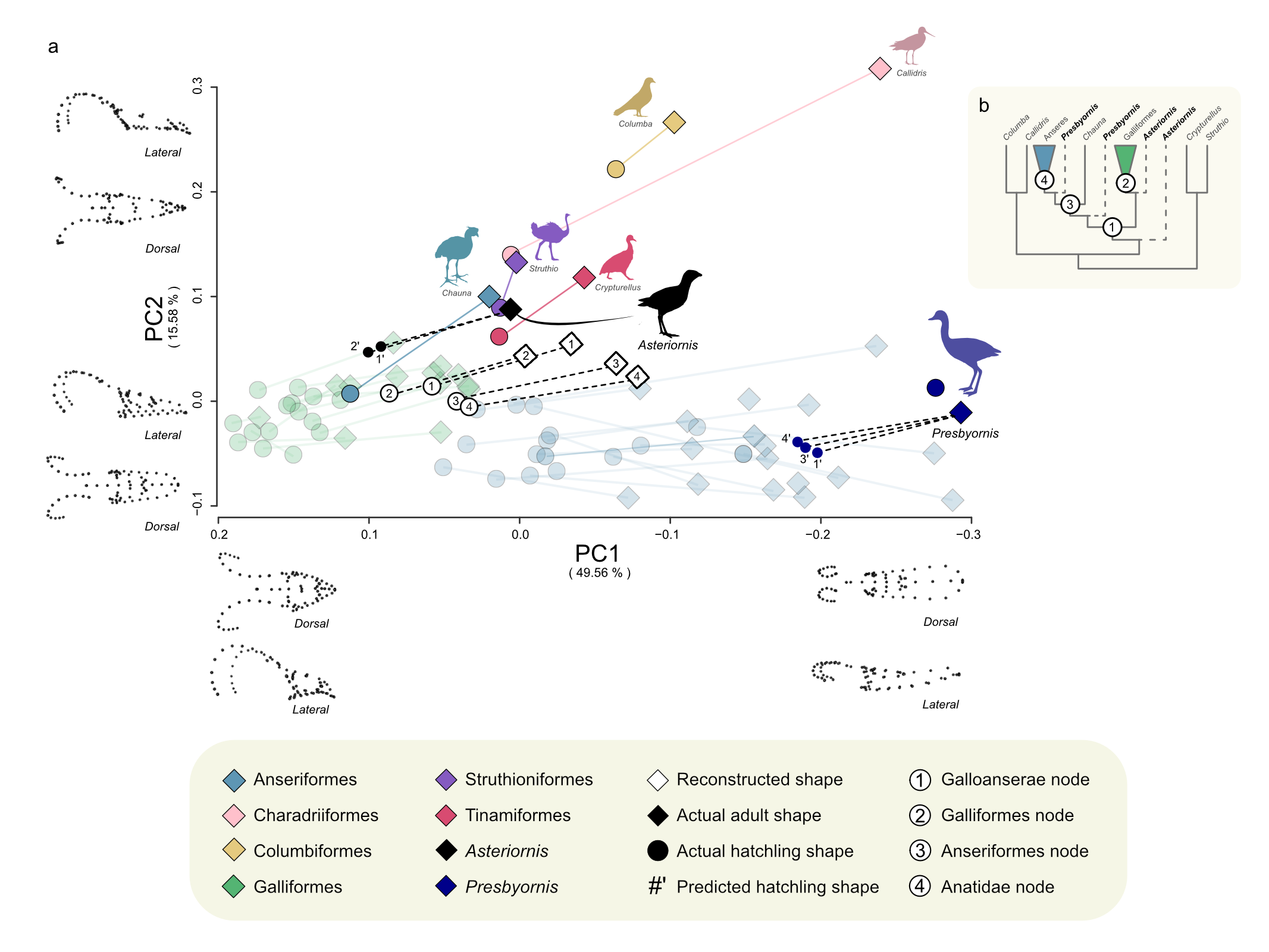
**Figure 6: Expanded cranial ontogenetic morphospace of Neornithes with a focus on Galloanserae.** a) PCA plot of cranial disparity at hatchling and adult stages of crown group birds. In addition to Galloanserae, the fossil total clade galloanserans *Presbyornis* and *Asteriornis* are included (see methods for description of estimating hatchling shape for fossils), as well representatives of the major extant bird clades Palaeognathae and Neoaves and projections of reconstructed ancestral shapes. b) simplified phylogeny of Neornithes showing relationships among taxa in a).

### Extended Data Figures

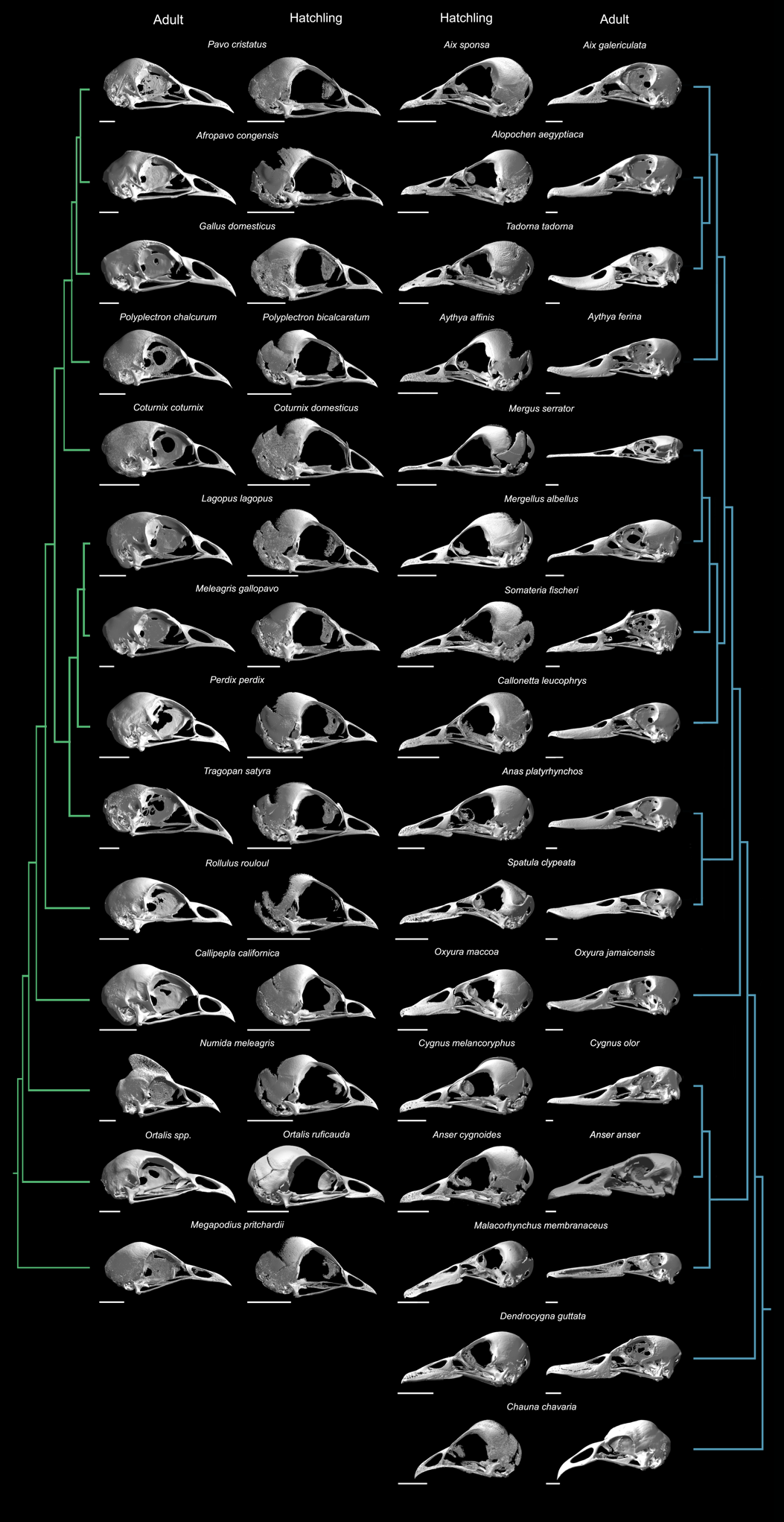

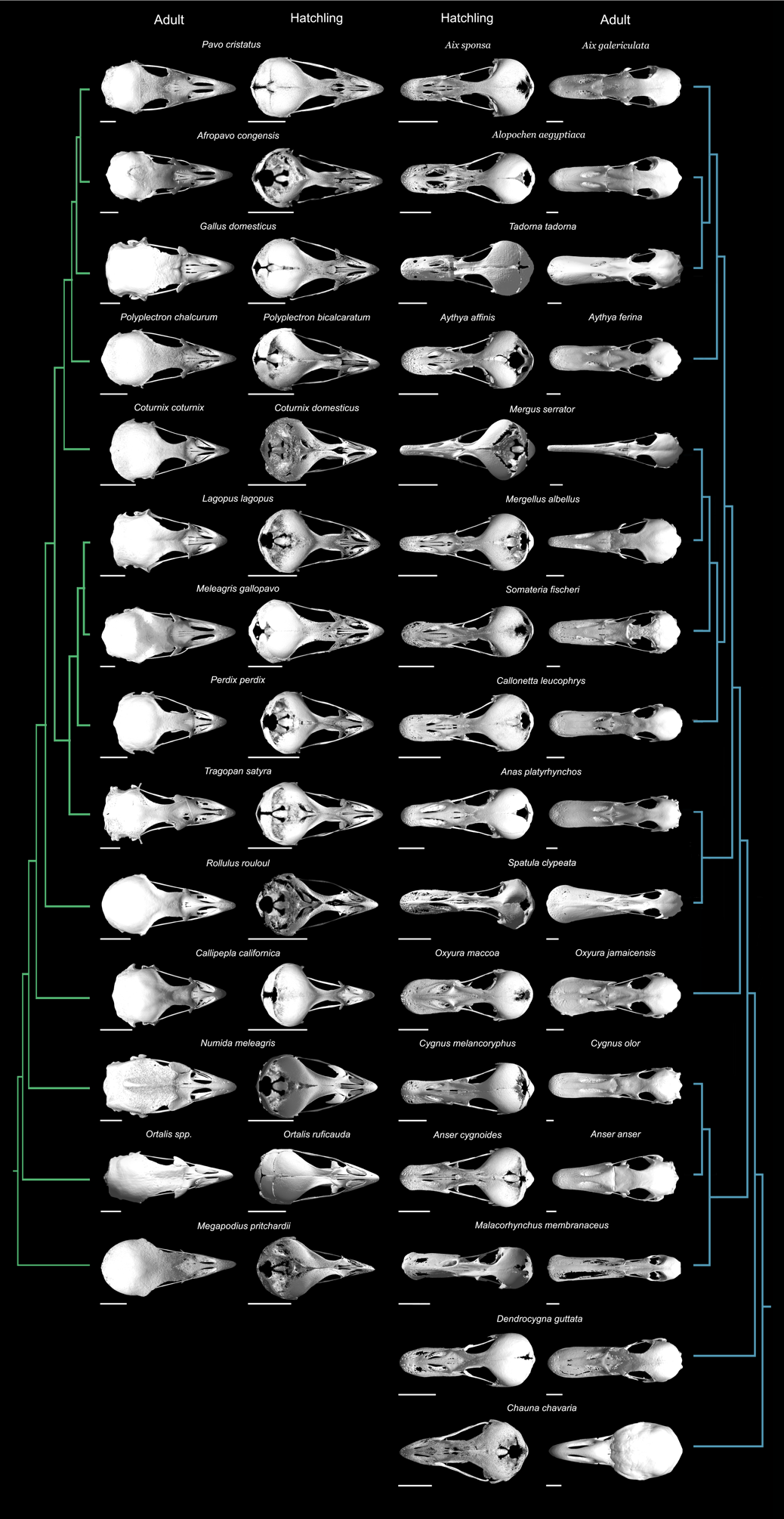
**Extended Data Figure 1: Lateral view of the crown Galloanserae skulls used in this study (Dataset 1).**

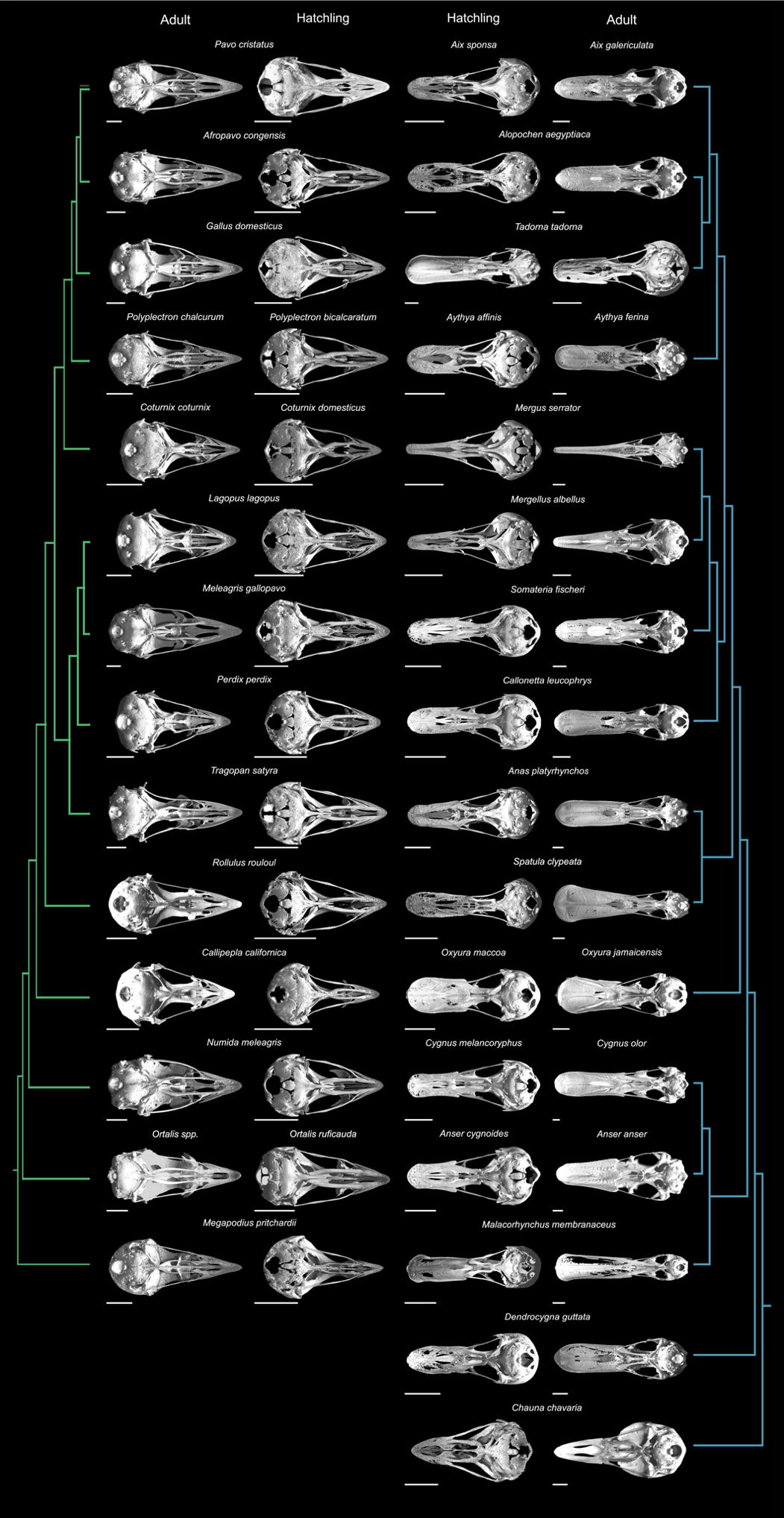
**Extended Data Figure 2: Dorsal view of the crown Galloanserae skulls used in this study (Dataset 1).**

**Extended Data Figure 3: Ventral view of the crown Galloanserae skulls used in this study (Dataset 1).**

**Extended Data Figure 4: Cranial shape convergence and divergence heatmaps in the post-hatching ontogeny of crown Galloanserae.** a) Heatmap showing the interspecific pairwise variance in the angle between ontogenetic shape vectors. b) Heatmap showing the interspecific pairwise ontogenetic convergence, or divergence between Galloanserae skulls.

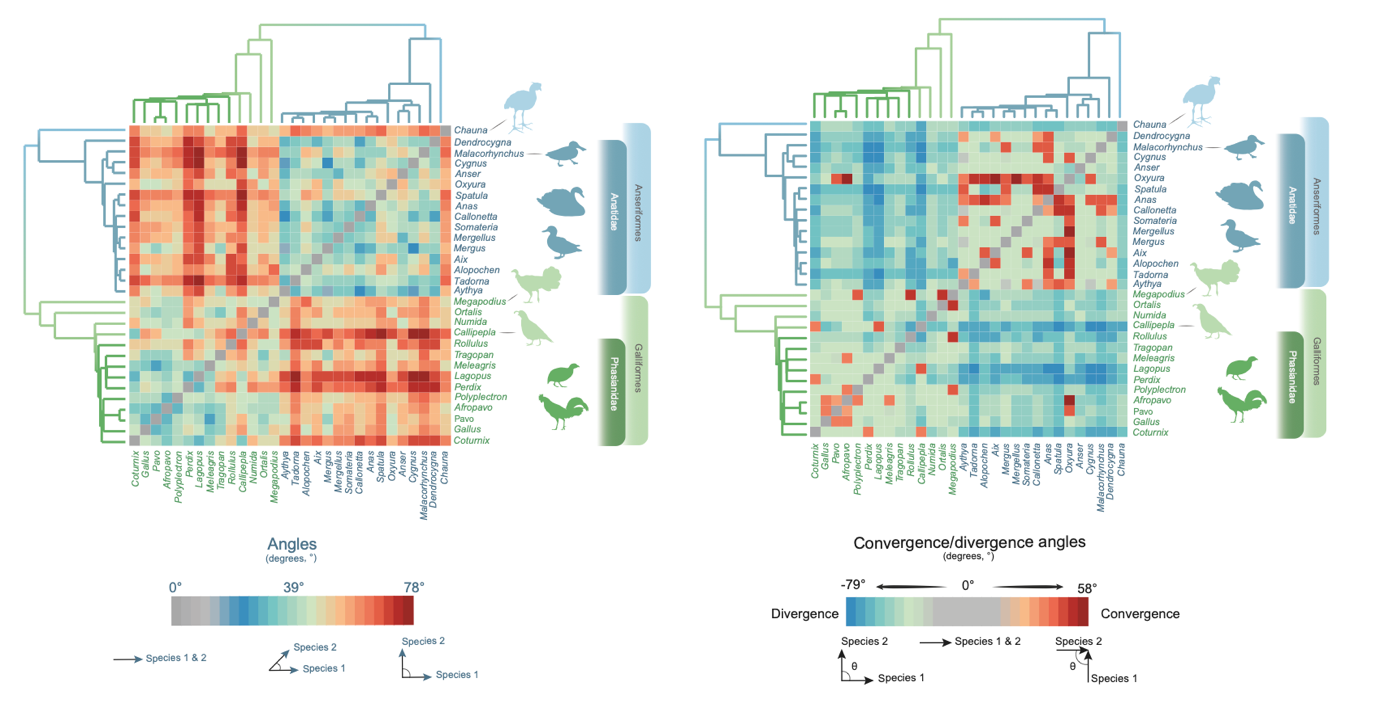

a

b

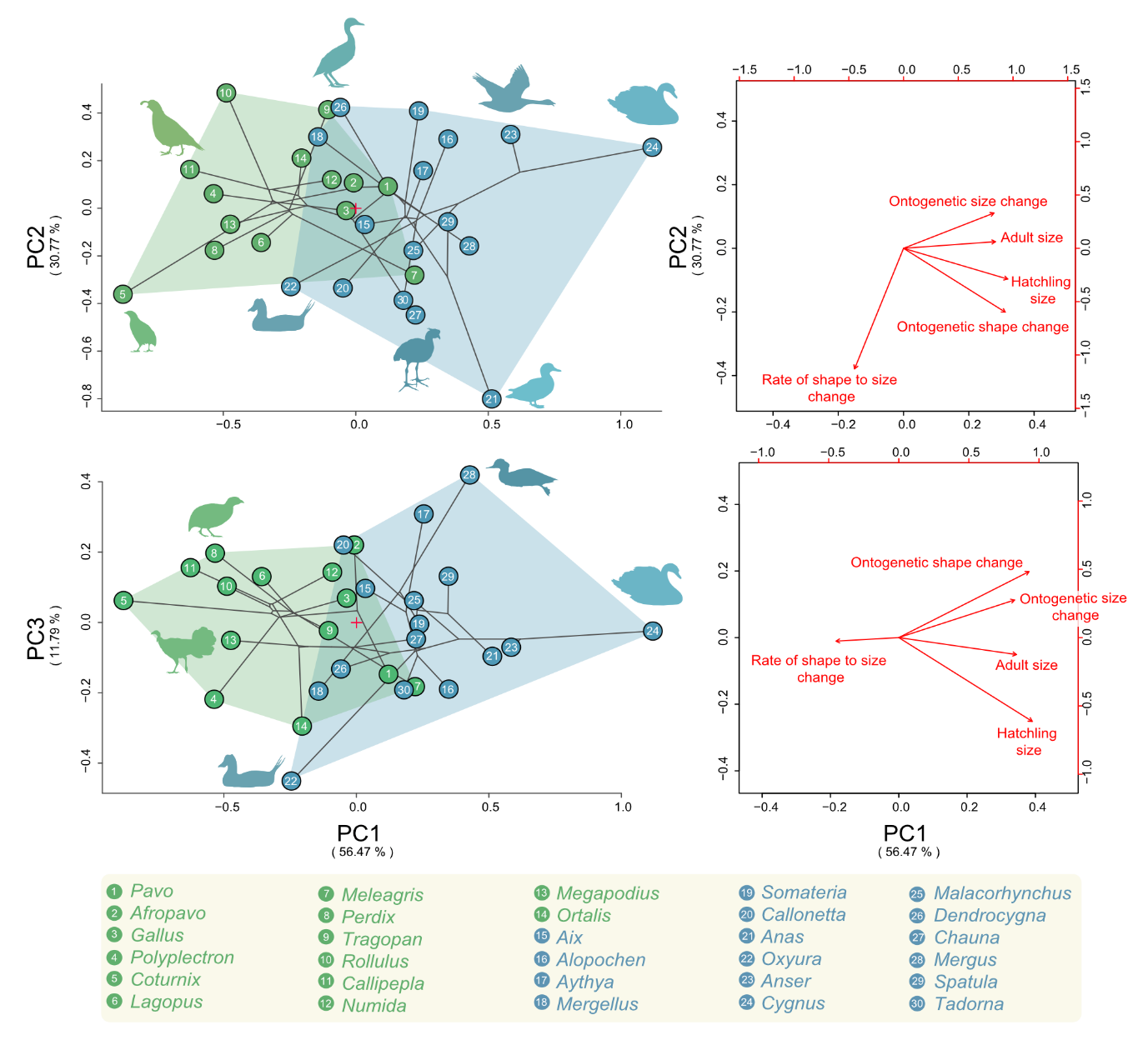

a

b

**Extended Data Figure 5: Ontogenetic-phylomorphospace** **of crown Galloanserae skulls.** a: Left)	The phylogenetic clustering of anseriform and galliform cranial ontogenies and intraclade ontogenetic variance. Right) biplot showing the ontogenetic traits accounting for ontogenetic variance within Galloanserae. b: similar to (a) but showing a different projection of the morphospace with PC1 and PC3. The origins (0,0) of the projected axes are indicated with a red cross.

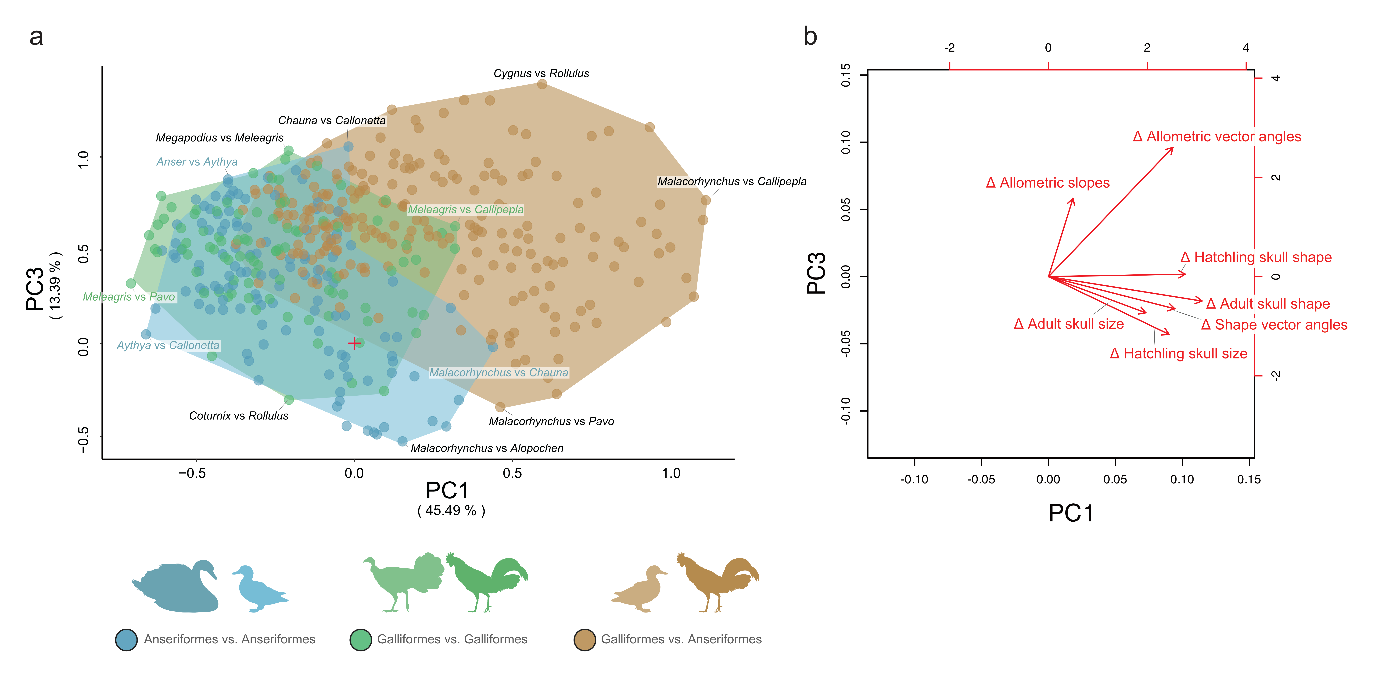
**Extended Data Figure 6: Alternative projection of Pairwise-Ontogenetic-Morphospace of crown Galloanserae skulls.** Left) Distribution of inter- and intraclade differences with the morphospace showing the difference between inter- and intraclade ontogenetic variance. Right) biplot showing the difference in ontogenetic traits accounting for pairwise ontogenetic differences within Galloanserae. △ symbolizes difference in a trait. The origin (0,0) of the projected axes is shown with a red cross.

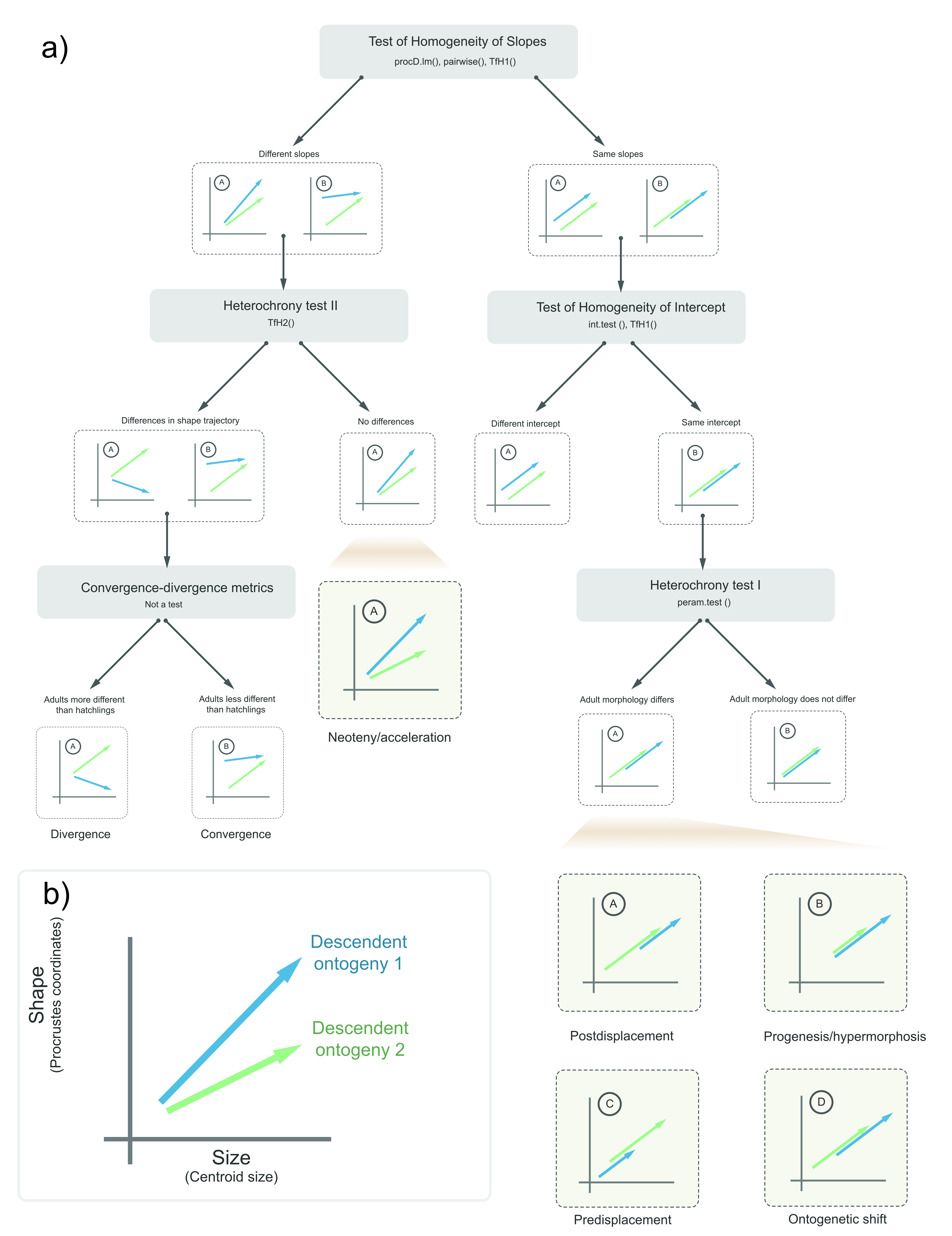
**Extended Data Figure 7: Flow-chart of allometric analysis used for determining the allometric differences between clades.** a) flow-chart illustrating the series of tests that was carried out and the possible directions of analysis based on the results of each test. b) legend showing the colour scheme for differentiating between ontogenies.

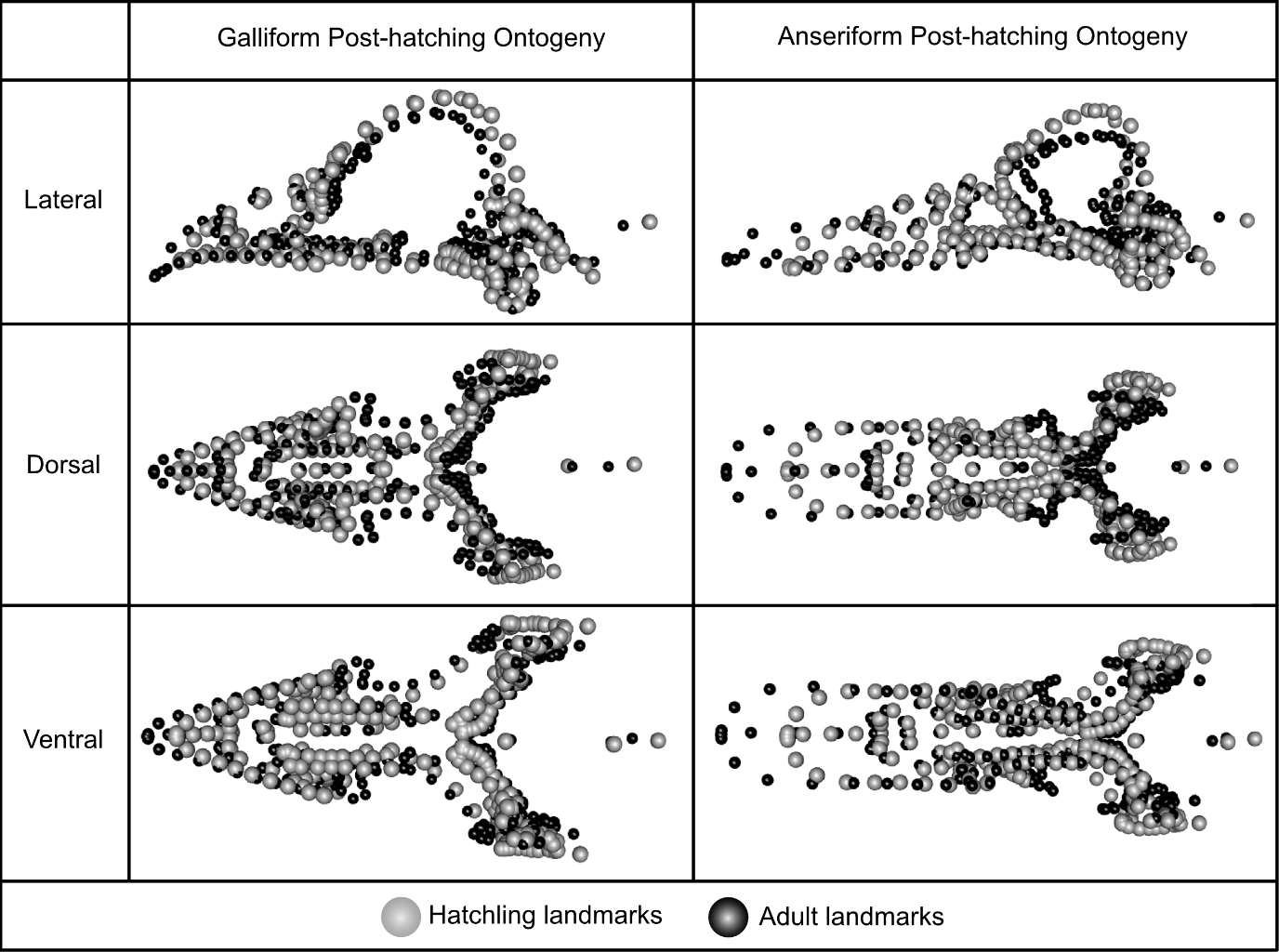
**Extended Data Figure 8: Mean cranial post-hatching ontogenetic change in crown Galliformes and Anseriformes.** Average landmark configurations of hatchling skull shape for galliform and anseriform hatchlings, superimposed on the average landmark configurations of adult galliforms and anseriforms. The difference in the displacement of the landmarks illustrates intraclade post-hatching cranial ontogenetic change, and the differences in these displacements between the left and right columns illustrates interclade variance in cranial ontogenetic change.

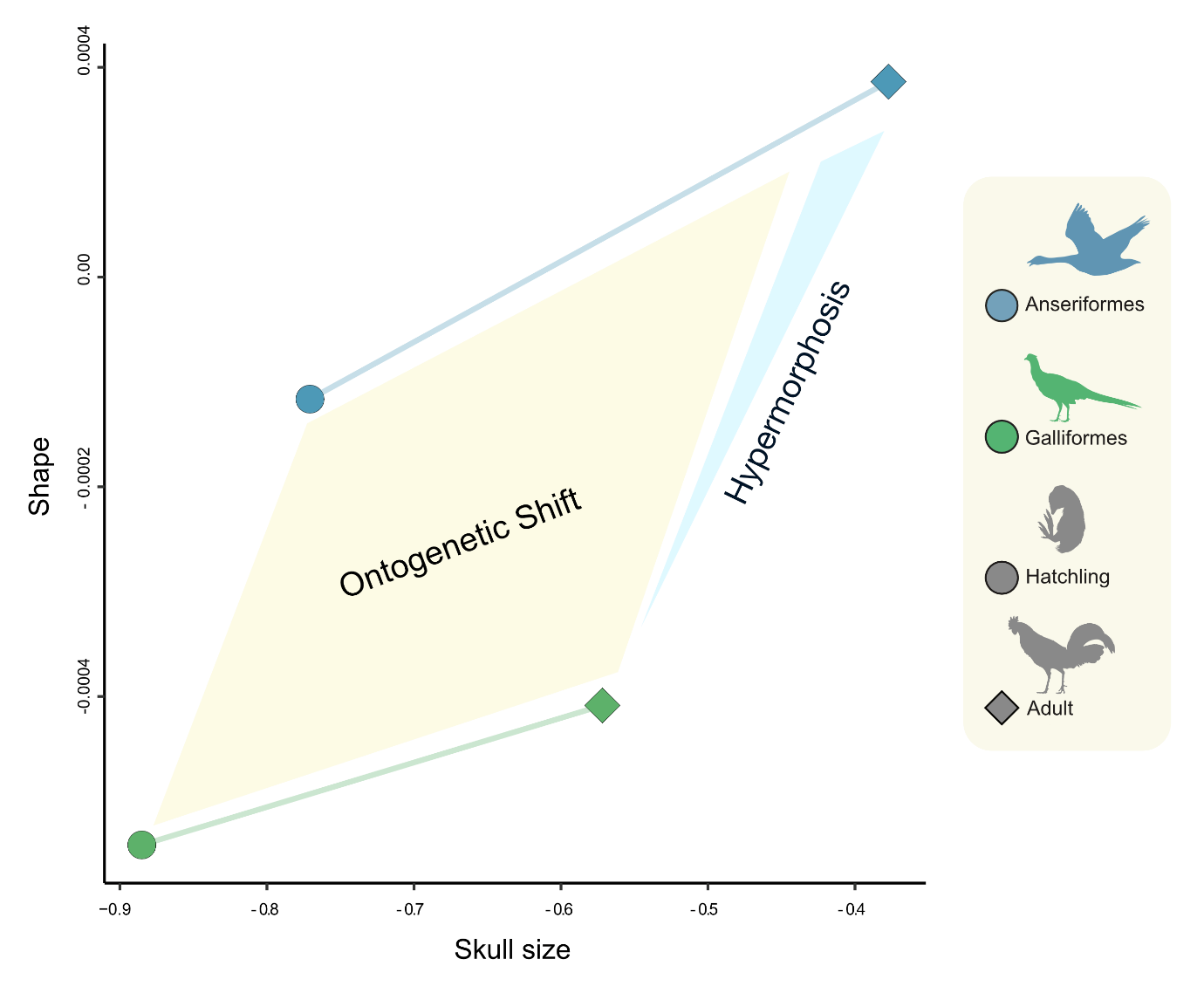
**Extended Data Figure 9: Illustration of heterochronic differences in the post-hatching ontogeny of Galliformes and Anseriformes.** The mean size and shapes of each clade were used to illustrate the combined roles of interclade ontogenetic shift and relative hypermorphosis in differentiating anseriform from galliform post-hatching ontogenetic trajectories.

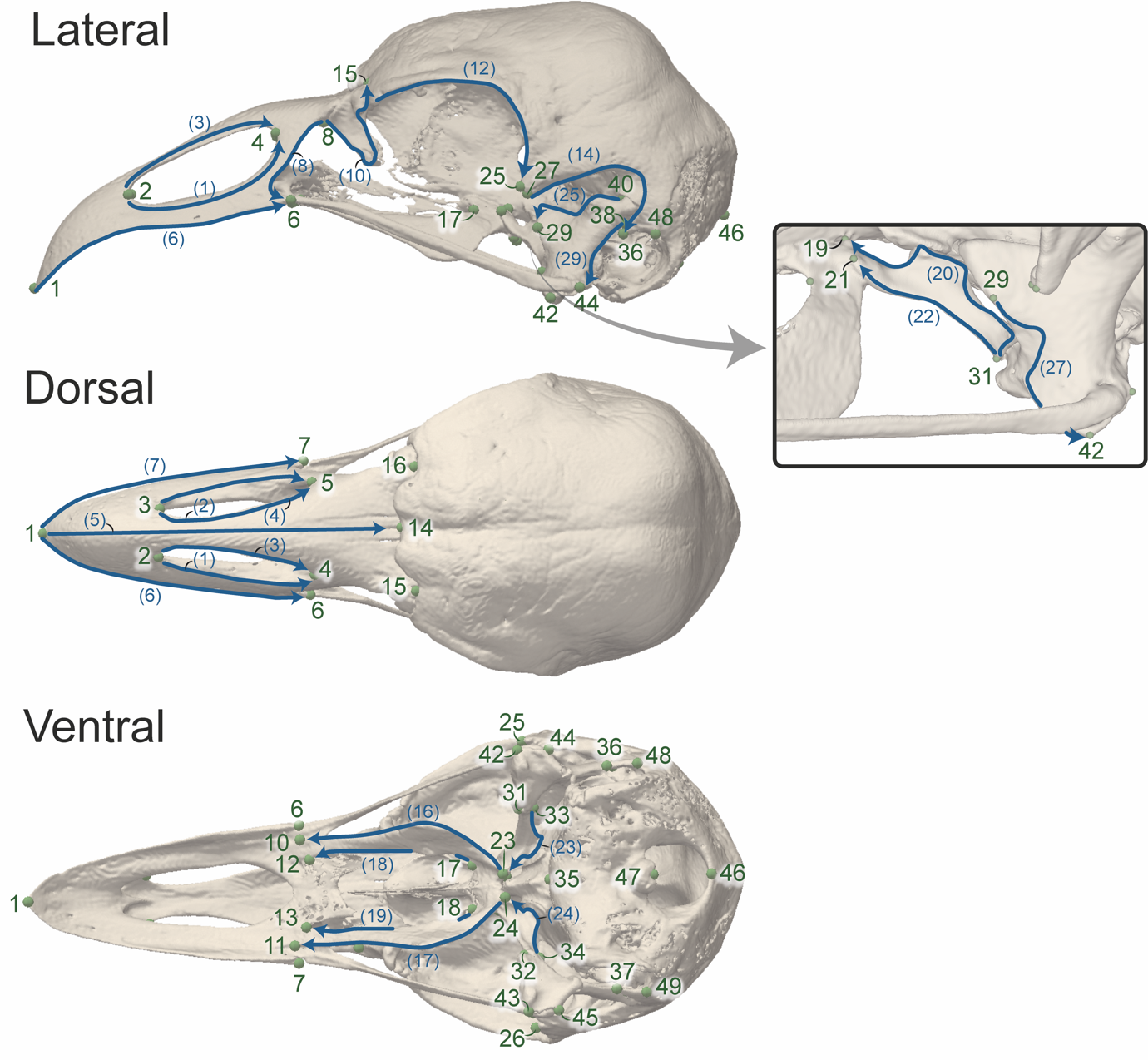
**Extended Data Figure 10: Landmark and semilandmark configuration used for analysis of Galloanserae skull disparity illustrated with an adult *Chauna chavaria* skull.** The fixed landmarks are shown in green, and the curves of semi-landmarks are shown in blue. The number of semi-landmarks per curve is shown in brackets.

### Supplementary materials

Supplementary Results

*Extended Results. Detailed patterns of shape variation.*

Adult galliform lacrimals exhibit proportionally wider heads than hatchlings, while adult anseriform lacrimals have a longer head and descending process, which extends caudally. Caudal to the lacrimal, the adult orbit is relatively smaller than that of hatchlings, but in Anseriformes the dorsal side of the orbit appears flat at the adult stage (Extended Data Figure 8). On the other side of the orbit, the postorbital process appears longer at the adult stage in both clades, but the galliform process is caudally directed. Ventral to the orbit, the palatines appear proportionally longer in the adults of both clades than in hatchlings, but the caudal end of the palatine is relatively wider in anseriform adults than in galliforms (Extended Data Figure 8). Caudal to the palatal region, the braincase appears narrower and shallower in the adults of both clades relative to hatchlings (Extended Data Figure 8).

Moreover, the lacrimals of galliforms exhibit laterally extended heads with a tapering descending process and ventrally-directed foot, while anseriforms have rostrocaudally extended lacrimal heads, a wider descending process, and foot that directed ventrocaudally. The orbit is larger relative to skull size, and the braincase longer and wider in Galliformes with respect to Anseriformes. Other distinguishing interclade features among hatchlings include a more robust quadrate with thicker articular processes in Anseriformes relative to Galliformes. Furthermore, the caudal end of the palatines is wider in Anseriformes than in Galliformes, and anseriforms exhibit a distinctive rostral process of the pterygoid (Figure 1 & Extended Data Figure 8).

*Extended Results. Detailed patterns of ontogenetic variation.*

Regarding intraclade ontogenetic variance, a few Anseriformes clades have distinct ontogenetic trajectories. *Cygnus* has one of the largest skull sizes and ontogenetic size change, but one of the smallest allometric rates. *Anas* also has a large hatchling skull size, but with greatest amount of ontogenetic shape change and allometric rate. *Callonetta* and *Aythya* have one of the smallest hatchling skull sizes, but the former also has the smallest adult skull size. *Oxyura* exhibits the smallest amount of ontogenetic size change, and *Mergellus*, along with *Dendrocygna,* has the smallest amount of ontogenetic shape change. *Mergus,* which is sister to *Mergellus,* has one of the greatest amounts of ontogenetic shape change. The ontogeny of *Somateria* is also distinguishable by its low allometric rate. The ontogeny of *Chauna,* with its distinctive galliform-like skull shape, exhibits average values for hatchling and adult skull sizesand amount of ontogenetic size change, but a greater allometric slope and amount of ontogenetic shape change than reconstructed for the last common ancestor of Anseriformes and Anatidze.

Within Galliformes, *Meleagris* and *Pavo* have two of the largest skull sizes, with the former exhibiting one of the greatest amounts of ontogenetic shape change in our sample. *Ortalis* also has one of the largest hatchling skull sizes, although it is otherwise comparable to other galliforms in most respects. This is notable inasmuch that, as a representative of Cracidae, it may be considered the most developmentally altricial taxon in our dataset^72^, a parameter increasingly recognised as a correlate of increased evolvability in birds^27,73,74^. *Coturnix* and *Callipepla* have one of the smallest skull sizes in our sample, with the former exhibiting the smallest amount of ontogenetic size change and greatest allometric rate in our galliform sample. *Afropavo* and *Gallus* also exhibit among the greatest degrees of ontogenetic shape change, which might be to be associated with adult skull size as *Gallus* has one of the largest adult skull sizes in our galliform sample and *Afropavo* exhibits one of the greatest amounts of ontogenetic size change. *Rollulus* and *Polyplectron* exhibit some of the lowest amounts of ontogenetic shape change, with the former exhibiting one of the lowest allometric rates in our sample. The ontogenetic trajectory of *Tragopan* similarly exhibits an unusually low allometric rate.

Supplementary discussion

Methodological limitations to deducing ontogenetic convergence and divergence

The analysis of ontogenetic convergence/divergence between clades was limited by the availability of only one sample per species at each stage. Moreover, we used a number of methods for quantifying ontogenetic convergence/divergence due to the limitations in each of them. For example, the distance-based and angle-based methods of determining ontogenetic convergence/divergence differ in their accuracy and the conclusions that can be drawn from them. The angle-based method quantifies the angular distance between two points, representing two adult shapes, in shape space. The angular distance between two points, i.e. two adult shapes, neglects the direction of ontogenetic vectors, whether convergent or divergent (Supplementary Figure 6a&b). Equally convergent and divergent trajectories have the same angular distance, because the angular distance triangle is based on the adult shape points and the intersection of the vectors emanating from them (Supplementary Figure 6a&b). Another limitation is that the angular distance between two adult shape points remains the same regardless of how far the hatchling points are from the adult points, for as long as they remain within the angular distance triangle (Supplementary Figure 6c&d). Subsequently, a highly or slightly divergent trajectories would have similar angular distances (Supplementary Figure 6). For example, the ontogenetic vector of *Afropavo* appears highly convergent with *Oxyura* in the angle-based heatmaps (Extended Data Figure 4a), while they appear only slightly convergent in the distance-based heatmap (Figure 2c).

Similarly, the distance-based method also has a few limitations. When comparing two pairs of ontogenies, if the interhatchling and interadult distances of a pair is a multiple of the other pair, the amount of convergence, or divergence, will be equal despite the much larger interhatchling and interadult shape distances of the first pair (Supplementary Figure 6e&f). This limitation is exemplified by the relative divergence of *Chauna* from *Callonetta*, and *Coturnix* from *Callonetta.* The distance-based heatmap indicates that the divergence between these two pairs is equal (Figure 2c), while the angle-based heatmaps show that *Chauna* and *Callonetta* have more divergent ontogenies than *Coturnix* from *Callonetta* (Extended Data Figure 4a).

Another limitation of the distance-based method is that it overlooks the relative orientation of a pair of ontogenetic vectors, obfuscating the nature of the divergence between two clades. When two ontogenetic vectors have proximate origins in morphospace and similar orientations, but different lengths, the nature of their divergence appears different using different methods. The similarity of the vectors’ orientations would produce a low angular distance between them, indicating low divergence (Supplementary Figure 6). On the other hand, the large distance between their adults would indicate high divergence (Supplementary Figure 6g&h). This issue is seen in the divergence between *Mergus* and *Mergellus*, where the distance-based heatmap shows that these two species are more divergent from one-another than either of them are to other anatids. However, the skull shapes of *Mergus* and *Mergellus* are most like each other within our anatid sample (Figure 1b, & Supplementary Figure 2-4), indicating the extent of the divergence shown in the angle-based heatmaps (Extended Data Figure 4a) is more realistic.

These methodological limitations indicate the importance of considering the results of the shape vector space (Figure 1), as well as the distance-based, the angle-based and signed angle-based heatmaps (Figure 2, Extended Data Figures 4) to yield an accurate picture of ontogenetic convergence/divergence.

*Limitations of deducing ancestral ontogenies*

Predicting the hatchling shape of extinct ancestral taxa poses several challenges which need to be addressed in further studies. One of these difficulties is the reliance on deformed or reconstructed adult fossil skulls, which might bias the prediction of hatchling skull shape. A systematic examination of the impact of skull deformation, and retrodeformation approaches, on hatchling skull prediction is needed. Another difficulty is in choosing the ontogenetic vector to be applied to the adult fossil skull shape. In our study, the ancestral shapes were similar to one-another (Supplementary Figure 8) which would lead to similar ontogenetic vectors, leading to similar hatchling shapes. Therefore, a systematic determination of the range of plausible vectors and plausible hatchling skull shapes is needed. Finally, the precision and accuracy of predicted hatchling skull shape could be influenced by the ontogenetic variance of a clade, because highly disparate clade, such as Anseriformes, would predict a wider range of hatchling shapes than a more constrained clade, such as Galliformes. Therefore, a systematic study of the influence of clade disparity on predicting skull shapes is needed along with the use of fossil clades with known hatchling skull shapes.

### Supplementary tables

**Table S1: Pairwise absolute differences between mean skull shapes of crown galliform and anseriform hatchlings and adults. †p<0.01**

|  | Anseriformes Adults | Anseriformes Hatchlings | Galliformes Adults | Galliformes Hatchlings |
| --- | --- | --- | --- | --- |
| Anseriformes Adults | 0 |  |  |  |
| Anseriformes Hatchlings | **0.16†** | 0 |  |  |
| Galliformes Adults | **0.25†** | **0.16†** | 0 |  |
| Galliformes Hatchlings | **0.31†** | **0.18†** | **0.13†** | 0 |

**Table S2: Procrustes variances in skull shape of crown galliform and anseriform hatchlings and adults.**

| Adult Anseriformes | Adult Galliformes | Anseriformes Hatchlings | Galliformes Hatchlings |
| --- | --- | --- | --- |
| 0.04568827 | 0.02316714 | 0.01662196 | 0.02982064 |

**Table S3: Pairwise absolute differences between Procrustes variances in skull shape of crown galliform and anseriform hatchlings and adults. †p<0.01, * p<0.05**

|  | Adult Anseriformes | Adult Galliformes | Anseriformes Hatchlings | Galliformes Hatchlings |
| --- | --- | --- | --- | --- |
| Adult Anseriformes | 0 |  |  |  |
| Adult Galliformes | **0.022521129†** | 0 |  |  |
| Anseriformes Hatchlings | **0.0290663†** | 0.006545176 | 0 |  |
| Galliformes Hatchlings | **0.01586763†** | 0.006653498 | **0.013198673*** | 0 |

**Table S4: Results of statistical tests comparing the allometric ontogeny of crown Galliformes versus Anseriformes; and Anatidae versus Phasianidae.**

| **Test** | **Difference (d)** | **P-value** |
| --- | --- | --- |
| **Anseriformes vs. Galliformes** | **Difference (d)** | **p-value** |
| Test of Homogeneity of Slopes (HOS) | 0.000809099 | 0.0817 |
| Test of Homogeneity of Intercept | 0.01262752 | 0.4678532 |
| Peramorphosis test | 0.2914082 | **0.0004** |
| **Anatidae vs. Phasianidae** |  |  |
| Test of Homogeneity of Slopes (HOS) | 0.000799563 | 0.9846 |
| Test of Homogeneity of Intercept | 0.00926048 | 0.5914409 |
| Peramorphosis test | 0.3072193 | **0.0006** |

**Table S5: Clade-specific values of some ontogenetic variables distinguishing crown galliform and anseriform cranial ontogenies.**

| **Variable** | **Anseriformes** | **Galliformes** |
| --- | --- | --- |
| Mean hatchling skull centroid size | 0.17 | 0.13 |
| Mean adult skull centroid size | 0.43 | 0.28 |
| Mean amount of ontogenetic shape change | 0.193 | 0.16 |
| Mean amount of ontogenetic size change | 0.393 | 0.313 |

**Table S6: Interclade differences, between crown Galliformes and Anseriformes, in some cranial ontogenetic variables.**

| **Variable** | **Amount of difference** | **p-value** |
| --- | --- | --- |
| Mean hatchling skulls centroid size | 0.037 | **0.0006** |
| Mean adult skulls centroid size | 0.15 | **0.0004** |
| Mean hatchling skulls shape | 0.18 | **0.0004** |
| Mean adult skulls shape | 0.25 | **0.0004** |
| Mean ontogenetic shape change | 0.03 | **0.0016** |
| Mean ontogenetic size change | 0.08 | **0.0026** |

**Table S7: The log-likelihood of the fit of four evolutionary models to the adult and hatchling skull shapes evaluated using the generalised information criterion.** The model with the highest log-likelihood is shown is highlighted in grey.

|  | **Brownian Motion** | **Ornstein-Uhlenbeck** | **Early Burst** | **Lambda** |
| --- | --- | --- | --- | --- |
| **Adult shapes** | 47735.12 | 48151.05 | 47735.11 | 48545.62 |
| **Hatchling shapes** | 48402.72 | 48887.48 | 48402.72 | 49391.01 |

**Table S8: List of taxa and specimens used for the investigation of ontogenetic cranial disparity**

| **Family** | **Clade** | **Name** | **Age** | **Accession code** |
| --- | --- | --- | --- | --- |
| Anseriformes | Anatidae | *Aix sponsa* | Hatchling | DJF_frozen_1 |
|  |  | *Aix galericulata* | Adult | DJF_frozen_2 |
|  |  | *Alopochen aegyptiaca* | Hatchling | DJF_frozen_3 |
|  |  |  | Adult | NHMUK 1932.11.13.1 |
|  |  | *Aythya affinis* | Near-hatching | DJF_frozen_4 |
|  |  |  | Adult | NHMUK_s.1986.60.6 |
|  |  | *Mergellus albellus* | Hatchling | DJF_frozen_5 |
|  |  |  | Adult | DJF_frozen_6 |
|  |  | *Somateria fischeri* | Hatchling | DJF_frozen_7 |
|  |  |  | Adult | DJF_frozen_8 |
|  |  | *Callonetta leucophrys* | Hatchling | DJF_frozen_9 |
|  |  |  | Adult | DJF_frozen_10 |
|  |  | *Anas platyrhynchos* | Hatchling | OUVC10613 |
|  |  |  | Adult | UMZC 225 |
|  |  | *Oxyura maccoa* | Hatchling | DJF_frozen_11 |
|  |  | *Oxyura jamaicensis* | Adult | NHMUK S 1985.70.4 |
|  |  | *Anser cygnoides* | Hatchling | AD_hatch_R1 |
|  |  | *Anser anser* | Adult | USN_291065 |
|  |  | *Cygnus melancoryphus* | Hatchling | DJF_frozen_12 |
|  |  | *Cygnus olor* | Adult | NHMUK 1854.6.25.2 |
|  |  | *Malacorhynchus membranaceus* | Hatchling (1 Day) | DJF_frozen_13 |
|  |  |  | Chick | DJF_frozen_14 |
|  |  |  | Adult | UMZC 12-Ana-33-a-1 |
|  |  | *Dendrocygna guttata* | Hatchling (1 Day) | DJF_frozen_15 |
|  |  |  | Adult | DJF_frozen_16 |
|  |  | *Tadorna tadorna* | Chick | BMNH_No_Reg |
|  |  |  | Adult | BMNH.S-2004.13.1 |
|  |  | *Mergus serrator* | Chick | BMNH_No_Reg |
|  |  |  | Adult | DJF_frozen_17 |
|  |  | *Spatula clypeata* | Chick | BMNH_1929.6.15.5 |
|  |  |  | Adult | BMNH_A/1970.21.4 |
|  | Anseranatidae | *Anseranas semipalmata* | Hatchling | O2377 |
|  |  |  | Adult | NHMUK1852.7.22.1 |
|  | Anhimidae | *Chauna chavaria* | Hatchling | BMNH_ 1923.5.24.30 |
|  |  |  | Adult | OUMNH23790 |
| Galliformes | Phasianidae | *Pavo cristatus* | Chick | BMNH.1930.1.6.2 |
|  |  |  | Adult | UMZC 396.D |
|  |  | *Afropavo congensis* | Near-hatching | DJF_frozen_18 |
|  |  |  | Adult | NHMUK S/1989.19.16 |
|  |  | *Gallus domesticus* | Hatchling | GG46R4 |
|  |  |  | Adult | UMZC_400 |
|  |  | *Polyplectron bicalcaratum* | Hatchling | BMNH.A-2011.9.13 |
|  |  | *Polyplectron chalcurum* | Adult | BMNH.1931.10.5.5 |
|  |  | *Coturnix coturnix* | Hatchling | CJ45R4 |
|  |  | *Coturnix domesticus* | Adult | UMMZ_224005 |
|  |  | *Lagopus lagopus* | Hatchling | BMNH.1925.11.6.3 |
|  |  |  | Adult | UMZC_uncatalogued_frozen18 |
|  |  | *Meleagris gallopavo* | Hatchling (up to 7 days) | OUVC 12483 |
|  |  |  | Adult | BMNH.S-2017.40.83 |
|  |  | *Perdix perdix* | Hatchling | BMNH.1931.6.8.10 |
|  |  |  | Adult | UMZC_uncatalogued_frozen21 |
|  |  | *Tragopan satyra* | Hatchling | BMNH.A-2013.3.11 |
|  |  |  | Adult | NHMUK S 2010.1.36 |
|  |  | *Rollulus rouloul* | Near-hatching | DJF_frozen_19 |
|  |  |  | Adult | NHMUK 1871.7.20.87 |
|  | Odontophoridae | *Callipepla californica* | Hatchling | BMNH.A.1985.17.22 |
|  |  |  | Adult | BMNH.A.1985.17.22 |
|  | Numididae | *Numida meleagris* | Near-hatching | DJF_frozen_20 |
|  |  |  | Adult | UMZC 393 |
|  | Cracidae | *Ortalis ruficauda* | Chick | BMNH_No_Reg |
|  |  | *Ortalis spp.* | Adult | UMMZ 155489 |
|  | Megapodiidae | *Megapodius pritchardii* | Hatchling | UMZC 14/Meg/6/g/1 |
|  |  |  | Adult | BMNH.1940.12.8.133 |
| Columbiformes | Columbidae | *Columba livia* | Chick | DJF_frozen_21 |
|  |  |  | Adult | FMNH_347273 |
| Tinamiformes | Tinamidae | *Crypturellus tataupa* | Near-hatching | DJF_frozen_22 |
|  |  |  | Adult | UMZC uncatalogued |
| Struthioniformes | Struthionidae | *Struthio camelus* | Near-hatching | TLG SC047 |
|  |  |  | Adult | TLG SC080 |
| Pan-Anseriformes | Presbyornithidae | *Presbyornis* ^†^ | Adult | UMNH.VP.29023.LLU1101 |
|  |  |  | Adult | USNM-299846 |
| Pan-Galloanserae | - | *Asteriornis* ^†^ | Adult | NHMM 2013 008.1 |

**Table S9: Landmarks used for analysis of cranial ontogenetic disparity.** All the landmarks listed were applied for collection 1 specimens. The reduced landmark configuration relied on the landmarks highlighted in grey.

| **Landmark Number** | **Abb.** | **Description** |
| --- | --- | --- |
| 1 | BT | Beak Tip |
| 2 | RTLN | Rostral Tip of left nostril |
| 3 | RTRN | Rostral Tip of right nostril |
| 4 | CTLN | Caudal Tip of left nostril |
| 5 | CTRN | Caudal Tip of right nostril |
| 6 | CTLM | Caudal Tip of left side of the beak |
| 7 | CTRM | Caudal Tip of right side of the beak |
| 8 | RLLFC | The rostral tip of the left lacrimal-frontal contact |
| 9 | RRLFC | The rostral tip of the right lacrimal-frontal contact |
| 10 | LLP | lateral edge of Left palatal-maxillary contact |
| 11 | LRP | lateral edge of right palatal-maxillary contact |
| 12 | MLP | Medial edge of Left palatal-maxillary contact |
| 13 | MRP | Medial edge of right palatal-maxillary contact |
| 14 | RIS | Rostral end of interfrontal suture |
| 15 | CLLFC | The caudal tip of the left lacrimal-frontal contact |
| 16 | CRLFC | The caudal tip of the right lacrimal-frontal contact |
| 17 | CLC | Caudal point left palatal choana |
| 18 | CRC | Caudal point right palatal choana |
| 19 | DLPPC | The dorsomedial tip of the left palatine-pterygoid contact |
| 20 | DRPPC | The dorsomedial tip of the right palatine-pterygoid contact |
| 21 | RLLP | The dorsolateral tip of the left pterygoid-palatine contact |
| 22 | RLRP | The dorsolateral tip of the right pterygoid-palatine contact |
| 23 | VLPPC | The ventromedial tip of the left palatine-pterygoid contact |
| 24 | VRPPC | The ventromedial tip of the right palatine-pterygoid contact |
| 25 | LPP | Tip of left postorbital process |
| 26 | RPP | Tip of Right postorbital process |
| 27 | RLS | The rostroventral tip of the left squamosal |
| 28 | RRS | The rostroventral tip of the right squamosal |
| 29 | COLQ | Tip of the crista orbitalis of the left quadrate |
| 30 | CORQ | Tip of the crista orbitalis of the right quadrate |
| 31 | LLPQC | Ventrolateral edge of the left pterygoid-quadrate contact |
| 32 | LRPQC | Ventrolateral edge of the right pterygoid-quadrate contact |
| 33 | MLPQC | Ventromedial edge of the left pterygoid-quadrate contact |
| 34 | MRPQC | Ventromedial edge of the right pterygoid-quadrate contact |
| 35 | MTBP | Medial tip of the basiparasphenoid |
| 36 | RTLZP | Rostral Tip of the left zygomatic process of the squamosal |
| 37 | RTRZP | Rostral Tip of the right zygomatic process of the squamosal |
| 38 | DLCLQ | Dorsal end of the lateral crest of the left quadrate |
| 39 | DLCRQ | Dorsal end of the lateral crest of the right quadrate |
| 40 | ROLQ | Rostral edge of the otic process of the left quardate |
| 41 | RORQ | Rostral edge of the otic process of the right quardate |
| 42 | LMCLQ | Buttress of the lateral mandibular condyle of the left quadrate |
| 43 | LMCRQ | Buttress of the lateral mandibular condyle of the right quadrate |
| 44 | CLQ | Caudal end of the left quadrate-quadratojugal contact |
| 45 | CRQ | Caudal end of the right quadrate-quadratojugal contact |
| 46 | DFM | Dorsal tip of foramen magnum |
| 47 | COC | Caudal end of occipital condyle |
| 48 | CLPP | caudolateral tip of the Left process paraoccipitalis |
| 49 | CRPP | caudolateral tip of the right process paraoccipitalis |

**Table S10: Semilandmarks used for analysis of cranial ontogenetic disparity.** All the semilandmarks listed were applied for collection 1 specimens. The reduced landmark configuration relied on the semilandmarks highlighted in grey.

| **Curve #** | **Description** | **Number of semi-landmarks** | **Origin (Landmark #)** | **End (Landmark #)** |
| --- | --- | --- | --- | --- |
| C1 | Lateral rim left nostril | 5 | 2 | 4 |
| C2 | Lateral rim right nostril | 5 | 3 | 5 |
| C3 | Medial rim left nostril | 5 | 2 | 4 |
| C4 | Medial rim right nostril | 5 | 3 | 5 |
| C5 | Tip of the beak to the caudal tip of the frontonasal hinge | 10 | 1 | 14 |
| C6 | The left lateral rim of the beak | 10 | 1 | 6 |
| C7 | The right lateral rim of the beak | 10 | 1 | 7 |
| C8 | The left caudal rim of the beak | 6 | 8 | 6 |
| C9 | The right caudal rim of the beak | 6 | 9 | 7 |
| C10 | The ventrolateral rim of the left lacrimal | 14 | 8 | 15 |
| C11 | The ventrolateral rim of the right lacrimal | 14 | 9 | 16 |
| C12 | The dorsal rim of the left orbit | 12 | 15 | 25 |
| C13 | The dorsal rim of the right orbit | 12 | 16 | 26 |
| C14 | Ventral rim of the left postorbital process extending from its tip to the tip of the zygomatic process | 10 | 27 | 36 |
| C15 | Ventral rim of the right postorbital process extending from its tip to the tip of the zygomatic process | 10 | 28 | 37 |
| C16 | From the medial edge of the left pterygoid-palatal contact through the lateral edge of the left palatine to its points of fusion with the maxillary | 12 | 23 | 10 |
| C17 | From the medial edge of the right pterygoid-palatal contact through the lateral edge of the right palatine to its points of fusion with the maxillary | 12 | 24 | 11 |
| C18 | The medial edge of the left palatine | 10 | 17 | 12 |
| C19 | The medial edge of the right palatine | 10 | 18 | 13 |
| C20 | The dorsal rim of the left pterygoid | 12 | 31 | 19 |
| C21 | The dorsal rim of the right pterygoid | 12 | 32 | 20 |
| C22 | The caudolateral rim of the left pterygoid | 6 | 31 | 21 |
| C23 | The caudolateral rim of the right pterygoid | 6 | 32 | 22 |
| C24 | The caudomedial rim of the left pterygoid | 10 | 33 | 23 |
| C25 | The caudomedial rim of the right pterygoid | 10 | 34 | 24 |
| C26 | Along the margin of the left quadrate connecting the crista orbitalis, the crista orbitocotylaris and the buttress of the lateral mandibular condyle | 7 | 29 | 42 |
| C27 | Along the margin of the right quadrate connecting the crista orbitalis, the crista orbitocotylaris and the buttress of the lateral mandibular condyle | 7 | 30 | 43 |
| C28 | Along the dorsal rim of the left quadrate connecting the crista orbitalis to the rostrodorsal edge of the otic process | 6 | 40 | 29 |
| C29 | Along the dorsal rim of the right quadrate connecting the crista orbitalis to the rostrodorsal edge of the otic process | 6 | 41 | 30 |
| C30 | Along the lateral crest of the left quadrate | 6 | 38 | 44 |
| C31 | Along the lateral crest of the right quadrate | 6 | 39 | 45 |

**Table S11: Homology criteria used for analysis of ontogenetic shape change.**

|  | **Adult** | **Hatchling** |
| --- | --- | --- |
| 1 | Beak Tip | Beak Tip |
| 2 | Rostral Tip of left nostril | Rostral Tip of left nostril |
| 3 | Rostral Tip of right nostril | Rostral Tip of right nostril |
| 4 | Caudal Tip of left nostril | Caudal Tip of left nostril |
| 5 | Caudal Tip of right nostril | Caudal Tip of right nostril |
| 6 | Caudal Tip of left side of the beak | Caudal Tip of left side of the beak |
| 7 | Caudal Tip of right side of the beak | Caudal Tip of right side of the beak |
| 8 | The rostral tip of the left lacrimal-frontal contact | The rostral tip of the left lacrimal closes to its contact with the frontal |
| 9 | The rostral tip of the right lacrimal-frontal contact | The rostral tip of the right lacrimal closes to its contact with the frontal |
| 10 | lateral edge of Left palatal-maxillary contact | The region on the surface of the left palatine that is directly medial to the caudolateral end of the palatal process of the left maxillary |
| 11 | lateral edge of right palatal-maxillary contact | The region on the surface of the right palatine that is directly medial to the caudolateral end of the palatal process of the right maxillary |
| 12 | Medial edge of Left palatal-maxillary contact | The region on the surface of the left palatine that is nearest to the caudolateral edge of the palatal process of the left maxillary |
| 13 | Medial edge of right palatal-maxillary contact | The region on the surface of the left palatine that is nearest to the caudolateral edge of the palatal process of the left maxillary |
| 14 | Rostral end of interfrontal suture | Rostral end of interfrontal suture |
| 15 | The caudal tip of the left lacrimal-frontal contact | The caudal tip of the left lacrimal closes to its contact with the frontal |
| 16 | The caudal tip of the right lacrimal-frontal contact | The caudal tip of the right lacrimal closes to its contact with the frontal |
| 17 | Caudal point left palatal choana | Caudal point left palatal choana |
| 18 | Caudal point right palatal choana | Caudal point right palatal choana |
| 19 | The dorsomedial tip of the left palatine-pterygoid contact | The dorsomedial tip of the left palatine-pterygoid contact |
| 20 | The dorsomedial tip of the right palatine-pterygoid contact | The dorsomedial tip of the right palatine-pterygoid contact |
| 21 | The dorsolateral tip of the left pterygoid-palatine contact | The dorsolateral tip of the left pterygoid-palatine contact |
| 22 | The dorsolateral tip of the right pterygoid-palatine contact | The dorsolateral tip of the right pterygoid-palatine contact |
| 23 | The ventromedial tip of the left palatine-pterygoid contact | The ventromedial tip of the left palatine-pterygoid contact |
| 24 | The ventromedial tip of the right palatine-pterygoid contact | The ventromedial tip of the right palatine-pterygoid contact |
| 25 | Tip of left postorbital process | The rostrolateral tip of the left laterosphenoid |
| 26 | Tip of Right postorbital process | The rostrolateral tip of the right laterosphenoid |
| 27 | The rostroventral tip of the left squamosal | The rostroventral tip of the left squamosal |
| 28 | The rostroventral tip of the right squamosal | The rostroventral tip of the right squamosal |
| 29 | Tip of the crista orbitalis of the left quadrate | The Rostrolateral edge of the orbital process of the left quadrate |
| 30 | Tip of the crista orbitalis of the right quadrate | The Rostrolateral edge of the orbital process of the right quadrate |
| 31 | Ventrolateral edge of the left pterygoid-quadrate contact | Ventrolateral edge of the left pterygoid-quadrate contact |
| 32 | Ventrolateral edge of the right pterygoid-quadrate contact | Ventrolateral edge of the right pterygoid-quadrate contact |
| 33 | Ventromedial edge of the left pterygoid-quadrate contact | Ventromedial edge of the left pterygoid-quadrate contact |
| 34 | Ventromedial edge of the right pterygoid-quadrate contact | Ventromedial edge of the right pterygoid-quadrate contact |
| 35 | Medial tip of the basiparasphenoid | Medial tip of the basiparasphenoid |
| 36 | Rostral Tip of the left zygomatic process of the squamosal | Rostral Tip of the left zygomatic process of the squamosal |
| 37 | Rostral Tip of the right zygomatic process of the squamosal | Rostral Tip of the right zygomatic process of the squamosal |
| 38 | Dorsal end of the lateral crest of the left quadrate | Dorsal end of the lateral crest of the left quadrate |
| 39 | Dorsal end of the lateral crest of the right quadrate | Dorsal end of the lateral crest of the right quadrate |
| 40 | Rostral edge of the otic process of the left quardate | Rostral edge of the otic process of the left quardate |
| 41 | Rostral edge of the otic process of the right quardate | Rostral edge of the otic process of the right quardate |
| 42 | Buttress of the lateral mandibular condyle of the left quadrate | Buttress of the lateral mandibular condyle of the left quadrate |
| 43 | Buttress of the lateral mandibular condyle of the right quadrate | Buttress of the lateral mandibular condyle of the right quadrate |
| 44 | Caudal end of the left quadrate-quadratojugal contact | Caudal end of the lateral crest of the left Quadrate |
| 45 | Caudal end of the right quadrate-quadratojugal contact | Caudal end of the lateral crest of the right Quadrate |
| 46 | Dorsal tip of foramen magnum | Dorsal tip of foramen magnum |
| 47 | Caudal end of occipital condyle | The caudomedial point of the basioccipital |
| 48 | caudolateral tip of the Left process paraoccipitalis | The ventrocaudal tip of the left squamosal |
| 49 | caudolateral tip of the right process paraoccipitalis | The ventrocaudal tip of the right squamosal |

### Supplementary Figures

**Supplementary Figure 1: The cranial disparity of crown Gallaonserae at adult and hatchling stages.** Adult anseriform disparity is the highest among all groups, while galliform disparity is similar at hatchling and adult stages.

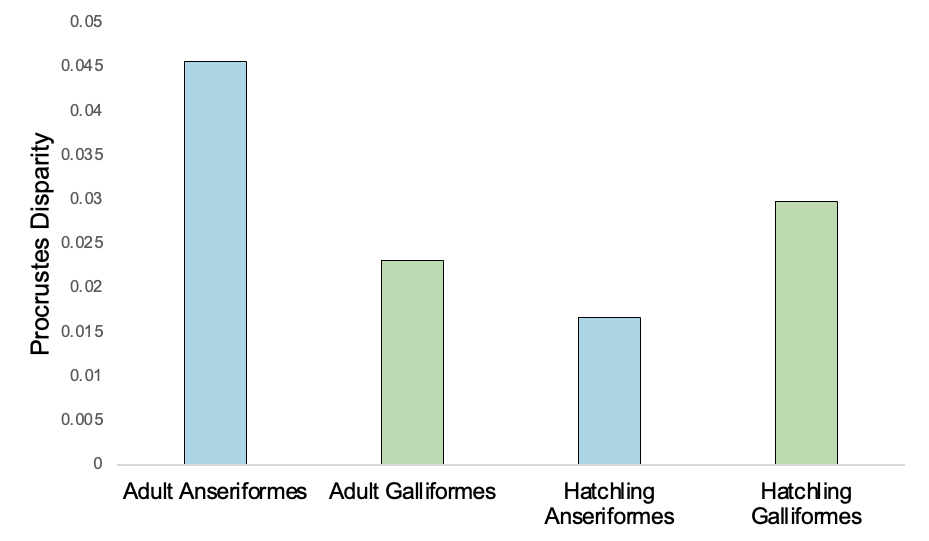

*

*

Procrustes Disparity

|  | Procrustes Disparity |
| --- | --- |
| Adult Anseriformes | 0.0456883 |
| Adult Galliformes | 0.0231671 |
| Hatchling Anseriformes | 0.016622 |
| Hatchling Galliformes | 0.0298206 |

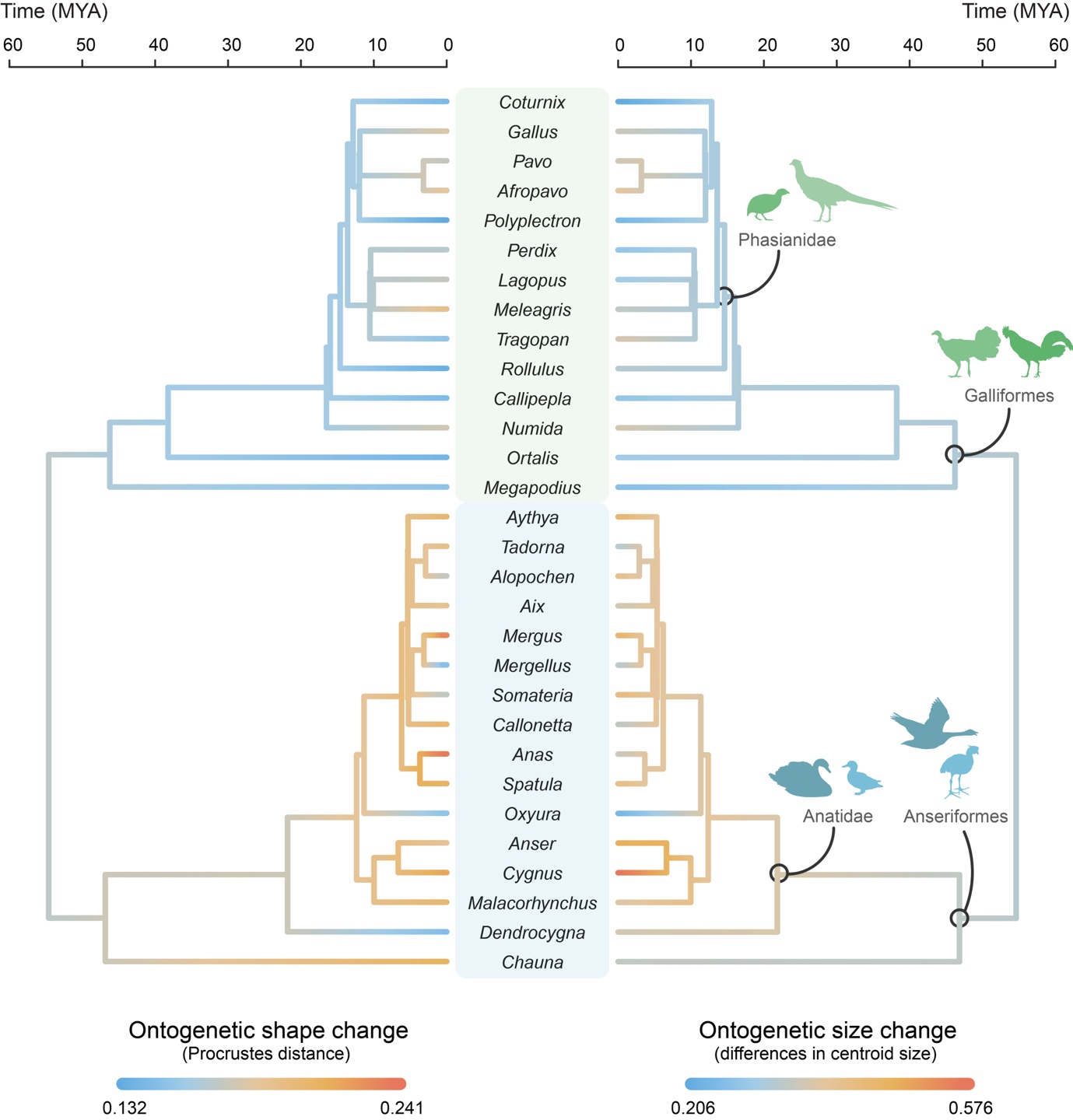
**Supplementary Figure 2: Variation and evolutionary history of the amount of ontogenetic size and shape change in the cranial ontogeny of Galloanserae.** Anseriformes exhibit a greater and more variable degree of ontogenetic change than Galliformes.

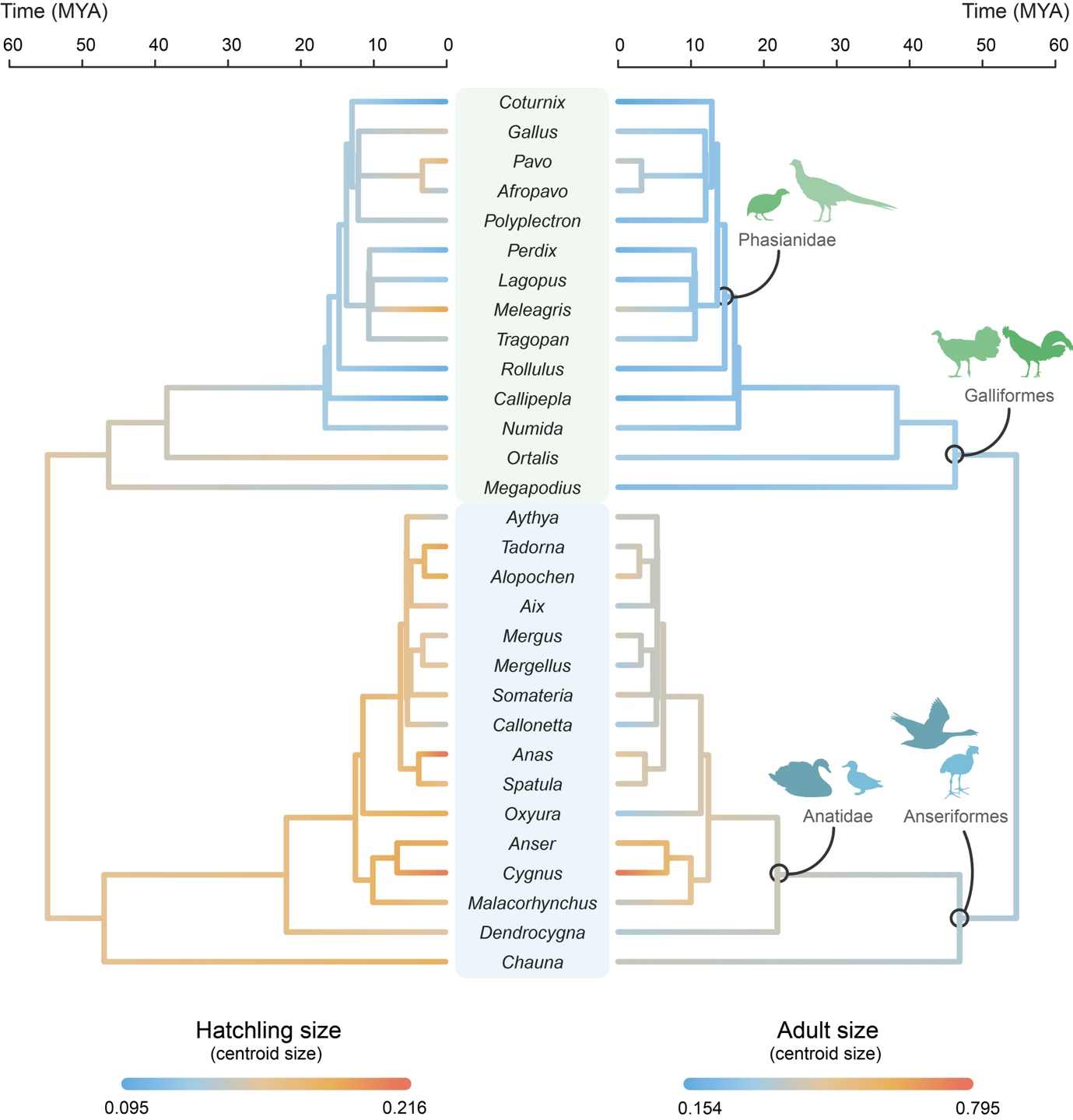
**Supplementary Figure 3: Variation and evolutionary history of Galloanserae hatchling and adult skull sizes.** Anseriformes exhibit larger skull sizes than Galliformes in general.

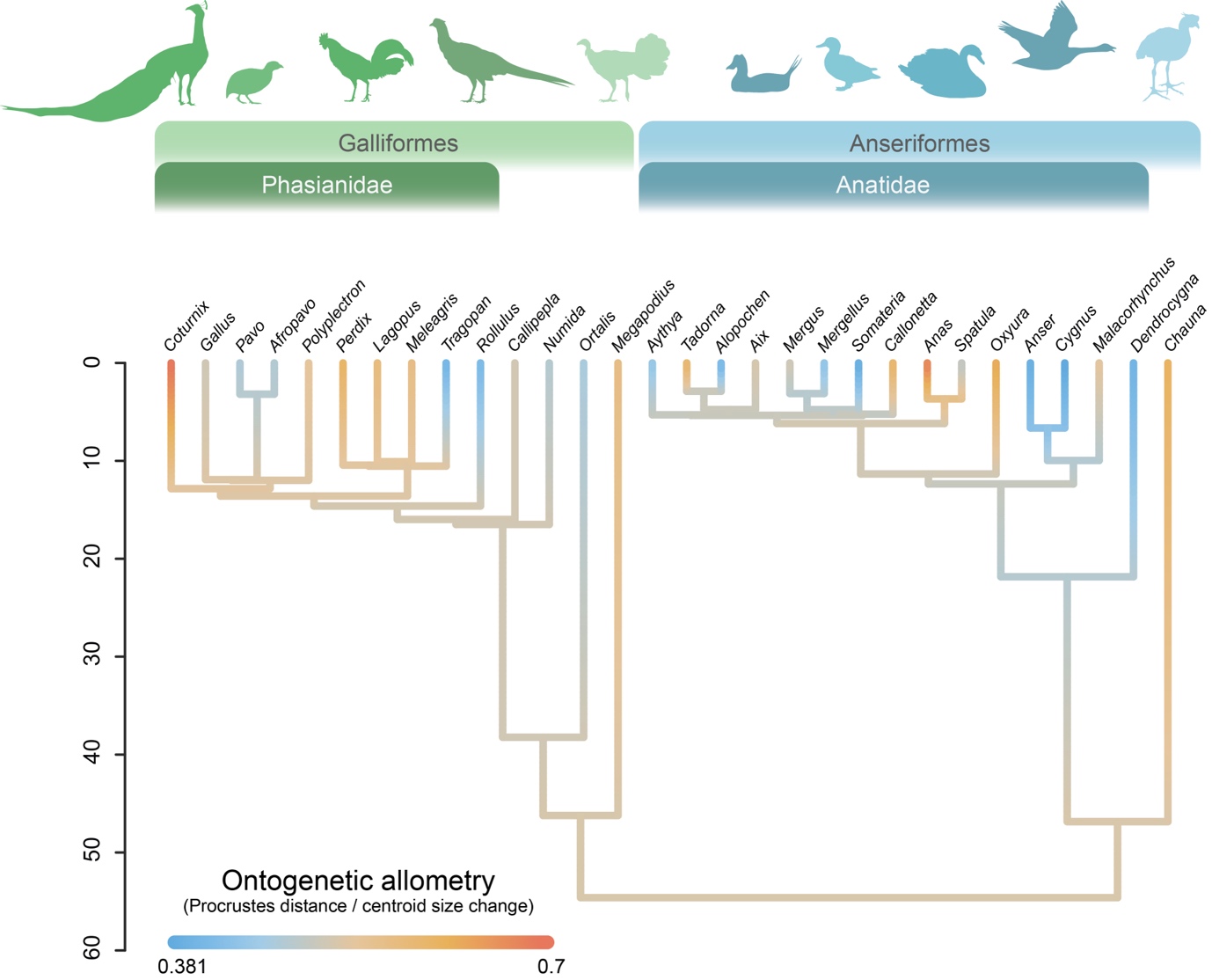
**Supplementary Figure 4: Variation and evolutionary history of rate of cranial allometric shape change in Galloanserae.** The inter- and intraclade variance in ontogenetic allometry is roughly equal between Anseriformes and Galliformes.

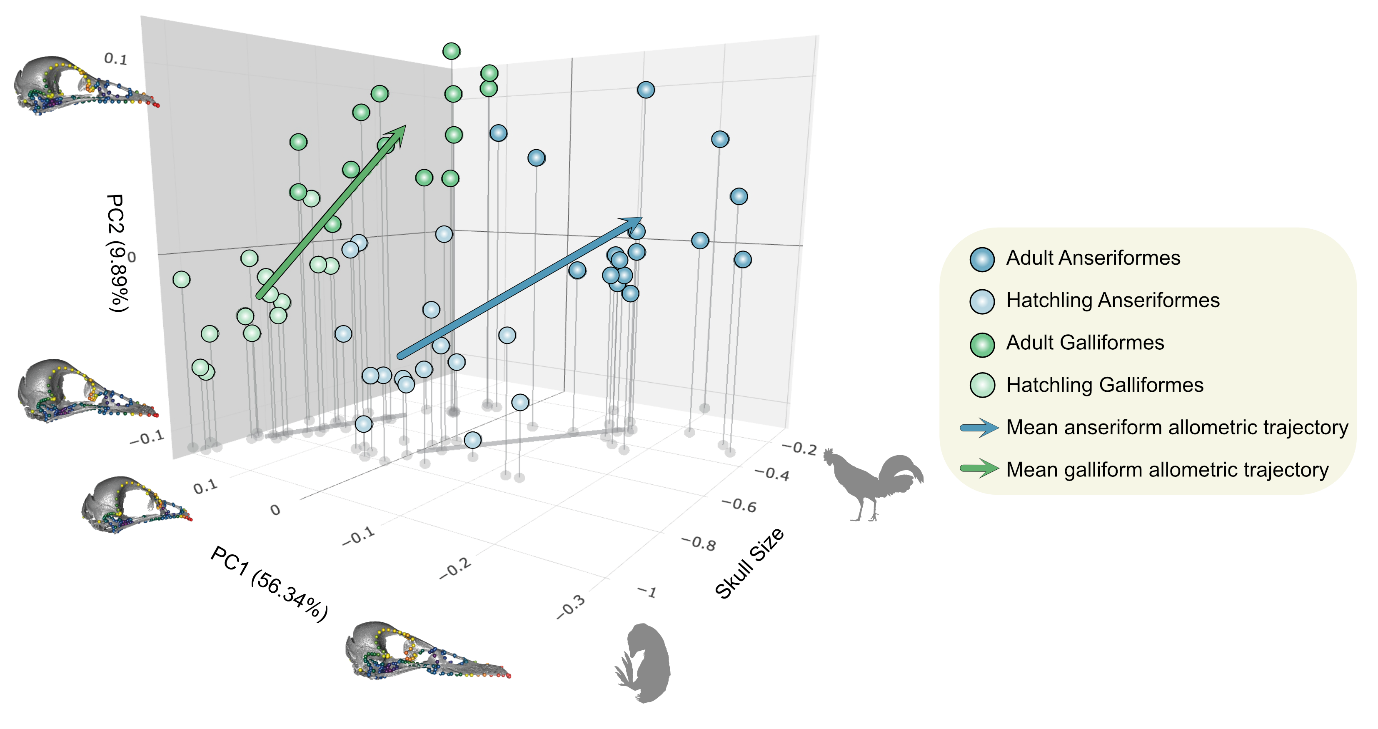
**Supplementary Figure 5: Crown Galloanserae cranial allometric space.** The trajectories were deduced by regressing shape and sizes of hatchling and adult skulls, and they display some phylogenetic clustering. The skull shapes associated with regression score variance are shown on the axis.

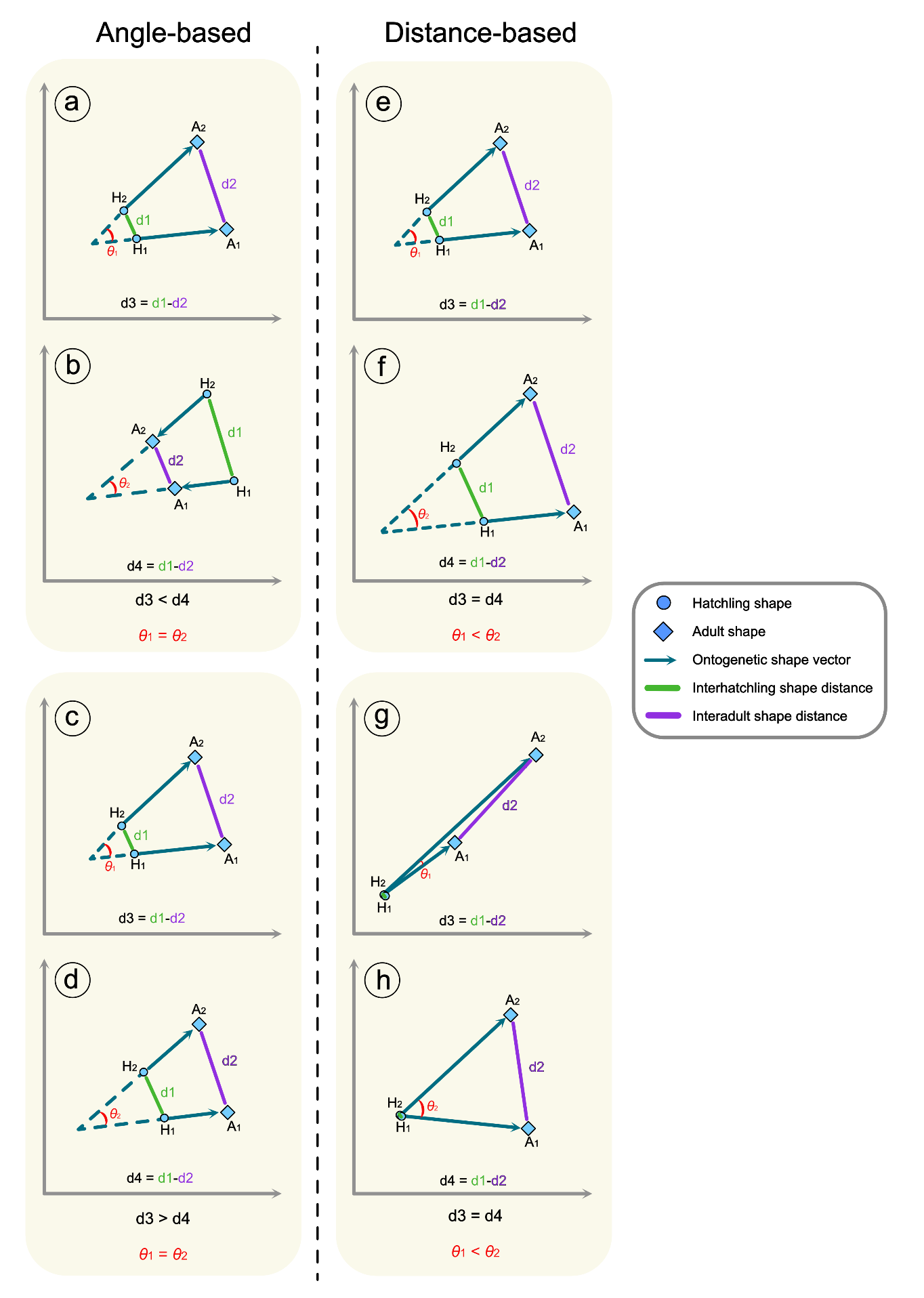
**Supplementary Figure 6: Limitations of the methods used for determining ontogenetic convergence/divergence.** Left) limitations of the angle-based method: a-b) Misperception of convergence and divergence. Convergent and divergent ontogenies are indistinguishable based the angular distance between interadult shape distance. They can only be distinguished by the relative interhatchling and interadult shape distances. c-d) Overestimation of divergence/convergence. The angular distance between adult shapes is the same regardless of the interhatchling shape distance, which. The distance-based method considers the interhatchling shape distance. Right) limitations of the distance-based method: e-f) Underestimation of divergence/convergence. Trajectories with the same interhatchling shape distance/interadult shape distance ratio have the same amount of divergence, regardless of the magnitude of the interadult shape distance. They are more accurately distinguished by the angular distance between them. g-h) Obfuscation of type of convergence/divergence. Pairs of trajectories with similar directions but with unequal lengths appear as divergent, or convergent, as pairs with equal trajectory length and different directions. The distunction between these two forms of divergence is discernible with the angular distances between them. Abbreviations: d1; the interhatchling shape distance, d2; interadult shape distance, $\theta$; the angle between two ontogenetic trajectories.

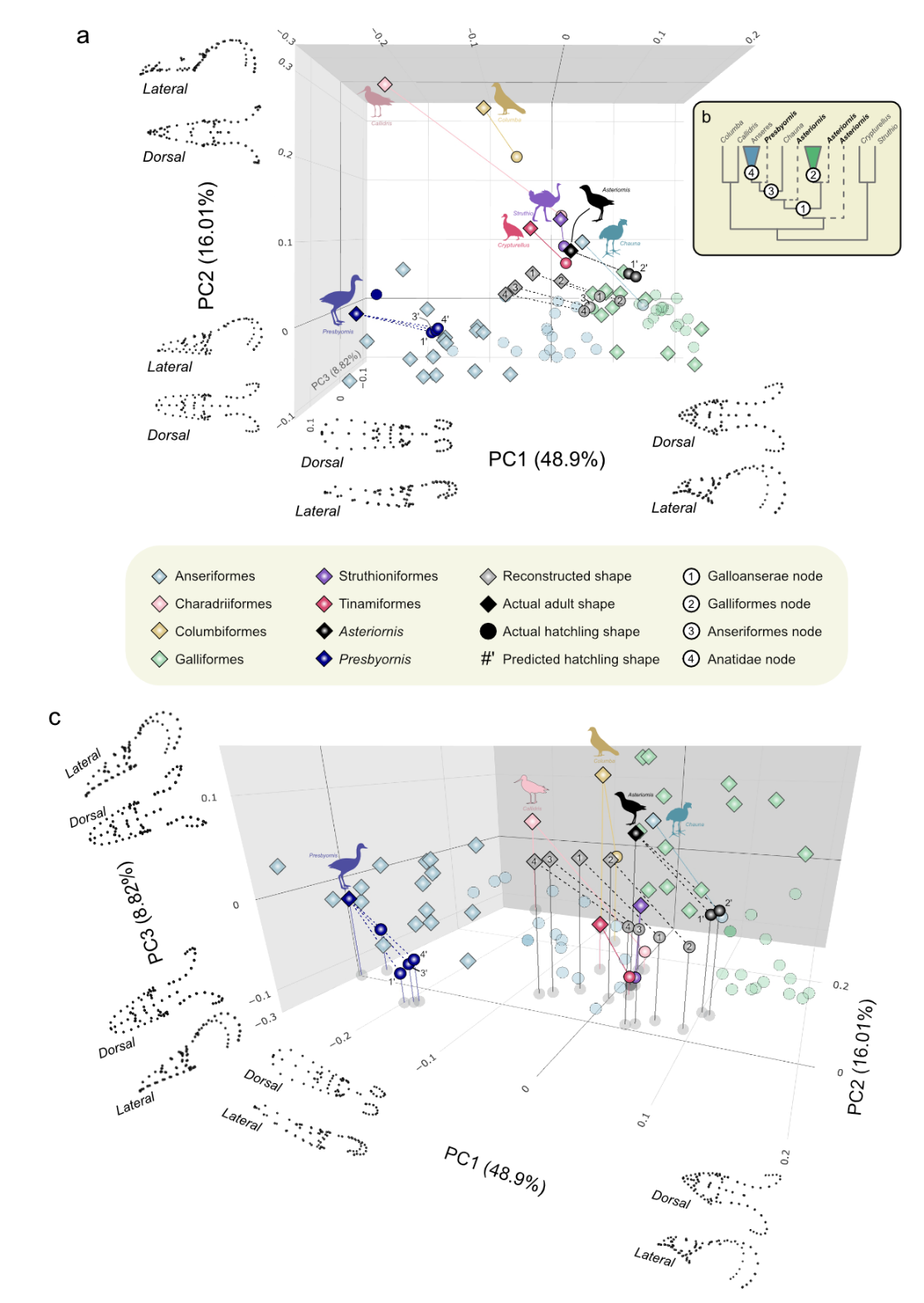

**Supplementary Figure 7: Cranial post-hatching ontogenetic morphospace of crown and stem Galloanserae along with some outgroups.** a) A 3D plot of the ontogenetic cranial disparity of crown Galloanserae showing phylogenetic clustering, except for *Chuana*, between Galliformes and Anseriformes. The reconstructed ancestral cranial ontogenies appeared between the two clusters of crown species, while *Presbyornis* ontogeny clustered near anatid species and *Asteriornis* ontogeny appeared near galliform species. The proximity of the skull shapes of *Chauna*, *Asteriornis* hatchling, and the most recent common ancestor of Galloanserae to the cluster of extnat Galliformes indicates the plesiomorphic nature of galliform cranial morphology. b) Simplified phylogenetic tree of the clades used in this study, with the specific nodes, that were used in ancestor reconstruction, are numbered. The phylogenetic position of *Asteriornis* is uncertain, so it was placed in three different candidate positions. c) Another view of (a) showing that the reconstructed ancestral gallaonseran cranial ontogenies is between that of crown Galliformes and Anseriformes.

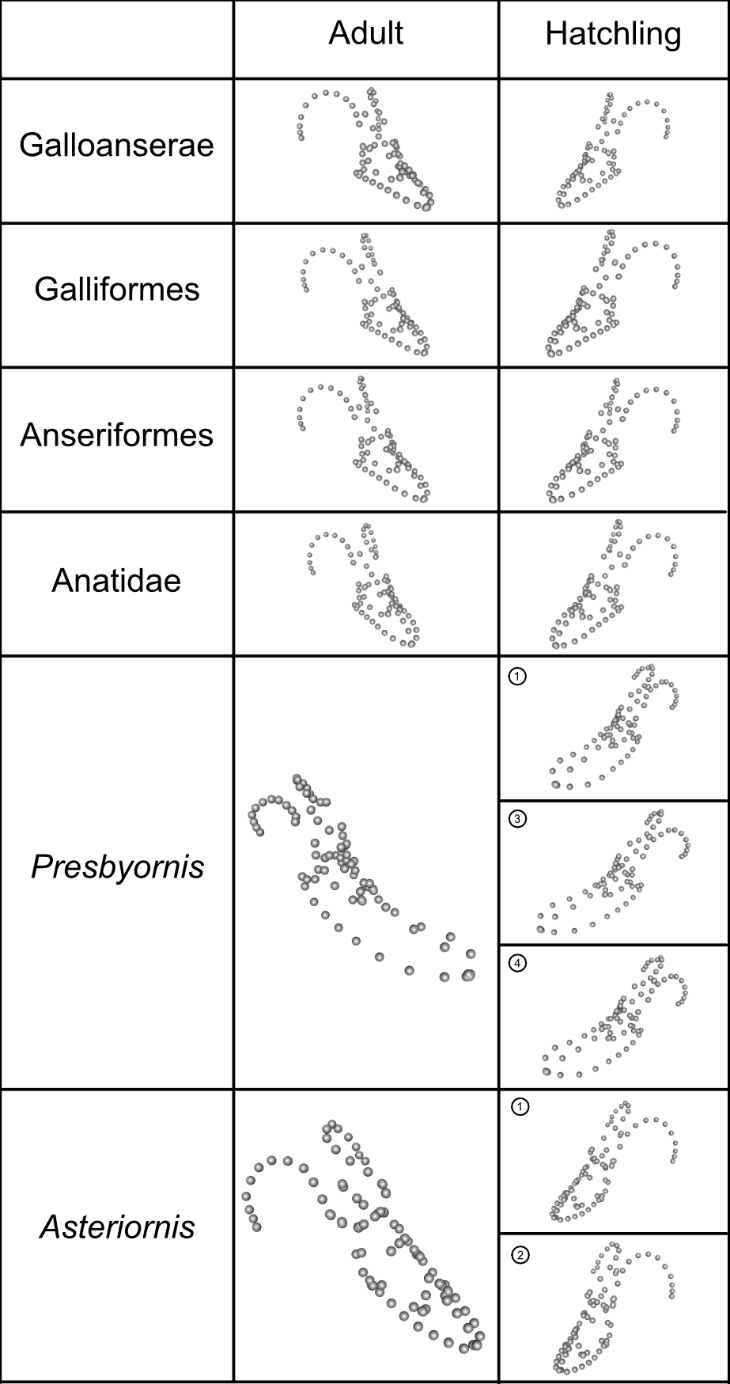

**Supplementary Figure 8: The reconstructed shapes of ancestral galloanseran skulls along with the predicted shapes of fossil hatchlings.** The reconstructed skull shapes of adults and hatching of the most recent common ancestor of each group is shown in the top four rows. The shape of adult *Asteriornis* is the landmark configuration applied to the reconstructed fossil, and the shape of adult *Presbyornis* is the landmark configuration applied to the segmented fossil skull. The shapes of the fossil hatchlings were predicted based on the ontogenies at the base of ① Galloanserae, ② Galliformes, ③ Anseriformes, and ④ Anatidae.
