## Supplementary material for "Macroevolutionary shifts in post-hatching ontogeny and the origin of craniofacial disparity in fowl (Aves: Galloanserae)": Code used for the analysis

```
# This code only includes the analyses for Arnaout et al., 202X

library(geomorph)
library(RRPP)
library(rgl)
library(tidyverse)
library(Morpho)
library(hot.dots)
library(tibble)
library(gdata)
library(dplyr)
library(gMOIP)
library(shapes)
library(ggplot2)
library(plotly)
library(morphospace)
library(Momocs)
library(magrittr)
library(reshape2)
library(viridis)
library(dplyr)
library(matlib)
library(phytools)
library(ape)
library(abind)
library(phytools)
library(ape)
library(devtools)
library(mvMORPH)
library("PlotTools")
library("dendextend")
library("wesanderson") ## Color
library("RColorBrewer")## Color


############Landmark Data Importation############
####Collection One
### Extant Galloanserae Landmarks
duck <- read.delim("Landmarks/Anas Platyrhynchos adult/
AllLandmarks.txt", header=TRUE)
duckling <- read.delim("Landmarks/Anas Platyrhynchos hatchling/
AllLandmarks.txt", header=TRUE)
chicken <- read.delim("Landmarks/Gallus domesticus adult/
AllLandmarks.txt", header=TRUE)
chick <- read.delim("Landmarks/Gallus domesticus hatchling/
AllLandmarks.txt", header=TRUE)
WDuck <- read.delim("Landmarks/Dendrocygna guttata adult/
```

```
AllLandmarks.txt", header=TRUE)
WDuckling <- read.delim("Landmarks/Dendrocygna guttata
hatchling/AllLandmarks.txt", header=TRUE)
AfropavoHatchling <- read.delim("Landmarks/Afropavo congensis
hatchling/AllLandmarks.txt", header=TRUE)
Afropavo <- read.delim("Landmarks/Afropavo congensis adult/
AllLandmarks.txt", header=TRUE)
Pavoling<- read.delim("Landmarks/Pavo cristatus hatchling/
AllLandmarks.txt", header=TRUE)
PavoAdult <- read.delim("Landmarks/Pavo cristatus adult/
AllLandmarks.txt", header=TRUE)
AythyaHatchling <- read.delim("Landmarks/Aythya affinis
hatchling/AllLandmarks.txt", header=TRUE)
AythyaAdult <- read.delim("Landmarks/Aythya ferina adult/
AllLandmarks.txt", header=TRUE)
megapodling <- read.delim("Landmarks/Megapodius pritchardii
hatchling/AllLandmarks.txt", header=TRUE)
megapodadult <- read.delim("Landmarks/Megapodius pritchardii
adult/AllLandmarks.txt", header=TRUE)
swanling <- read.delim("Landmarks/Cygnus melancoryphus
hatchling/AllLandmarks.txt", header=TRUE)
swanadult <- read.delim("Landmarks/Cygnus olor/
AllLandmarks.txt", header=TRUE)
Numidling <- read.delim("Landmarks/Numida meleagris embryo/
AllLandmarks.txt", header=TRUE)
NumidiaAdult <- read.delim("Landmarks/Numida meleagris adult/
AllLandmarks.txt", header=TRUE)
PinkDuckling <- read.delim("Landmarks/Malacorhynchus
membranaceus hatchling/AllLandmarks.txt", header=TRUE)
PinkDuckling2 <- read.delim("Landmarks/Malacorhynchus
membranaceus hatchling 2/AllLandmarks.txt", header=TRUE)
PinkDucklAdult <- read.delim("Landmarks/Malacorhynchus
membranaceus adult/AllLandmarks.txt", header=TRUE)
Gosling <- read.delim("Landmarks/Anser domesticus hatchling/
AllLandmarks.txt", header=TRUE)
Goose <- read.delim("Landmarks/Anser anser adult/
AllLandmarks.txt", header=TRUE)
Smewling <- read.delim("Landmarks/Mergellus Albellus hatchling/
AllLandmarks.txt", header=TRUE)
Smew <- read.delim("Landmarks/Mergellus Albellus adult/
AllLandmarks.txt", header=TRUE)
RingedTeal <- read.delim("Landmarks/Callonetta leucophrys/
AllLandmarks.txt", header=TRUE)
RingedTealing <- read.delim("Landmarks/Callonetta leucophrys
hatchling/AllLandmarks.txt", header=TRUE)
Partridgling <- read.delim("Landmarks/Rollulus rouloul
hatchling/AllLandmarks.txt", header=TRUE)
```

```
Partridge <- read.delim("Landmarks/Rollulus rouloul adult/
AllLandmarks.txt", header=TRUE)
OxyuraH <- read.delim("Landmarks/Oxyura maccoa hatchling/
AllLandmarks.txt", header=TRUE)
OxyuraA <- read.delim("Landmarks/Oxyura jamaicensis adult/
AllLandmarks.txt", header=TRUE)
Quail <- read.delim("Landmarks/Coturnix coturnix adult/
AllLandmarks.txt", header=TRUE)
Quailing <- read.delim("Landmarks/Coturnix domesticus hatchling/
AllLandmarks.txt", header=TRUE)
Screamling<- read.delim("Landmarks/Chauna chavaria hatchling/
AllLandmarks.txt", header=TRUE)
Screamer<- read.delim("Landmarks/Chauna chavaria/
AllLandmarks.txt", header=TRUE)
Turkling <- read.delim("Landmarks/Meleagris gallopavo hatchling/
AllLandmarks.txt", header=TRUE)
Turkey <- read.delim("Landmarks/Meleagris gallopavo adult/
AllLandmarks.txt", header=TRUE)
CalQuailing<- read.delim("Landmarks/Callipepla californica
hatchling/AllLandmarks.txt", header=TRUE)
CalQuail<- read.delim("Landmarks/Callipepla californica adult/
AllLandmarks.txt", header=TRUE)
Lagopusling <- read.delim("Landmarks/Lagopus lagopus hatchling/
AllLandmarks.txt", header=TRUE)
Lagopus <- read.delim("Landmarks/Lagopus lagopus adult/
AllLandmarks.txt", header=TRUE)
Aixling <- read.delim("Landmarks/Aix sponsa hatchling/
AllLandmarks.txt", header=TRUE)
Aix <- read.delim("Landmarks/Aix galericulata adult/
AllLandmarks.txt", header=TRUE)
EgyptianGosling<- read.delim("Landmarks/Alopochen aegyptiaca
hatchling/AllLandmarks.txt", header=TRUE)
EgyptianGoose<- read.delim("Landmarks/Alopochen aegyptiaca
adult/AllLandmarks.txt", header=TRUE)
Tragoling <- read.delim("Landmarks/Tragopan satyra hatchling/
AllLandmarks.txt", header=TRUE)
Tragopan <- read.delim("Landmarks/Tragopan satyra adult/
AllLandmarks.txt", header=TRUE)
Eiderling <- read.delim("Landmarks/Somateria fischeri hatchling/
AllLandmarks.txt", header=TRUE)
Eider <- read.delim("Landmarks/Somateria fischeri adult/
AllLandmarks.txt", header=TRUE)
Perdixling <- read.delim("Landmarks/Perdix perdix hatchling/
AllLandmarks.txt", header=TRUE)
Perdix <- read.delim("Landmarks/Perdix perdix adult/
AllLandmarks.txt", header=TRUE)
Pealing <- read.delim("Landmarks/Polyplectron bicalcaratum
```

```
hatchling/AllLandmarks.txt", header=TRUE)
Peacockpheasent <- read.delim("Landmarks/Polyplectorn chalcurum
adult/AllLandmarks.txt", header=TRUE)
Goosandling <- read.delim("Landmarks/Mergus serrator hatchling/
AllLandmarks.txt", header=TRUE)
Goosander <- read.delim("Landmarks/Mergus serrator adult/
AllLandmarks.txt", header=TRUE)
chachalacaling <- read.delim("Landmarks/Ortalis ruficauda
hatchling/AllLandmarks.txt", header=TRUE)
chachalaca <- read.delim("Landmarks/Ortalis spps. adult/
AllLandmarks.txt", header=TRUE)
Shoveling <- read.delim("Landmarks/Spatula clypeata hatchling/
AllLandmarks.txt", header=TRUE)
Shoveler <- read.delim("Landmarks/Spatula clypeata adult/
AllLandmarks.txt", header=TRUE)
Shellduckling <- read.delim("Landmarks/Tadorna tadorna
hatchling/AllLandmarks.txt", header=TRUE)
Shellduck <- read.delim("Landmarks/Tadorna tadorna adult/
AllLandmarks.txt", header=TRUE)


###Collection of eroded landmark configurations

##Galloanserae
Magpiegoose<- read.delim("Landmarks/Anseranas semipalmata adult/
AllLandmarks.txt", header=TRUE)
Magpiegosling <- read.delim("Landmarks/Anseranas semipalmata
hatchling/AllLandmarks.txt", header=TRUE)

###Outgroups
Gallinule <- read.delim("Landmarks/Porphyrio marninicus adult/
AllLandmarks.txt", header=TRUE)
Gallinuling <- read.delim("Landmarks/Porphyrio marninicus
hatchling/AllLandmarks.txt", header=TRUE)
Ruffling <- read.delim("Landmarks/Calidris pugnax hatchling/
AllLandmarks.txt", header=TRUE)
Ruff <- read.delim("Landmarks/Calidris pugnax adult/
AllLandmarks.txt", header=TRUE)
Pigeonling <- read.delim("Landmarks/Columba livia hatchling/
AllLandmarks.txt", header=TRUE)
Pigeon<- read.delim("Landmarks/Columba livia adult/
AllLandmarks.txt", header=TRUE)
Tinamouling <- read.delim("Landmarks/Crypturellus tataupa/
AllLandmarks.txt", header=TRUE)
Tinamou <- read.delim("Landmarks/Crypturellus tataupa adult/
AllLandmarks.txt", header=TRUE)
Ostrichling <- read.delim("Landmarks/Struthio camelus hatchling/
```

```
AllLandmarks.txt", header=TRUE)
Ostrich <- read.delim("Landmarks/Struthio camelus adult/
AllLandmarks.txt", header=TRUE)

###Fossils
Presby<- read.delim("Landmarks/Presbyornis/AllLandmarks.txt",
header=TRUE)
Presby2<- read.delim("Landmarks/Presbyornis 2/AllLandmarks.txt",
header=TRUE)
Asteriornis_Recon<- read.delim("Landmarks/Asteriornis Recon/
AllLandmarks.txt", header=TRUE)


#convert to matrices
Allducklandmarks <- data.matrix(duck[,2:4]/1000, rownames.force
= TRUE)
AllducklingLandmarks <- data.matrix(duckling[,2:4]/1000,
rownames.force = TRUE)
Allchickenlandmarks <- data.matrix(chicken[,2:4], rownames.force
= TRUE)
Allchicklandmarks <-data.matrix(chick[,2:4], rownames.force =
TRUE)
Allwducklandmarks <- data.matrix(WDuck[,2:4], rownames.force =
TRUE)
Allwducklinglandmarks <- data.matrix(WDuckling[,2:4],
rownames.force = TRUE)
AllAfropavoHatchlingLandmarks <-
data.matrix(AfropavoHatchling[,2:4], rownames.force = TRUE)
AllAfropavoLandmarks <- data.matrix(Afropavo[,2:4]/1000,
rownames.force = TRUE)
AllPavolingLandmarks <- data.matrix(Pavoling[,2:4],
rownames.force = TRUE)
AllpavoAdultLandmarks <- data.matrix(PavoAdult[,2:4],
rownames.force = TRUE)
AllAythyaHatchlingLandmarks <-
data.matrix(AythyaHatchling[,2:4], rownames.force = TRUE)
AllAythyaAdultLandmarks <- data.matrix(AythyaAdult[,2:4],
rownames.force = TRUE)
AllMegapodlingLandmarks <- data.matrix(megapodling[,2:4],
rownames.force = TRUE)
AllmegapodadultLandmarks <- data.matrix(megapodadult[,2:4],
rownames.force = TRUE)
AllswanlingLandmarks <- data.matrix(swanling[,2:4],
rownames.force = TRUE)
AllswanadultLandmarks <- data.matrix(swanadult[,2:4]/1000,
rownames.force = TRUE)
AllnumidlingLandmarks <- data.matrix(Numidling[,2:4],
```

```
rownames.force = TRUE)
AllnumidiaAdultLandmarks <- data.matrix(NumidiaAdult[,2:4]/1000,
rownames.force = TRUE)
AllPinkDucklingLandmarks <- data.matrix(PinkDuckling[,2:4]/1000,
rownames.force = TRUE)
AllPinkDuckling2Landmarks <- data.matrix(PinkDuckling2[,2:4],
rownames.force = TRUE)
AllPinkDuckLandmarks <- data.matrix(PinkDucklAdult[,2:4],
rownames.force = TRUE)
AllGoslinglandmarks <- data.matrix(Gosling[,2:4], rownames.force
= TRUE)
AllGooselandmarks <- data.matrix(Goose[,2:4]/1000,
rownames.force = TRUE)
AllSmewlinglandmarks <- data.matrix(Smewling[,2:4],
rownames.force = TRUE)
AllSmewlandmarks <- data.matrix(Smew[,2:4], rownames.force =
TRUE)
AllRingedTeallandmarks <- data.matrix(RingedTeal[,2:4],
rownames.force = TRUE)
AllRingedTealinglandmarks <- data.matrix(RingedTealing[,2:4],
rownames.force = TRUE)
AllPartridglinglandmarks <- data.matrix(Partridgling[,2:4]/1000,
rownames.force = TRUE)
AllPartridgelandmarks <- data.matrix(Partridge[,2:4]/1000,
rownames.force = TRUE)
AllOxyuraHlandmarks <- data.matrix(OxyuraH[,2:4], rownames.force
= TRUE)
AllOxyuraAlandmarks <- data.matrix(OxyuraA[,2:4], rownames.force
= TRUE)
Allquaillandmarks <- data.matrix(Quail[,2:4], rownames.force =
TRUE)
Allquailinglandmarks <- data.matrix(Quailing[,2:4],
rownames.force = TRUE)
AllScreamlinglandmarks <- data.matrix(Screamling[,2:4],
rownames.force = TRUE)
AllScreamerlandmarks <- data.matrix(Screamer[,2:4],
rownames.force = TRUE)
AllTurklinglandmarks <- data.matrix(Turkling[,2:4]/1000,
rownames.force = TRUE)
AllTurkeylandmarks <- data.matrix(Turkey[,2:4], rownames.force =
TRUE)
AllCalQuailinglandmarks <- data.matrix(CalQuailing[,2:4],
rownames.force = TRUE)
AllCalQuaillandmarks <- data.matrix(CalQuail[,2:4],
rownames.force = TRUE)
AllLagopuslinglandmarks <- data.matrix(Lagopusling[,2:4],
rownames.force = TRUE)
```

```
AllLagopuslandmarks <- data.matrix(Lagopus[,2:4], rownames.force
= TRUE)
AllAixlinglandmarks <- data.matrix(Aixling[,2:4], rownames.force
= TRUE)
AllAixlandmarks <- data.matrix(Aix[,2:4], rownames.force = TRUE)
AllEgyptianGoslinglandmarks <-
data.matrix(EgyptianGosling[,2:4]/1000, rownames.force = TRUE)
AllEgyptianGooselandmarks <- data.matrix(EgyptianGoose[,2:4]/
1000, rownames.force = TRUE)
AllTragolinglandmarks <- data.matrix(Tragoling[,2:4],
rownames.force = TRUE)
AllTragopanlandmarks <- data.matrix(Tragopan[,2:4],
rownames.force = TRUE)
AllEiderlinglandmarks <- data.matrix(Eiderling[,2:4],
rownames.force = TRUE)
AllEiderlandmarks <- data.matrix(Eider[,2:4], rownames.force =
TRUE)
AllPerdixlinglandmarks <- data.matrix(Perdixling[,2:4],
rownames.force = TRUE)
AllPerdixlandmarks <- data.matrix(Perdix[,2:4], rownames.force =
TRUE)
AllPealinglandmarks <- data.matrix(Pealing[,2:4], rownames.force
= TRUE)
AllPeacockpheasentlandmarks <-
data.matrix(Peacockpheasent[,2:4], rownames.force = TRUE)
AllGoosandlingLandmarks <- data.matrix(Goosandling[,2:4]/1000,
rownames.force = TRUE)
AllGoosanderLandmarks <- data.matrix(Goosander[,2:4]/1000,
rownames.force = TRUE)
AllchachalacalingLandmarks <- data.matrix(chachalacaling[,2:4]/
1000, rownames.force = TRUE)
AllchachalacaLandmarks <- data.matrix(chachalaca[,2:4]/1000,
rownames.force = TRUE)
AllShovelingLandmarks <- data.matrix(Shoveling[,2:4]/1000,
rownames.force = TRUE)
AllShovelerLandmarks <- data.matrix(Shoveler[,2:4]/1000,
rownames.force = TRUE)
AllShellducklingLandmarks <- data.matrix(Shellduckling[,2:4],
rownames.force = TRUE)
AllShellduckLandmarks <- data.matrix(Shellduck[,2:4]/1000,
rownames.force = TRUE)
AllPigeonlingLandmarks <- data.matrix(Pigeonling[,2:4]/1000,
rownames.force = TRUE)
AllPigeonLandmarks <- data.matrix(Pigeon[,2:4]/1000,
rownames.force = TRUE)
AlltinamoulingLandmarks <- data.matrix(Tinamouling[,2:4]/1000,
rownames.force = TRUE)
```

```
AlltinamouLandmarks <- data.matrix(Tinamou[,2:4]/1000,
rownames.force = TRUE)
AllOstrichlingLandmarks <- data.matrix(Ostrichling[,2:4]/1000,
rownames.force = TRUE)
AllOstrichLandmarks <- data.matrix(Ostrich[,2:4]/1000,
rownames.force = TRUE)
AllPresbylandmarks <- data.matrix(Presby[,2:4], rownames.force =
TRUE)
AllPresby2landmarks <- data.matrix(Presby2[,2:4]/1000,
rownames.force = TRUE)
AllAsteriornis_Reconlandmarks <-
data.matrix(Asteriornis_Recon[,2:4]/1000, rownames.force = TRUE)
AllMagpiegooselandmarks <- data.matrix(Magpiegoose[,2:4]/1000,
rownames.force = TRUE)
AllMagpiegoslinglandmarks <- data.matrix(Magpiegosling[,2:4],
rownames.force = TRUE)
AllGallinulelandmarks <- data.matrix(Gallinule[,2:4]/1000,
rownames.force = TRUE)
AllGallinulinglandmarks <- data.matrix(Gallinuling[,2:4]/1000,
rownames.force = TRUE)
AllRufflinglandmarks <- data.matrix(Ruffling[,2:4]/1000,
rownames.force = TRUE)
AllRufflandmarks <- data.matrix(Ruff[,2:4]/1000, rownames.force
= TRUE)


#convert to arrays

##Array_1 is for collection 1 without the Malacorhynchus
membranaceus chick
#Malacorhynchus membranaceus chick is excluded for heatmap
diagrams and other analyses

array_1<-
array(c(Allducklandmarks,AllducklingLandmarks,Allchickenlandmark
s,Allchicklandmarks,Allwducklandmarks,

Allwducklinglandmarks,AllAfropavoHatchlingLandmarks,AllAfropavoL
andmarks,AllPavolingLandmarks,

AllpavoAdultLandmarks,AllAythyaHatchlingLandmarks,AllAythyaAdult
Landmarks,AllMegapodlingLandmarks,

AllmegapodadultLandmarks,AllswanlingLandmarks,AllswanadultLandma
rks, AllnumidlingLandmarks,

AllnumidiaAdultLandmarks,AllPinkDucklingLandmarks,
```

```
AllPinkDuckLandmarks, AllGoslinglandmarks,

AllGooselandmarks,AllSmewlinglandmarks,AllSmewlandmarks,AllRinge
dTeallandmarks,

AllRingedTealinglandmarks,AllPartridglinglandmarks,AllPartridgel
andmarks,AllOxyuraHlandmarks,
                  AllOxyuraAlandmarks,
Allquaillandmarks,Allquailinglandmarks,AllScreamlinglandmarks,

AllScreamerlandmarks,AllTurklinglandmarks,AllTurkeylandmarks,All
CalQuailinglandmarks,

AllCalQuaillandmarks,AllLagopuslinglandmarks,AllLagopuslandmarks
,AllAixlinglandmarks, AllAixlandmarks,

AllEgyptianGoslinglandmarks,AllEgyptianGooselandmarks,AllTragoli
nglandmarks,AllTragopanlandmarks,

AllEiderlinglandmarks,AllEiderlandmarks,AllPerdixlinglandmarks,A
llPerdixlandmarks,

AllPealinglandmarks,AllPeacockpheasentlandmarks,AllGoosandlingLa
ndmarks,AllGoosanderLandmarks,

AllchachalacalingLandmarks,AllchachalacaLandmarks,AllShovelingLa
ndmarks,AllShovelerLandmarks,

AllShellducklingLandmarks,AllShellduckLandmarks), c(321,3,60))
#array_2 includes the Malacorhynchus membranaceus chick
array_2<-
array(c(Allducklandmarks,AllducklingLandmarks,Allchickenlandmark
s,Allchicklandmarks,Allwducklandmarks,

Allwducklinglandmarks,AllAfropavoHatchlingLandmarks,AllAfropavoL
andmarks,AllPavolingLandmarks,

AllpavoAdultLandmarks,AllAythyaHatchlingLandmarks,AllAythyaAdult
Landmarks,AllMegapodlingLandmarks,

AllmegapodadultLandmarks,AllswanlingLandmarks,AllswanadultLandma
rks, AllnumidlingLandmarks,

AllnumidiaAdultLandmarks,AllPinkDucklingLandmarks,
AllPinkDuckling2Landmarks, AllPinkDuckLandmarks,

AllGoslinglandmarks,AllGooselandmarks,AllSmewlinglandmarks,AllSm
```

```
ewlandmarks,AllRingedTeallandmarks,

AllRingedTealinglandmarks,AllPartridglinglandmarks,AllPartridgel
andmarks,AllOxyuraHlandmarks,
                  AllOxyuraAlandmarks,
Allquaillandmarks,Allquailinglandmarks,AllScreamlinglandmarks,

AllScreamerlandmarks,AllTurklinglandmarks,AllTurkeylandmarks,All
CalQuailinglandmarks,

AllCalQuaillandmarks,AllLagopuslinglandmarks,AllLagopuslandmarks
,AllAixlinglandmarks, AllAixlandmarks,

AllEgyptianGoslinglandmarks,AllEgyptianGooselandmarks,AllTragoli
nglandmarks,AllTragopanlandmarks,

AllEiderlinglandmarks,AllEiderlandmarks,AllPerdixlinglandmarks,A
llPerdixlandmarks,

AllPealinglandmarks,AllPeacockpheasentlandmarks,AllGoosandlingLa
ndmarks,AllGoosanderLandmarks,

AllchachalacalingLandmarks,AllchachalacaLandmarks,AllShovelingLa
ndmarks,AllShovelerLandmarks,

AllShellducklingLandmarks,AllShellduckLandmarks), c(321,3,61))
#array_3 is for collection 2

Erodedarray <- array_1[c(1:9,14:16,25:26,50:111,140:163),,1:60]
# subsample the first array
##add the other clades
Other_clades_array.1 <-
array(c(AllPigeonlingLandmarks,AllPigeonLandmarks,Alltinamouling
Landmarks,AlltinamouLandmarks,

AllOstrichlingLandmarks,AllOstrichLandmarks), c(321,3,6))
Other_clades_array.2<-
Other_clades_array.1[c(1:9,14:16,25:26,50:111,140:163),,1:6]

Other_clades_array.3 <-
array(c(AllPresbylandmarks,AllPresby2landmarks,
AllAsteriornis_Reconlandmarks,AllMagpiegooselandmarks,

AllMagpiegoslinglandmarks,AllGallinulelandmarks,AllGallinulingla
ndmarks,AllRufflinglandmarks,AllRufflandmarks), c(100,3,9))

array_3 <-
```

```
abind(Erodedarray,Other_clades_array.2,Other_clades_array.3)#com
bine the two arrays

##array dimensions

##array_1
taxons_1 <- as.factor(c("Anas", "Anas", "Gallus", "Gallus",
"Dendrocygna", "Dendrocygna", "Afropavo", "Afropavo",
                         "Pavo", "Pavo", "Aythya", "Aythya",
"Megapodius","Megapodius", "Cygnus", "Cygnus", "Numida",
                         "Numida","Malacorhynchus",
"Malacorhynchus", "Anser", "Anser", "Mergellus",
                         "Mergellus","Callonetta","Callonetta",
"Rollulus", "Rollulus", "Oxyura","Oxyura", "Coturnix",
                         "Coturnix", "Chauna", "Chauna",
"Meleagris", "Meleagris","Callipepla","Callipepla", "Lagopus",
                         "Lagopus",
"Aix","Aix","Alopochen","Alopochen","Tragopan","Tragopan",
"Somateria", "Somateria",
                         "Perdix","Perdix","Polyplectron",
"Polyplectron","Mergus","Mergus","Ortalis","Ortalis", "Spatula",
                         "Spatula","Tadorna","Tadorna"))
families_1 <- as.factor(c("Anseriformes", "Anseriformes",
"Galliformes", "Galliformes", "Anseriformes",
                           "Anseriformes", "Galliformes",
"Galliformes", "Galliformes", "Galliformes",
                           "Anseriformes", "Anseriformes",
"Galliformes", "Galliformes", "Anseriformes",
                           "Anseriformes", "Galliformes",
"Galliformes","Anseriformes","Anseriformes",
                           "Anseriformes", "Anseriformes",
"Anseriformes", "Anseriformes", "Anseriformes",
                           "Anseriformes", "Galliformes",
"Galliformes", "Anseriformes","Anseriformes",
                           "Galliformes",
"Galliformes","Anseriformes","Anseriformes", "Galliformes",
                           "Galliformes", "Galliformes",
"Galliformes", "Galliformes", "Galliformes",

"Anseriformes","Anseriformes","Anseriformes","Anseriformes",
"Galliformes",

"Galliformes","Anseriformes","Anseriformes", "Galliformes",
"Galliformes",
                           "Galliformes",
"Galliformes","Anseriformes","Anseriformes", "Galliformes",
"Galliformes",
```

```
"Anseriformes","Anseriformes","Anseriformes","Anseriformes"))

age_1 <-
c("adult","hatchling","adult","hatchling","adult","hatchling","h
atchling","adult","hatchling","adult","hatchling","adult","hatch
ling","adult","hatchling","adult","hatchling","adult","hatchling
","adult","hatchling","adult","hatchling","adult","adult","hatch
ling","hatchling","adult","hatchling","adult","adult","hatchling
","hatchling","adult","hatchling","adult","hatchling","adult","h
atchling","adult","hatchling","adult","hatchling","adult","hatch
ling","adult","hatchling","adult","hatchling","adult","hatchling
","adult","hatchling","adult","hatchling","adult","hatchling","a
dult","hatchling","adult")

age_1 <- as.factor(age_1)
taxon.names <- interaction(taxons_1,age_1)
dimnames(array_1)[[3]]<- taxon.names
dimnames(array_1)[[2]]<- c("x","y","z")

##array_2
taxons_2 <- as.factor(c("Anas", "Anas", "Gallus", "Gallus",
"Dendrocygna", "Dendrocygna", "Afropavo", "Afropavo",
                        "Pavo", "Pavo", "Aythya", "Aythya",
"Megapodius","Megapodius", "Cygnus", "Cygnus", "Numida",
                        "Numida","Malacorhynchus",
"Malacorhynchus","Malacorhynchus", "Anser", "Anser",
"Mergellus",
                        "Mergellus","Callonetta","Callonetta",
"Rollulus", "Rollulus", "Oxyura","Oxyura", "Coturnix",
                        "Coturnix", "Chauna", "Chauna",
"Meleagris", "Meleagris","Callipepla","Callipepla", "Lagopus",
                        "Lagopus",
"Aix","Aix","Alopochen","Alopochen","Tragopan","Tragopan",
"Somateria", "Somateria",
                        "Perdix","Perdix","Polyplectron",
"Polyplectron","Mergus","Mergus","Ortalis","Ortalis", "Spatula",
                        "Spatula","Tadorna","Tadorna"))
families_2 <- as.factor(c("Anseriformes", "Anseriformes",
"Galliformes", "Galliformes", "Anseriformes",
                          "Anseriformes", "Galliformes",
"Galliformes", "Galliformes", "Galliformes",
                          "Anseriformes", "Anseriformes",
"Galliformes", "Galliformes", "Anseriformes","Anseriformes",
                          "Anseriformes", "Galliformes",
"Galliformes","Anseriformes","Anseriformes",
                          "Anseriformes", "Anseriformes",
```

```
"Anseriformes", "Anseriformes", "Anseriformes",
                          "Anseriformes", "Galliformes",
"Galliformes", "Anseriformes","Anseriformes",
                          "Galliformes",
"Galliformes","Anseriformes","Anseriformes", "Galliformes",
                          "Galliformes", "Galliformes",
"Galliformes", "Galliformes", "Galliformes",

"Anseriformes","Anseriformes","Anseriformes","Anseriformes",
"Galliformes",

"Galliformes","Anseriformes","Anseriformes", "Galliformes",
"Galliformes",
                          "Galliformes",
"Galliformes","Anseriformes","Anseriformes", "Galliformes",
"Galliformes",

"Anseriformes","Anseriformes","Anseriformes","Anseriformes"))

age_2 <-
c("adult","hatchling","adult","hatchling","adult","hatchling","h
atchling","adult","hatchling","adult","hatchling",

"adult","hatchling","adult","hatchling","adult","hatchling","adu
lt","hatchling","chick","adult","hatchling","adult",

"hatchling","adult","adult","hatchling","hatchling","adult","hat
chling","adult","adult","hatchling","hatchling",

"adult","hatchling","adult","hatchling","adult","hatchling","adu
lt","hatchling","adult","hatchling","adult",

"hatchling","adult","hatchling","adult","hatchling","adult","hat
chling","adult","hatchling","adult","hatchling",
           "adult","hatchling","adult","hatchling","adult")

age_2 <- as.factor(age_2)
taxon.names <- interaction(taxons_2,age_2)
dimnames(array_2)[[3]]<- taxon.names
dimnames(array_2)[[2]]<- c("x","y","z")

###array_3
taxons_3 <- as.factor(c("Anas", "Anas", "Gallus", "Gallus",
"Dendrocygna", "Dendrocygna", "Afropavo", "Afropavo",
                        "Pavo", "Pavo", "Aythya", "Aythya",
"Megapodius","Megapodius", "Cygnus", "Cygnus", "Numida",
"Numida",
```

```
                        "Malacorhynchus", "Malacorhynchus",
"Anser", "Anser", "Mergellus", "Mergellus","Callonetta",
                        "Callonetta", "Rollulus", "Rollulus",
"Oxyura","Oxyura", "Coturnix","Coturnix", "Chauna",
                        "Chauna", "Meleagris",
"Meleagris","Callipepla","Callipepla", "Lagopus", "Lagopus",
"Aix","Aix",

"Alopochen","Alopochen","Tragopan","Tragopan", "Somateria",
"Somateria","Perdix","Perdix",
                        "Polyplectron",
"Polyplectron","Mergus","Mergus","Ortalis","Ortalis", "Spatula",
"Spatula",
                        "Tadorna","Tadorna","Columba",
"Columba","Crypturellus","Crypturellus", "Struthio",
"Struthio","Presbyornis",
                        "Presbyornis
2","Asteriornis","Anseranas",
"Anseranas","Porphyrio","Porphyrio", "Calidris", "Calidris"))
families_3 <- as.factor(c("Anseriformes", "Anseriformes",
"Galliformes", "Galliformes", "Anseriformes",
                          "Anseriformes", "Galliformes",
"Galliformes", "Galliformes", "Galliformes",
                          "Anseriformes", "Anseriformes",
"Galliformes", "Galliformes", "Anseriformes",
                          "Anseriformes", "Galliformes",
"Galliformes","Anseriformes", "Anseriformes",
                          "Anseriformes", "Anseriformes",
"Anseriformes", "Anseriformes","Anseriformes",
                          "Anseriformes", "Galliformes",
"Galliformes", "Anseriformes","Anseriformes",
                          "Galliformes",
"Galliformes","Anseriformes","Anseriformes", "Galliformes",
                          "Galliformes", "Galliformes",
"Galliformes", "Galliformes", "Galliformes",

"Anseriformes","Anseriformes","Anseriformes","Anseriformes",
"Galliformes",

"Galliformes","Anseriformes","Anseriformes", "Galliformes",
"Galliformes",
                          "Galliformes",
"Galliformes","Anseriformes","Anseriformes", "Galliformes",

"Galliformes","Anseriformes","Anseriformes","Anseriformes","Anse
riformes",
                          "Columbiformes",
```

```
"Columbiformes","Tinamiformes","Tinamiformes",
"Struthioniformes",

"Struthioniformes","PanAnseriformes","PanAnseriformes","PanGallo
anserae", "Anseriformes",
                          "Anseriformes", "Gruiformes",
"Gruiformes","Charadriiformes","Charadriiformes"))
age_3 <-
c("adult","hatchling","adult","hatchling","adult","hatchling","h
atchling","adult","hatchling","adult",

"hatchling","adult","hatchling","adult","hatchling","adult","hat
chling","adult","hatchling","adult",

"hatchling","adult","hatchling","adult","adult","hatchling","hat
chling","adult","hatchling","adult",

"adult","hatchling","hatchling","adult","hatchling","adult","hat
chling","adult","hatchling","adult",

"hatchling","adult","hatchling","adult","hatchling","adult","hat
chling","adult","hatchling","adult",

"hatchling","adult","hatchling","adult","hatchling","adult","hat
chling","adult","hatchling","adult",

"hatchling","adult","hatchling","adult","hatchling","adult","adu
lt","adult","adult","adult",
           "hatchling","adult","hatchling","hatchling","adult")

age_3 <- as.factor(age_3)
taxon.names <- interaction(taxons_3,age_3)
dimnames(array_3)[[3]]<- taxon.names
dimnames(array_3)[[2]]<- c("x","y","z")

#### Import semilandmarks

Semilandmarklocations <- read.csv("Landmarks/
Semilandmarklocations.csv") ## for arrays 1 and 2

FossilSemilandmarklocations <- read.csv("Landmarks/
FossilSemiLandmarkLocations.csv") ## for array 3


##### Generalized Procrustes Analysis

gpa_1 <- gpagen(array_1, curves = Semilandmarklocations,
```

```
max.iter = NULL,tol = 1e-04, ProcD = FALSE, approxBE = TRUE, sen
= 0.5, Proj = TRUE, verbose = T,print.progress = TRUE, Parallel
= FALSE)
gpa_2 <- gpagen(array_2, curves = Semilandmarklocations,
max.iter = NULL,tol = 1e-04, ProcD = FALSE, approxBE = TRUE, sen
= 0.5, Proj = TRUE, verbose = T,print.progress = TRUE, Parallel
= FALSE)
gpa_3 <- gpagen(array_3, curves = FossilSemilandmarklocations,
max.iter = NULL,tol = 1e-04, ProcD = FALSE, approxBE = TRUE, sen
= 0.5, Proj = TRUE, verbose = T,print.progress = TRUE, Parallel
= FALSE)


###PCA Analysis using array_1

pca<- gm.prcomp(gpa_1$coords)

######PCA plot

specimen.colors <- c("#4E99B7", "#4E99B7", "#4EB773", "#4EB773",
"#4E99B7",
                     "#4E99B7", "#4EB773", "#4EB773", "#4EB773",
"#4EB773",
                     "#4E99B7", "#4E99B7", "#4EB773", "#4EB773",
"#4E99B7",
                     "#4E99B7", "#4EB773", "#4EB773","#4E99B7",
"#4E99B7","#4E99B7",
                     "#4E99B7", "#4E99B7", "#4E99B7", "#4E99B7",
"#4E99B7",
                     "#4E99B7", "#4EB773", "#4EB773",
"#4E99B7","#4E99B7",
                     "#4EB773", "#4EB773","#4E99B7","#4E99B7",
"#4EB773",
                     "#4EB773", "#4EB773", "#4EB773", "#4EB773",
"#4EB773",
                     "#4E99B7","#4E99B7","#4E99B7","#4E99B7",
"#4EB773",
                     "#4EB773","#4E99B7","#4E99B7", "#4EB773",
"#4EB773",
                     "#4EB773", "#4EB773","#4E99B7","#4E99B7",
"#4EB773",

"#4EB773","#4E99B7","#4E99B7","#4E99B7","#4E99B7")

shapes <- c(23, 21, 23, 21, 23, 21, 21, 23, 21,23,
            21, 23, 21,23, 21, 23, 21, 23, 21,21,
            23, 21, 23, 21, 23, 23, 21, 21, 23,
```

```
            21,23,23, 21, 21,23, 21,23, 21,23,
            21,23, 21,23, 21,23, 21,23, 21,23,
            21,23, 21,23, 21,23, 21,23, 21,23,
            21,23)

ggplot(data=pca$x[,1:2], aes(x=pca$x[,1], y= pca$x[,2], color =
specimen.colors, legend = F, group = age_2, label = taxons_2))+
  theme(text = element_text(family = "Helvetica"))+
  geom_point(shape = shapes,size = 4, stroke= 0.2,color =
"black",fill =specimen.colors, alpha =
10)+scale_color_manual(values=c("#4E99B7", "#5EB16B"))+
scale_shape_manual(values=c(21, 22))+scale_y_reverse()+
  theme(axis.title.x = element_blank(),axis.text.x =
element_blank(),axis.text.y = element_blank(),axis.title.y =
element_blank(),axis.line = element_line(colour =
"black"),panel.grid.major = element_blank(),panel.background =
element_blank(),panel.grid.minor = element_blank())
geom_segment(aes(x = pca$x[2], y = pca$x[2,2], xend = pca$x[1],
yend = pca$x[1,2]), linewidth = 1,color ='#4E99B7',alpha =
0.009)+
  geom_segment(aes(x = pca$x[4], y = pca$x[4,2], xend =
pca$x[3], yend = pca$x[3,2]), linewidth = 1,color
='#5EB16B',alpha = 0.009)+
  geom_segment(aes(x = pca$x[6], y = pca$x[6,2], xend =
pca$x[5], yend = pca$x[5,2]), linewidth = 1,color
='#4E99B7',alpha = 0.009)+
  geom_segment(aes(x = pca$x[7], y = pca$x[7,2], xend =
pca$x[8], yend = pca$x[8,2]), linewidth = 1,color
='#5EB16B',alpha = 0.009)+
  geom_segment(aes(x = pca$x[9], y = pca$x[9,2], xend =
pca$x[10], yend = pca$x[10,2]), linewidth = 1,color
='#5EB16B',alpha = 0.009)+
  geom_segment(aes(x = pca$x[11], y = pca$x[11,2], xend =
pca$x[12], yend = pca$x[12,2]), linewidth = 1,color
='#4E99B7',alpha = 0.009)+
  geom_segment(aes(x = pca$x[13], y = pca$x[13,2], xend =
pca$x[14], yend = pca$x[14,2]), linewidth = 1,color
='#5EB16B',alpha = 0.009)+
  geom_segment(aes(x = pca$x[15], y = pca$x[15,2], xend =
pca$x[16], yend = pca$x[16,2]), linewidth = 1,color
='#4E99B7',alpha = 0.009)+
  geom_segment(aes(x = pca$x[17], y = pca$x[17,2], xend =
pca$x[18], yend = pca$x[18,2]), linewidth = 1,color
='#5EB16B',alpha = 0.009)+
  geom_segment(aes(x = pca$x[19], y = pca$x[19,2], xend =
pca$x[20], yend = pca$x[20,2]), linewidth = 1,color
='#4E99B7',alpha = 0.009)+
```

```
  geom_segment(aes(x = pca$x[20], y = pca$x[20,2], xend =
pca$x[21], yend = pca$x[21,2]), linewidth = 1,color
='#4E99B7',alpha = 0.009)+
  geom_segment(aes(x = pca$x[22], y = pca$x[22,2], xend =
pca$x[23], yend = pca$x[23,2]), linewidth = 1,color
='#4E99B7',alpha = 0.009)+
  geom_segment(aes(x = pca$x[24], y = pca$x[24,2], xend =
pca$x[25], yend = pca$x[25,2]), linewidth = 1,color
='#4E99B7',alpha = 0.009)+
  geom_segment(aes(x = pca$x[27], y = pca$x[27,2], xend =
pca$x[26], yend = pca$x[26,2]), linewidth = 1,color
='#4E99B7',alpha = 0.009)+
  geom_segment(aes(x = pca$x[28], y = pca$x[28,2], xend =
pca$x[29], yend = pca$x[29,2]), linewidth = 1,color
='#5EB16B',alpha = 0.009)+
  geom_segment(aes(x = pca$x[30], y = pca$x[30,2], xend =
pca$x[31], yend = pca$x[31,2]), linewidth = 1,color
='#4E99B7',alpha = 0.009)+
  geom_segment(aes(x = pca$x[33], y = pca$x[33,2], xend =
pca$x[32], yend = pca$x[32,2]), linewidth = 1,color
='#5EB16B',alpha = 0.009)+
  geom_segment(aes(x = pca$x[34], y = pca$x[34,2], xend =
pca$x[35], yend = pca$x[35,2]), linewidth = 1,color
='#4E99B7',alpha = 0.009)+
  geom_segment(aes(x = pca$x[36], y = pca$x[36,2], xend =
pca$x[37], yend = pca$x[37,2]), linewidth = 1,color
='#5EB16B',alpha = 0.009)+
  geom_segment(aes(x = pca$x[38], y = pca$x[38,2], xend =
pca$x[39], yend = pca$x[39,2]), linewidth = 1,color
='#5EB16B',alpha = 0.009)+
  geom_segment(aes(x = pca$x[40], y = pca$x[40,2], xend =
pca$x[41], yend = pca$x[41,2]), linewidth = 1,color
='#5EB16B',alpha = 0.009)+
  geom_segment(aes(x = pca$x[42], y = pca$x[42,2], xend =
pca$x[43], yend = pca$x[43,2]), linewidth = 1,color
='#4E99B7',alpha = 0.009)+
  geom_segment(aes(x = pca$x[44], y = pca$x[44,2], xend =
pca$x[45], yend = pca$x[45,2]), linewidth = 1,color
='#4E99B7',alpha = 0.009)+
  geom_segment(aes(x = pca$x[46], y = pca$x[46,2], xend =
pca$x[47], yend = pca$x[47,2]), linewidth = 1,color
='#5EB16B',alpha = 0.009)+
  geom_segment(aes(x = pca$x[48], y = pca$x[48,2], xend =
pca$x[49], yend = pca$x[49,2]), linewidth = 1,color
='#4E99B7',alpha = 0.009)+
  geom_segment(aes(x = pca$x[50], y = pca$x[50,2], xend =
pca$x[51], yend = pca$x[51,2]), linewidth = 1,color
```

```
='#5EB16B',alpha = 0.009)+
  geom_segment(aes(x = pca$x[52], y = pca$x[52,2], xend =
pca$x[53], yend = pca$x[53,2]), linewidth = 1,color
='#5EB16B',alpha = 0.009)+
  geom_segment(aes(x = pca$x[54], y = pca$x[54,2], xend =
pca$x[55], yend = pca$x[55,2]), linewidth = 1,color
='#4E99B7',alpha = 0.009)+
  geom_segment(aes(x = pca$x[56], y = pca$x[56,2], xend =
pca$x[57], yend = pca$x[57,2]), linewidth = 1,color
='#5EB16B',alpha = 0.009)+
  geom_segment(aes(x = pca$x[58], y = pca$x[58,2], xend =
pca$x[59], yend = pca$x[59,2]), linewidth = 1,color
='#4E99B7',alpha = 0.009)+
  geom_segment(aes(x = pca$x[60], y = pca$x[60,2], xend =
pca$x[61], yend = pca$x[61,2]), linewidth = 1,color
='#4E99B7',alpha = 0.009)


######PCA axis shapes

per_lm_variance <- function(shape.data){

  variances<-rowSums(apply(shape.data ,c(1,2),var))



  cols1<-
colorRampPalette(c("#D8C5C5","#D8BABA","#D8ADAD","#D8A3A3","#D89
393","#D87C7C","#D86060","red"))

  cols<-cols1(100)

  #calculate log rates:
  x=(log10(variances))
  xlims<-NULL
  tol <- 1e-06
  xlims <- range(x) + c(-tol, tol)
  nbin=100
  breaks <- 0:nbin/nbin * (xlims[2] - xlims[1]) + xlims[1]
  whichColor <- function(p, cols, breaks) {
    i <- 1
    while (p >= breaks[i] && p > breaks[i + 1]) i <- i +
        1
    cols[i]
  }
  variancecolors <- sapply(x, whichColor, cols = cols, breaks =
breaks)
```

```
  variance_table <- tibble("Per_Lm_Variance" = variances,
"Log_Variance" = x, "Variance_Colors" = variancecolors)
  return(variance_table)
} ##function for visualising per landmark variance

my.shapes <- gpa_2$coords
my.variances <- per_lm_variance(shape.data = my.shapes)

mean.spec.name <- findMeanSpec(gpa_2$coords)
mesh.original <- read.ply("Landmarks/Cygnus melancoryphus
hatchling/Upper Skull.ply", ShowSpecimen = FALSE, addNormals =
TRUE)
coords.original <- array_2[,,15]
M <- mshape(gpa_2$coords)

#PC1
PC <- 1

PC.temp<-pca$x[,1]

preds<-shape.predictor(gpa_2$coords, x =
as.numeric( pca$x[,PC] ), Intercept = F,
                        pred1 = quantile(PC.temp, probs = 0.1),
pred2 = quantile(PC.temp, probs = 0.9))


# now we need to put all shapes in the same space to plot warped
mesh and hot.dots on top of it

open3d()
par3d(windowRect = c(0,0,700,250))
Sys.sleep(1)
mfrow3d(nr = 1, nc = 2, byrow = TRUE, sharedMouse = TRUE)
mesh.min <- tps3d(mesh.original, refmat = coords.original,
tarmat = preds$pred1)
shade3d(mesh.min, col= 8, alpha = 1)
spheres3d(preds$pred1, col = my.variances$Variance_Colors,
radius = 0.003)



next3d()
mesh.max <- tps3d(mesh.original, refmat = coords.original,
tarmat = preds$pred2)
shade3d(mesh.max, col= 8, alpha = 1)
```

```
spheres3d(preds$pred2, col = my.variances$Variance_Colors,
radius = 0.003)


#PC2
PC <- 2

PC.temp<-pca$x[,2]

preds<-shape.predictor(gpa_2$coords, x =
as.numeric( pca$x[,PC] ), Intercept = F,
                       pred1 = quantile(PC.temp, probs = 0.1),
pred2 = quantile(PC.temp, probs = 0.9))


# now we need to put all shapes in the same space to plot warped
mesh and hot.dots on top of it

open3d()
par3d(windowRect = c(0,0,700,250))
Sys.sleep(1)
mfrow3d(nr = 1, nc = 2, byrow = TRUE, sharedMouse = TRUE)
mesh.min <- tps3d(mesh.original, refmat = coords.original,
tarmat = preds$pred1)
shade3d(mesh.min, col= 8, alpha = 1)
spheres3d(preds$pred1, col = my.variances$Variance_Colors,
radius = 0.003)


next3d()
mesh.max <- tps3d(mesh.original, refmat = coords.original,
tarmat = preds$pred2)
shade3d(mesh.max, col= 8, alpha = 1)
spheres3d(preds$pred2, col = my.variances$Variance_Colors,
radius = 0.003)


####################################
###########Allometry analysis#######
####################################
##Regression analysis of shape on size

gdf <- geomorph.data.frame(coords = gpa_2$coords, CS=
log10(gpa_2$Csize), age=age_2, taxons = taxons_2, families =
families_2 )
fit <- procD.lm(coords ~ CS, data = gdf, iter = 9999,
print.progress = T)
```

```
summary(fit, stat.table =T)

##Regression plot
allom.plot <- plot(fit, type = "regression", predictor =
log10(gpa_2$Csize), reg.type ="RegScore")
allo.data <- data.frame(gdf$CS,allom.plot$RegScore)
ggplot(data=allo.data, aes(x=gdf$CS, y= allom.plot$RegScore,
color = specimen.colors, legend = F, group = age_2, label =
taxons_2))+
  theme(text = element_text(family = "Helvetica"))
+geom_point(shape = shapes,size = 5, stroke= 0.1,color =
"black",fill =specimen.colors, alpha =
1)+scale_color_manual(values=c("#4E99B7", "#5EB16B"))+
scale_shape_manual(values=c(21, 22))+
  theme(axis.title.x = element_blank(),axis.text.x =
element_blank(),axis.text.y = element_blank(),axis.title.y =
element_blank(),axis.line = element_line(colour =
"black"),panel.grid.major = element_blank(),panel.background =
element_blank(),panel.grid.minor = element_blank())+
  geom_segment(aes(x = gdf$CS[2], y = allom.plot$RegScore[2],
xend = gdf$CS[1], yend = allom.plot$RegScore[1]), linewidth =
1,color ='#4E99B7',alpha = 0.009)+
  geom_segment(aes(x = gdf$CS[4], y = allom.plot$RegScore[4],
xend = gdf$CS[3], yend = allom.plot$RegScore[3]), linewidth =
1,color ='#5EB16B',alpha = 0.009)+
  geom_segment(aes(x = gdf$CS[6], y = allom.plot$RegScore[6],
xend = gdf$CS[5], yend = allom.plot$RegScore[5]), linewidth =
1,color ='#4E99B7',alpha = 0.009)+
  geom_segment(aes(x = gdf$CS[7], y = allom.plot$RegScore[7],
xend = gdf$CS[8], yend = allom.plot$RegScore[8]), linewidth =
1,color ='#5EB16B',alpha = 0.009)+
  geom_segment(aes(x = gdf$CS[9], y = allom.plot$RegScore[9],
xend = gdf$CS[10], yend = allom.plot$RegScore[10]), linewidth =
1,color ='#5EB16B',alpha = 0.009)+
  geom_segment(aes(x = gdf$CS[11], y = allom.plot$RegScore[11],
xend = gdf$CS[12], yend = allom.plot$RegScore[12]), linewidth =
1,color ='#4E99B7',alpha = 0.009)+
  geom_segment(aes(x = gdf$CS[13], y = allom.plot$RegScore[13],
xend = gdf$CS[14], yend = allom.plot$RegScore[14]), linewidth =
1,color ='#5EB16B',alpha = 0.009)+
  geom_segment(aes(x = gdf$CS[15], y = allom.plot$RegScore[15],
xend = gdf$CS[16], yend = allom.plot$RegScore[16]), linewidth =
1,color ='#4E99B7',alpha = 0.009)+
  geom_segment(aes(x = gdf$CS[17], y = allom.plot$RegScore[17],
xend = gdf$CS[18], yend = allom.plot$RegScore[18]), linewidth =
1,color ='#5EB16B',alpha = 0.009)+
  geom_segment(aes(x = gdf$CS[19], y = allom.plot$RegScore[19],
```

```
xend = gdf$CS[20], yend = allom.plot$RegScore[20]), linewidth =
1,color ='#4E99B7',alpha = 0.009)+
  geom_segment(aes(x = gdf$CS[20], y = allom.plot$RegScore[20],
xend = gdf$CS[21], yend = allom.plot$RegScore[21]), linewidth =
1,color ='#4E99B7',alpha = 0.009)+
  geom_segment(aes(x = gdf$CS[22], y = allom.plot$RegScore[22],
xend = gdf$CS[23], yend = allom.plot$RegScore[23]), linewidth =
1,color ='#4E99B7',alpha = 0.009)+
  geom_segment(aes(x = gdf$CS[24], y = allom.plot$RegScore[24],
xend = gdf$CS[25], yend = allom.plot$RegScore[25]), linewidth =
1,color ='#4E99B7',alpha = 0.009)+
  geom_segment(aes(x = gdf$CS[27], y = allom.plot$RegScore[27],
xend = gdf$CS[26], yend = allom.plot$RegScore[26]), linewidth =
1,color ='#4E99B7',alpha = 0.009)+
  geom_segment(aes(x = gdf$CS[28], y = allom.plot$RegScore[28],
xend = gdf$CS[29], yend = allom.plot$RegScore[29]), linewidth =
1,color ='#5EB16B',alpha = 0.009)+
  geom_segment(aes(x = gdf$CS[30], y = allom.plot$RegScore[30],
xend = gdf$CS[31], yend = allom.plot$RegScore[31]), linewidth =
1,color ='#4E99B7',alpha = 0.009)+
  geom_segment(aes(x = gdf$CS[33], y = allom.plot$RegScore[33],
xend = gdf$CS[32], yend = allom.plot$RegScore[32]), linewidth =
1,color ='#5EB16B',alpha = 0.009)+
  geom_segment(aes(x = gdf$CS[34], y = allom.plot$RegScore[34],
xend = gdf$CS[35], yend = allom.plot$RegScore[35]), linewidth =
1,color ='#4E99B7',alpha = 0.009)+
  geom_segment(aes(x = gdf$CS[36], y = allom.plot$RegScore[36],
xend = gdf$CS[37], yend = allom.plot$RegScore[37]), linewidth =
1,color ='#5EB16B',alpha = 0.009)+
  geom_segment(aes(x = gdf$CS[38], y = allom.plot$RegScore[38],
xend = gdf$CS[39], yend = allom.plot$RegScore[39]), linewidth =
1,color ='#5EB16B',alpha = 0.009)+
  geom_segment(aes(x = gdf$CS[40], y = allom.plot$RegScore[40],
xend = gdf$CS[41], yend = allom.plot$RegScore[41]), linewidth =
1,color ='#5EB16B',alpha = 0.009)+
  geom_segment(aes(x = gdf$CS[42], y = allom.plot$RegScore[42],
xend = gdf$CS[43], yend = allom.plot$RegScore[43]), linewidth =
1,color ='#4E99B7',alpha = 0.009)+
  geom_segment(aes(x = gdf$CS[44], y = allom.plot$RegScore[44],
xend = gdf$CS[45], yend = allom.plot$RegScore[45]), linewidth =
1,color ='#4E99B7',alpha = 0.009)+
  geom_segment(aes(x = gdf$CS[46], y = allom.plot$RegScore[46],
xend = gdf$CS[47], yend = allom.plot$RegScore[47]), linewidth =
1,color ='#5EB16B',alpha = 0.009)+
  geom_segment(aes(x = gdf$CS[48], y = allom.plot$RegScore[48],
xend = gdf$CS[49], yend = allom.plot$RegScore[49]), linewidth =
1,color ='#4E99B7',alpha = 0.009)+
```

```
  geom_segment(aes(x = gdf$CS[50], y = allom.plot$RegScore[50],
xend = gdf$CS[51], yend = allom.plot$RegScore[51]), linewidth =
1,color ='#5EB16B',alpha = 0.009)+
  geom_segment(aes(x = gdf$CS[52], y = allom.plot$RegScore[52],
xend = gdf$CS[53], yend = allom.plot$RegScore[53]), linewidth =
1,color ='#5EB16B',alpha = 0.009)+
  geom_segment(aes(x = gdf$CS[54], y = allom.plot$RegScore[54],
xend = gdf$CS[55], yend = allom.plot$RegScore[55]), linewidth =
1,color ='#4E99B7',alpha = 0.009)+
  geom_segment(aes(x = gdf$CS[56], y = allom.plot$RegScore[56],
xend = gdf$CS[57], yend = allom.plot$RegScore[57]), linewidth =
1,color ='#5EB16B',alpha = 0.009)+
  geom_segment(aes(x = gdf$CS[58], y = allom.plot$RegScore[58],
xend = gdf$CS[59], yend = allom.plot$RegScore[59]), linewidth =
1,color ='#4E99B7',alpha = 0.009)+
  geom_segment(aes(x = gdf$CS[60], y = allom.plot$RegScore[60],
xend = gdf$CS[61], yend = allom.plot$RegScore[61]), linewidth =
1,color ='#4E99B7',alpha = 0.009)#+geom_label()

#Regression score axis shapes

####regression score shapes

RS <- 1


preds<-shape.predictor(gpa$coords, x =
as.numeric(allom.plot$RegScore[,RS] ), Intercept = F,
                       pred1 = quantile(allom.plot$RegScore,
probs = 0.1), pred2 = quantile(allom.plot$RegScore, probs =
0.9))

open3d()
par3d(windowRect = c(0,0,700,250))
Sys.sleep(1)
mfrow3d(nr = 1, nc = 2, byrow = TRUE, sharedMouse = TRUE)
mesh.min <- tps3d(mesh.original, refmat = coords.original,
tarmat = preds$pred1)
shade3d(mesh.min, col= 8, alpha = 1)
spheres3d(preds$pred1, col = my.variances$Variance_Colors,
radius = 0.003)


next3d()
mesh.max <- tps3d(mesh.original, refmat = coords.original,
tarmat = preds$pred2)
shade3d(mesh.max, col= 8, alpha = 1)
```

```
spheres3d(preds$pred2, col = my.variances$Variance_Colors,
radius = 0.003)

rgl.snapshot( "RS_min_max.shapes.png" )


###plot for mean heterochronic components of evolution

Hatchling.Galliformes<- intersect(which(age_1 ==
"hatchling"),which(families_1 == "Galliformes"))
Hatchling.Anseriformes<- intersect(which(age_1 ==
"hatchling"),which(families_1 == "Anseriformes"))
Adult.Galliformes<- intersect(which(age_1 ==
"adult"),which(families_1 == "Galliformes"))
Adult.Anseriformes<- intersect(which(age_1 ==
"adult"),which(families_1 == "Anseriformes"))

mean.galli.hatch.size<-
mean(log10(gpa_1$Csize[Hatchling.Galliformes]))
mean.galli.hatch.shape <-
mshape(gpa_1$coords[,,Hatchling.Galliformes])
mean.anseri.hatch.size<-
mean(log10(gpa_1$Csize[Hatchling.Anseriformes]))
mean.anseri.hatch.shape <-
mshape(gpa_1$coords[,,Hatchling.Anseriformes])
mean.galli.adult.size<-
mean(log10(gpa_1$Csize[Adult.Galliformes]))
mean.galli.adult.shape<-
mshape(gpa_1$coords[,,Adult.Galliformes])
mean.anseri.adult.size<-
mean(log10(gpa_1$Csize[Adult.Anseriformes]))
mean.anseri.adult.shape<-
mshape(gpa_1$coords[,,Adult.Anseriformes])

sizes <-
c(mean.galli.hatch.size,mean.anseri.hatch.size,mean.galli.adult.
size,mean.anseri.adult.size)
shapes <-
array(c(mean.galli.hatch.shape,mean.anseri.hatch.shape,mean.gall
i.adult.shape,mean.anseri.adult.shape),c(321,3,4))
families.2 <-
c("Galliformes","Anseriformes","Galliformes","Anseriformes")

gdf.2 <- geomorph.data.frame(coords = shapes, CS= sizes,
families = families.2 )
fit.2 <- procD.lm(coords ~ CS, data = gdf.2, iter = 9999,
print.progress = T)
```

```
allom.plot.2 <- plot(fit.2, type = "regression", predictor =
sizes, reg.type ="RegScore")
allo.data.2 <- data.frame(gdf.2$CS,allom.plot.2$RegScore)

specimen.colors.2 <- c("#5EB16B","#4E99B7","#5EB16B","#4E99B7")
shapes.2 <- c(21, 21, 23, 23)
ggplot(data=allo.data.2, aes(x=gdf.2$CS, y=
allom.plot.2$RegScore, legend = F))+geom_point(shape =
shapes.2,size = 5, stroke= 0.1,color = "black",fill
=specimen.colors.2, alpha = 1)+scale_y_continuous(limits =
c(-0.2, 0.2))
theme(text = element_text(family = "Helvetica"))
+theme(axis.title.x = element_blank(),axis.text.x =
element_blank(),axis.text.y = element_blank(),axis.title.y =
element_blank(),axis.line = element_line(colour =
"black"),panel.grid.major = element_blank(),panel.background =
element_blank(),panel.grid.minor = element_blank())+
  geom_segment(aes(x = gdf.2$CS[2], y =
allom.plot.2$RegScore[2], xend = gdf.2$CS[4], yend =
allom.plot.2$RegScore[4]), linewidth = 1,color ='#4E99B7',alpha
= 0.09)+
  geom_segment(aes(x = gdf.2$CS[1], y =
allom.plot.2$RegScore[1], xend = gdf.2$CS[3], yend =
allom.plot.2$RegScore[3]), linewidth = 1,color ='#5EB16B',alpha
= 0.09)

########3d allo plot
array<- data.frame(gpa_1$Csize, pca$x[,1],pca$x[,2], age_1,
families_1)

mean.galli.hatch.pc1 <- mean(pca$x[Hatchling.Galliformes,1])
mean.galli.hatch.pc2 <- mean(pca$x[Hatchling.Galliformes,2])
mean.anseri.hatch.pc1 <- mean(pca$x[Hatchling.Anseriformes,1])
mean.anseri.hatch.pc2 <- mean(pca$x[Hatchling.Anseriformes,2])
mean.galli.adult.pc1 <- mean(pca$x[Adult.Galliformes,1])
mean.galli.adult.pc2 <- mean(pca$x[Adult.Galliformes,2])
mean.anseri.adult.pc1 <- mean(pca$x[Adult.Anseriformes,1])
mean.anseri.adult.pc2 <- mean(pca$x[Adult.Anseriformes,2])

plot_ly(array,x= ~log10(gpa_1$Csize),y=~pca$x[,1],z=~pca$x[,2],
marker = list(opacity
=0.8,line=list(width=1,color='black')),type = 'scatter3d',mode =
'markers',color =
as.factor(interaction(families_1,age_1)),colors=c("#4E99B7",
"#4EB773","#a6ccdb","#a6dbb9"))%>% layout(scene = list(xaxis =
list(title = 'Size', showbackground = TRUE, zerolinewidth =
18,backgroundcolor = "rgba(232,232,232,1)"), yaxis = list(title
```

```
= 'PC1 (56.34%)', showbackground = TRUE, backgroundcolor =
"rgba(211,211,211,1)"),zaxis = list(title = 'PC2 (9.89%)')))%>%
  add_trace(x = c(mean.galli.hatch.size,
mean.galli.adult.size),y = c(mean.galli.hatch.pc1,
mean.galli.adult.pc1), z = c(mean.galli.hatch.pc2,
mean.galli.adult.pc2),  type = 'scatter3d',mode = 'lines',line =
list(color = '#5EB16B',dash = 'longdashdot',width = 8),inherit =
FALSE, showlegend = F)%>%
  add_trace(x = c(mean.galli.hatch.size,
mean.galli.adult.size),y = c(mean.galli.hatch.pc1,
mean.galli.adult.pc1), z = c(rep(list(-0.1098726),2),
rep(list(-0.1098726),2)),  type = 'scatter3d',mode =
'lines',line = list(color = '#848688',width = 8),inherit =
FALSE, showlegend = F,opacity = 0.5)%>%
  add_trace(x = c(mean.anseri.hatch.size,
mean.anseri.adult.size),y = c(mean.anseri.hatch.pc1,
mean.anseri.adult.pc1), z = c(mean.anseri.hatch.pc2,
mean.anseri.adult.pc2),  type = 'scatter3d',mode = 'lines',line
= list(color = '#4E99B7',dash = 'longdashdot', width =
8),inherit = FALSE, showlegend = F)%>%
  add_trace(x = c(mean.anseri.hatch.size,
mean.anseri.adult.size),y = c(mean.anseri.hatch.pc1,
mean.anseri.adult.pc1), z = c(rep(list(-0.1098726),2),
rep(list(-0.1098726),2)),  type = 'scatter3d',mode =
'lines',line = list(color = '#848688', width = 8),inherit =
FALSE, opacity = 0.5, showlegend = F)%>%
  add_trace(x = ~log10(gpa_1$Csize), y = ~pca$x[,1], z =
~min(pca$x[,2])-0.01, inherit = F, mode = "markers", type =
"scatter3d", marker = list(size = 5, color = "#848688"), opacity
= 0.3)%>%
  add_trace(x = c(log10(gpa_1$Csize[1]),log10(gpa_1$Csize[1])),
y = c(list(pca$x[1,1]),list(pca$x[1,1])), z =
c(list(pca$x[1,2]),rep(list(-0.1098726),2)), inherit = F, mode =
"lines", type = "scatter3d",line = list(color =
'#848688'),showlegend = F, opacity = 0.5)%>%
  add_trace(x = c(log10(gpa_1$Csize[2]),log10(gpa_1$Csize[2])),
y = c(list(pca$x[2,1]),list(pca$x[2,1])), z =
c(list(pca$x[2,2]),rep(list(-0.1098726),2)), inherit = F, mode =
"lines", type = "scatter3d",line = list(color =
'#848688'),showlegend = F, opacity = 0.5)%>%
  add_trace(x = c(log10(gpa_1$Csize[3]),log10(gpa_1$Csize[3])),
y = c(list(pca$x[3,1]),list(pca$x[3,1])), z =
c(list(pca$x[3,2]),rep(list(-0.1098726),2)), inherit = F, mode =
"lines", type = "scatter3d",line = list(color =
'#848688'),showlegend = F, opacity = 0.5)%>%
  add_trace(x = c(log10(gpa_1$Csize[4]),log10(gpa_1$Csize[4])),
y = c(list(pca$x[4,1]),list(pca$x[4,1])), z =
```

```
c(list(pca$x[4,2]),rep(list(-0.1098726),2)), inherit = F, mode =
"lines", type = "scatter3d",line = list(color =
'#848688'),showlegend = F, opacity = 0.5)%>%
  add_trace(x = c(log10(gpa_1$Csize[5]),log10(gpa_1$Csize[5])),
y = c(list(pca$x[5,1]),list(pca$x[5,1])), z =
c(list(pca$x[5,2]),rep(list(-0.1098726),2)), inherit = F, mode =
"lines", type = "scatter3d",line = list(color =
'#848688'),showlegend = F, opacity = 0.5)%>%
  add_trace(x = c(log10(gpa_1$Csize[6]),log10(gpa_1$Csize[6])),
y = c(list(pca$x[6,1]),list(pca$x[6,1])), z =
c(list(pca$x[6,2]),rep(list(-0.1098726),2)), inherit = F, mode =
"lines", type = "scatter3d",line = list(color =
'#848688'),showlegend = F, opacity = 0.5)%>%
  add_trace(x = c(log10(gpa_1$Csize[7]),log10(gpa_1$Csize[7])),
y = c(list(pca$x[7,1]),list(pca$x[7,1])), z =
c(list(pca$x[7,2]),rep(list(-0.1098726),2)), inherit = F, mode =
"lines", type = "scatter3d",line = list(color =
'#848688'),showlegend = F, opacity = 0.5)%>%
  add_trace(x = c(log10(gpa_1$Csize[8]),log10(gpa_1$Csize[8])),
y = c(list(pca$x[8,1]),list(pca$x[8,1])), z =
c(list(pca$x[8,2]),rep(list(-0.1098726),2)), inherit = F, mode =
"lines", type = "scatter3d",line = list(color =
'#848688'),showlegend = F, opacity = 0.5)%>%
  add_trace(x = c(log10(gpa_1$Csize[9]),log10(gpa_1$Csize[9])),
y = c(list(pca$x[9,1]),list(pca$x[9,1])), z =
c(list(pca$x[9,2]),rep(list(-0.1098726),2)), inherit = F, mode =
"lines", type = "scatter3d",line = list(color =
'#848688'),showlegend = F, opacity = 0.5)%>%
  add_trace(x =
c(log10(gpa_1$Csize[10]),log10(gpa_1$Csize[10])), y =
c(list(pca$x[10,1]),list(pca$x[10,1])), z =
c(list(pca$x[10,2]),rep(list(-0.1098726),2)), inherit = F, mode
= "lines", type = "scatter3d",line = list(color =
'#848688'),showlegend = F, opacity = 0.5)%>%
  add_trace(x =
c(log10(gpa_1$Csize[11]),log10(gpa_1$Csize[11])), y =
c(list(pca$x[11,1]),list(pca$x[11,1])), z =
c(list(pca$x[11,2]),rep(list(-0.1098726),2)), inherit = F, mode
= "lines", type = "scatter3d",line = list(color =
'#848688'),showlegend = F, opacity = 0.5)%>%
  add_trace(x =
c(log10(gpa_1$Csize[12]),log10(gpa_1$Csize[12])), y =
c(list(pca$x[12,1]),list(pca$x[12,1])), z =
c(list(pca$x[12,2]),rep(list(-0.1098726),2)), inherit = F, mode
= "lines", type = "scatter3d",line = list(color =
'#848688'),showlegend = F, opacity = 0.5)%>%
  add_trace(x =
```

```
c(log10(gpa_1$Csize[13]),log10(gpa_1$Csize[13])), y =
c(list(pca$x[13,1]),list(pca$x[13,1])), z =
c(list(pca$x[13,2]),rep(list(-0.1098726),2)), inherit = F, mode
= "lines", type = "scatter3d",line = list(color =
'#848688'),showlegend = F, opacity = 0.5)%>%
  add_trace(x =
c(log10(gpa_1$Csize[14]),log10(gpa_1$Csize[14])), y =
c(list(pca$x[14,1]),list(pca$x[14,1])), z =
c(list(pca$x[14,2]),rep(list(-0.1098726),2)), inherit = F, mode
= "lines", type = "scatter3d",line = list(color =
'#848688'),showlegend = F, opacity = 0.5)%>%
  add_trace(x =
c(log10(gpa_1$Csize[15]),log10(gpa_1$Csize[15])), y =
c(list(pca$x[15,1]),list(pca$x[15,1])), z =
c(list(pca$x[15,2]),rep(list(-0.1098726),2)), inherit = F, mode
= "lines", type = "scatter3d",line = list(color =
'#848688'),showlegend = F, opacity = 0.5)%>%
  add_trace(x =
c(log10(gpa_1$Csize[16]),log10(gpa_1$Csize[16])), y =
c(list(pca$x[16,1]),list(pca$x[16,1])), z =
c(list(pca$x[16,2]),rep(list(-0.1098726),2)), inherit = F, mode
= "lines", type = "scatter3d",line = list(color =
'#848688'),showlegend = F, opacity = 0.5)%>%
  add_trace(x =
c(log10(gpa_1$Csize[17]),log10(gpa_1$Csize[17])), y =
c(list(pca$x[17,1]),list(pca$x[17,1])), z =
c(list(pca$x[17,2]),rep(list(-0.1098726),2)), inherit = F, mode
= "lines", type = "scatter3d",line = list(color =
'#848688'),showlegend = F, opacity = 0.5)%>%
  add_trace(x =
c(log10(gpa_1$Csize[18]),log10(gpa_1$Csize[18])), y =
c(list(pca$x[18,1]),list(pca$x[18,1])), z =
c(list(pca$x[18,2]),rep(list(-0.1098726),2)), inherit = F, mode
= "lines", type = "scatter3d",line = list(color =
'#848688'),showlegend = F, opacity = 0.5)%>%
  add_trace(x =
c(log10(gpa_1$Csize[19]),log10(gpa_1$Csize[19])), y =
c(list(pca$x[19,1]),list(pca$x[19,1])), z =
c(list(pca$x[19,2]),rep(list(-0.1098726),2)), inherit = F, mode
= "lines", type = "scatter3d",line = list(color =
'#848688'),showlegend = F, opacity = 0.5)%>%
  add_trace(x =
c(log10(gpa_1$Csize[20]),log10(gpa_1$Csize[20])), y =
c(list(pca$x[20,1]),list(pca$x[20,1])), z =
c(list(pca$x[20,2]),rep(list(-0.1098726),2)), inherit = F, mode
= "lines", type = "scatter3d",line = list(color =
'#848688'),showlegend = F, opacity = 0.5)%>%
```

```
  add_trace(x =
c(log10(gpa_1$Csize[21]),log10(gpa_1$Csize[21])), y =
c(list(pca$x[21,1]),list(pca$x[21,1])), z =
c(list(pca$x[21,2]),rep(list(-0.1098726),2)), inherit = F, mode
= "lines", type = "scatter3d",line = list(color =
'#848688'),showlegend = F, opacity = 0.5)%>%
  add_trace(x =
c(log10(gpa_1$Csize[22]),log10(gpa_1$Csize[22])), y =
c(list(pca$x[22,1]),list(pca$x[22,1])), z =
c(list(pca$x[22,2]),rep(list(-0.1098726),2)), inherit = F, mode
= "lines", type = "scatter3d",line = list(color =
'#848688'),showlegend = F, opacity = 0.5)%>%
  add_trace(x =
c(log10(gpa_1$Csize[23]),log10(gpa_1$Csize[23])), y =
c(list(pca$x[23,1]),list(pca$x[23,1])), z =
c(list(pca$x[23,2]),rep(list(-0.1098726),2)), inherit = F, mode
= "lines", type = "scatter3d",line = list(color =
'#848688'),showlegend = F, opacity = 0.5)%>%
  add_trace(x =
c(log10(gpa_1$Csize[24]),log10(gpa_1$Csize[24])), y =
c(list(pca$x[24,1]),list(pca$x[24,1])), z =
c(list(pca$x[24,2]),rep(list(-0.1098726),2)), inherit = F, mode
= "lines", type = "scatter3d",line = list(color =
'#848688'),showlegend = F, opacity = 0.5)%>%
  add_trace(x =
c(log10(gpa_1$Csize[25]),log10(gpa_1$Csize[25])), y =
c(list(pca$x[25,1]),list(pca$x[25,1])), z =
c(list(pca$x[25,2]),rep(list(-0.1098726),2)), inherit = F, mode
= "lines", type = "scatter3d",line = list(color =
'#848688'),showlegend = F, opacity = 0.5)%>%
  add_trace(x =
c(log10(gpa_1$Csize[26]),log10(gpa_1$Csize[26])), y =
c(list(pca$x[26,1]),list(pca$x[26,1])), z =
c(list(pca$x[26,2]),rep(list(-0.1098726),2)), inherit = F, mode
= "lines", type = "scatter3d",line = list(color =
'#848688'),showlegend = F, opacity = 0.5)%>%
  add_trace(x =
c(log10(gpa_1$Csize[27]),log10(gpa_1$Csize[27])), y =
c(list(pca$x[27,1]),list(pca$x[27,1])), z =
c(list(pca$x[27,2]),rep(list(-0.1098726),2)), inherit = F, mode
= "lines", type = "scatter3d",line = list(color =
'#848688'),showlegend = F, opacity = 0.5)%>%
  add_trace(x =
c(log10(gpa_1$Csize[28]),log10(gpa_1$Csize[28])), y =
c(list(pca$x[28,1]),list(pca$x[28,1])), z =
c(list(pca$x[28,2]),rep(list(-0.1098726),2)), inherit = F, mode
= "lines", type = "scatter3d",line = list(color =
```

```
'#848688'),showlegend = F, opacity = 0.5)%>%
  add_trace(x =
c(log10(gpa_1$Csize[29]),log10(gpa_1$Csize[29])), y =
c(list(pca$x[29,1]),list(pca$x[29,1])), z =
c(list(pca$x[29,2]),rep(list(-0.1098726),2)), inherit = F, mode
= "lines", type = "scatter3d",line = list(color =
'#848688'),showlegend = F, opacity = 0.5)%>%
  add_trace(x =
c(log10(gpa_1$Csize[30]),log10(gpa_1$Csize[30])), y =
c(list(pca$x[30,1]),list(pca$x[30,1])), z =
c(list(pca$x[30,2]),rep(list(-0.1098726),2)), inherit = F, mode
= "lines", type = "scatter3d",line = list(color =
'#848688'),showlegend = F, opacity = 0.5)%>%
  add_trace(x =
c(log10(gpa_1$Csize[31]),log10(gpa_1$Csize[31])), y =
c(list(pca$x[31,1]),list(pca$x[31,1])), z =
c(list(pca$x[31,2]),rep(list(-0.1098726),2)), inherit = F, mode
= "lines", type = "scatter3d",line = list(color =
'#848688'),showlegend = F, opacity = 0.5)%>%
  add_trace(x =
c(log10(gpa_1$Csize[32]),log10(gpa_1$Csize[32])), y =
c(list(pca$x[32,1]),list(pca$x[32,1])), z =
c(list(pca$x[32,2]),rep(list(-0.1098726),2)), inherit = F, mode
= "lines", type = "scatter3d",line = list(color =
'#848688'),showlegend = F, opacity = 0.5)%>%
  add_trace(x =
c(log10(gpa_1$Csize[33]),log10(gpa_1$Csize[33])), y =
c(list(pca$x[33,1]),list(pca$x[33,1])), z =
c(list(pca$x[33,2]),rep(list(-0.1098726),2)), inherit = F, mode
= "lines", type = "scatter3d",line = list(color =
'#848688'),showlegend = F, opacity = 0.5)%>%
  add_trace(x =
c(log10(gpa_1$Csize[34]),log10(gpa_1$Csize[34])), y =
c(list(pca$x[34,1]),list(pca$x[34,1])), z =
c(list(pca$x[34,2]),rep(list(-0.1098726),2)), inherit = F, mode
= "lines", type = "scatter3d",line = list(color =
'#848688'),showlegend = F, opacity = 0.5)%>%
  add_trace(x =
c(log10(gpa_1$Csize[35]),log10(gpa_1$Csize[35])), y =
c(list(pca$x[35,1]),list(pca$x[35,1])), z =
c(list(pca$x[35,2]),rep(list(-0.1098726),2)), inherit = F, mode
= "lines", type = "scatter3d",line = list(color =
'#848688'),showlegend = F, opacity = 0.5)%>%
  add_trace(x =
c(log10(gpa_1$Csize[36]),log10(gpa_1$Csize[36])), y =
c(list(pca$x[36,1]),list(pca$x[36,1])), z =
c(list(pca$x[36,2]),rep(list(-0.1098726),2)), inherit = F, mode
```

```
= "lines", type = "scatter3d",line = list(color =
'#848688'),showlegend = F, opacity = 0.5)%>%
  add_trace(x =
c(log10(gpa_1$Csize[37]),log10(gpa_1$Csize[37])), y =
c(list(pca$x[37,1]),list(pca$x[37,1])), z =
c(list(pca$x[37,2]),rep(list(-0.1098726),2)), inherit = F, mode
= "lines", type = "scatter3d",line = list(color =
'#848688'),showlegend = F, opacity = 0.5)%>%
  add_trace(x =
c(log10(gpa_1$Csize[38]),log10(gpa_1$Csize[38])), y =
c(list(pca$x[38,1]),list(pca$x[38,1])), z =
c(list(pca$x[38,2]),rep(list(-0.1098726),2)), inherit = F, mode
= "lines", type = "scatter3d",line = list(color =
'#848688'),showlegend = F, opacity = 0.5)%>%
  add_trace(x =
c(log10(gpa_1$Csize[39]),log10(gpa_1$Csize[39])), y =
c(list(pca$x[39,1]),list(pca$x[39,1])), z =
c(list(pca$x[39,2]),rep(list(-0.1098726),2)), inherit = F, mode
= "lines", type = "scatter3d",line = list(color =
'#848688'),showlegend = F, opacity = 0.5)%>%
  add_trace(x =
c(log10(gpa_1$Csize[40]),log10(gpa_1$Csize[40])), y =
c(list(pca$x[40,1]),list(pca$x[40,1])), z =
c(list(pca$x[40,2]),rep(list(-0.1098726),2)), inherit = F, mode
= "lines", type = "scatter3d",line = list(color =
'#848688'),showlegend = F, opacity = 0.5)%>%
  add_trace(x =
c(log10(gpa_1$Csize[41]),log10(gpa_1$Csize[41])), y =
c(list(pca$x[41,1]),list(pca$x[41,1])), z =
c(list(pca$x[41,2]),rep(list(-0.1098726),2)), inherit = F, mode
= "lines", type = "scatter3d",line = list(color =
'#848688'),showlegend = F, opacity = 0.5)%>%
  add_trace(x =
c(log10(gpa_1$Csize[42]),log10(gpa_1$Csize[42])), y =
c(list(pca$x[42,1]),list(pca$x[42,1])), z =
c(list(pca$x[42,2]),rep(list(-0.1098726),2)), inherit = F, mode
= "lines", type = "scatter3d",line = list(color =
'#848688'),showlegend = F, opacity = 0.5)%>%
  add_trace(x =
c(log10(gpa_1$Csize[43]),log10(gpa_1$Csize[43])), y =
c(list(pca$x[43,1]),list(pca$x[43,1])), z =
c(list(pca$x[43,2]),rep(list(-0.1098726),2)), inherit = F, mode
= "lines", type = "scatter3d",line = list(color =
'#848688'),showlegend = F, opacity = 0.5)%>%
  add_trace(x =
c(log10(gpa_1$Csize[44]),log10(gpa_1$Csize[44])), y =
c(list(pca$x[44,1]),list(pca$x[44,1])), z =
```

```
c(list(pca$x[44,2]),rep(list(-0.1098726),2)), inherit = F, mode
= "lines", type = "scatter3d",line = list(color =
'#848688'),showlegend = F, opacity = 0.5)%>%
  add_trace(x =
c(log10(gpa_1$Csize[45]),log10(gpa_1$Csize[45])), y =
c(list(pca$x[45,1]),list(pca$x[45,1])), z =
c(list(pca$x[45,2]),rep(list(-0.1098726),2)), inherit = F, mode
= "lines", type = "scatter3d",line = list(color =
'#848688'),showlegend = F, opacity = 0.5)%>%
  add_trace(x =
c(log10(gpa_1$Csize[46]),log10(gpa_1$Csize[46])), y =
c(list(pca$x[46,1]),list(pca$x[46,1])), z =
c(list(pca$x[46,2]),rep(list(-0.1098726),2)), inherit = F, mode
= "lines", type = "scatter3d",line = list(color =
'#848688'),showlegend = F, opacity = 0.5)%>%
  add_trace(x =
c(log10(gpa_1$Csize[47]),log10(gpa_1$Csize[47])), y =
c(list(pca$x[47,1]),list(pca$x[47,1])), z =
c(list(pca$x[47,2]),rep(list(-0.1098726),2)), inherit = F, mode
= "lines", type = "scatter3d",line = list(color =
'#848688'),showlegend = F, opacity = 0.5)%>%
  add_trace(x =
c(log10(gpa_1$Csize[48]),log10(gpa_1$Csize[48])), y =
c(list(pca$x[48,1]),list(pca$x[48,1])), z =
c(list(pca$x[48,2]),rep(list(-0.1098726),2)), inherit = F, mode
= "lines", type = "scatter3d",line = list(color =
'#848688'),showlegend = F, opacity = 0.5)%>%
  add_trace(x =
c(log10(gpa_1$Csize[49]),log10(gpa_1$Csize[49])), y =
c(list(pca$x[49,1]),list(pca$x[49,1])), z =
c(list(pca$x[49,2]),rep(list(-0.1098726),2)), inherit = F, mode
= "lines", type = "scatter3d",line = list(color =
'#848688'),showlegend = F, opacity = 0.5)%>%
  add_trace(x =
c(log10(gpa_1$Csize[50]),log10(gpa_1$Csize[50])), y =
c(list(pca$x[50,1]),list(pca$x[50,1])), z =
c(list(pca$x[50,2]),rep(list(-0.1098726),2)), inherit = F, mode
= "lines", type = "scatter3d",line = list(color =
'#848688'),showlegend = F, opacity = 0.5)%>%
  add_trace(x =
c(log10(gpa_1$Csize[51]),log10(gpa_1$Csize[51])), y =
c(list(pca$x[51,1]),list(pca$x[51,1])), z =
c(list(pca$x[51,2]),rep(list(-0.1098726),2)), inherit = F, mode
= "lines", type = "scatter3d",line = list(color =
'#848688'),showlegend = F, opacity = 0.5)%>%
  add_trace(x =
c(log10(gpa_1$Csize[52]),log10(gpa_1$Csize[52])), y =
```

```
c(list(pca$x[52,1]),list(pca$x[52,1])), z =
c(list(pca$x[52,2]),rep(list(-0.1098726),2)), inherit = F, mode
= "lines", type = "scatter3d",line = list(color =
'#848688'),showlegend = F, opacity = 0.5)%>%
  add_trace(x =
c(log10(gpa_1$Csize[53]),log10(gpa_1$Csize[53])), y =
c(list(pca$x[53,1]),list(pca$x[53,1])), z =
c(list(pca$x[53,2]),rep(list(-0.1098726),2)), inherit = F, mode
= "lines", type = "scatter3d",line = list(color =
'#848688'),showlegend = F, opacity = 0.5)%>%
  add_trace(x =
c(log10(gpa_1$Csize[54]),log10(gpa_1$Csize[54])), y =
c(list(pca$x[54,1]),list(pca$x[54,1])), z =
c(list(pca$x[54,2]),rep(list(-0.1098726),2)), inherit = F, mode
= "lines", type = "scatter3d",line = list(color =
'#848688'),showlegend = F, opacity = 0.5)%>%
  add_trace(x =
c(log10(gpa_1$Csize[55]),log10(gpa_1$Csize[55])), y =
c(list(pca$x[55,1]),list(pca$x[55,1])), z =
c(list(pca$x[55,2]),rep(list(-0.1098726),2)), inherit = F, mode
= "lines", type = "scatter3d",line = list(color =
'#848688'),showlegend = F, opacity = 0.5)%>%
  add_trace(x =
c(log10(gpa_1$Csize[56]),log10(gpa_1$Csize[56])), y =
c(list(pca$x[56,1]),list(pca$x[56,1])), z =
c(list(pca$x[56,2]),rep(list(-0.1098726),2)), inherit = F, mode
= "lines", type = "scatter3d",line = list(color =
'#848688'),showlegend = F, opacity = 0.5)%>%
  add_trace(x =
c(log10(gpa_1$Csize[57]),log10(gpa_1$Csize[57])), y =
c(list(pca$x[57,1]),list(pca$x[57,1])), z =
c(list(pca$x[57,2]),rep(list(-0.1098726),2)), inherit = F, mode
= "lines", type = "scatter3d",line = list(color =
'#848688'),showlegend = F, opacity = 0.5)%>%
  add_trace(x =
c(log10(gpa_1$Csize[58]),log10(gpa_1$Csize[58])), y =
c(list(pca$x[58,1]),list(pca$x[58,1])), z =
c(list(pca$x[58,2]),rep(list(-0.1098726),2)), inherit = F, mode
= "lines", type = "scatter3d",line = list(color =
'#848688'),showlegend = F, opacity = 0.5)%>%
  add_trace(x =
c(log10(gpa_1$Csize[59]),log10(gpa_1$Csize[59])), y =
c(list(pca$x[59,1]),list(pca$x[59,1])), z =
c(list(pca$x[59,2]),rep(list(-0.1098726),2)), inherit = F, mode
= "lines", type = "scatter3d",line = list(color =
'#848688'),showlegend = F, opacity = 0.5)%>%
  add_trace(x =
```

```
c(log10(gpa_1$Csize[60]),log10(gpa_1$Csize[60])), y =
c(list(pca$x[60,1]),list(pca$x[60,1])), z =
c(list(pca$x[60,2]),rep(list(-0.1098726),2)), inherit = F, mode
= "lines", type = "scatter3d",line = list(color =
'#848688'),showlegend = F, opacity = 0.5)


##########################################
####### Shape Disparity Analysis##########
##########################################

##### mean shape comparison
gdf <- geomorph.data.frame(coords = gpa_1$coords, CS=
gpa_1$Csize, age=age_1, taxons = taxons_1, families =
families_1 )
fit<-procD.lm(coords~ families*age, data = gdf, iter = 9999,
print.progress = F)
summary(fit)

##Interclade hatchlings
gdf2 <- geomorph.data.frame(coords = gpa_1$coords[,,age_1 ==
"hatchling"], CS= gpa_1$Csize[age_1 == "hatchling"], taxons =
taxons_1[age_1 == "hatchling"], families = families_1[age_1 ==
"hatchling"] )
fit2<-procD.lm( coords~ families, data = gdf2, iter = 9999,
print.progress = F)
anova(fit2)
PW2 <- pairwise(fit2, groups = families_1[age_1 == "hatchling"])
summary(PW2, stat.table = F)

fit2a <- procD.lm( CS~ families, data = gdf2, iter = 9999,
print.progress = F)
anova(fit2a)
PW2a <- pairwise(fit2a, groups = families_1[age_1 ==
"hatchling"])
summary(PW2a, stat.table = F)

##Interclade adults
gdf3 <- geomorph.data.frame(coords = gpa_1$coords[,,age_1 ==
"adult"], CS= gpa_1$Csize[age_1 == "adult"], taxons =
taxons_1[age_1 == "adult"], families = families_1[age_1 ==
"adult"] )
fit3<-procD.lm( coords~ families, data = gdf3, iter = 9999,
print.progress = F)
anova(fit3)
PW3 <- pairwise(fit3, groups = families_1[age_1 == "adult"])
summary(PW3, stat.table = F)
```

```
fit3a <- procD.lm( CS~ families, data = gdf3, iter = 9999,
print.progress = F)
anova(fit3a)
PW3a <- pairwise(fit3a, groups = families_1[age_1 == "adult"])
summary(PW3a, stat.table = F)

##within Galliformes
gdf4 <- geomorph.data.frame(coords = gpa_1$coords[,,families_1
== "Galliformes"], CS= gpa_1$Csize[families_1 == "Galliformes"],
taxons = taxons_1[families_1 == "Galliformes"], age =
age_1[families_1 == "Galliformes"],families =
families_1[families_1 == "Galliformes"] )
fit4<-procD.lm( coords~ age, data = gdf4, iter = 9999,
print.progress = F)
anova(fit4)
PW4 <- pairwise(fit4, groups = age_1[families_1 ==
"Galliformes"])
summary(PW4, stat.table = F)

##within Anseriformes

gdf5 <- geomorph.data.frame(coords = gpa_1$coords[,,families_1
== "Anseriformes"], CS= gpa_1$Csize[families_1 ==
"Anseriformes"], taxons = taxons_1[families_1 ==
"Anseriformes"], age = age_1[families_1 ==
"Anseriformes"],families = families_1[families_1 ==
"Anseriformes"] )
fit5<-procD.lm( coords~ age, data = gdf5, iter = 9999,
print.progress = F)
anova(fit5)
PW5 <- pairwise(fit5, groups = age_1[families_1 ==
"Anseriformes"])
summary(PW5, stat.table = F)

##between hatchlings of one clade and the adults of another

Hatchling.Galliformes<- intersect(which(age_1 ==
"hatchling"),which(families_1 == "Galliformes"))
Hatchling.Anseriformes<- intersect(which(age_1 ==
"hatchling"),which(families_1 == "Anseriformes"))
Adult.Galliformes<- intersect(which(age_1 ==
"adult"),which(families_1 == "Galliformes"))
Adult.Anseriformes<- intersect(which(age_1 ==
"adult"),which(families_1 == "Anseriformes"))

across1 <- c(Hatchling.Galliformes,Adult.Anseriformes)
```

```
gdf6 <- geomorph.data.frame(coords = gpa_1$coords[,,across1],
CS= gpa_1$Csize[across1], taxons = taxons_1[across1], age =
age_1[across1],families = families_1[across1] )
fit6<-procD.lm( coords~ age, data = gdf6, iter = 9999,
print.progress = F)
anova(fit6)
PW6 <- pairwise(fit6, groups = age_1[across1])
summary(PW6, stat.table = F)

across2<- c(Hatchling.Anseriformes,Adult.Galliformes)

gdf7 <- geomorph.data.frame(coords = gpa_1$coords[,,across2],
CS= gpa_1$Csize[across2], taxons = taxons_1[across2], age =
age_1[across2],families = families_1[across2] )
fit7<-procD.lm( coords~ age, data = gdf7, iter = 9999,
print.progress = F)
anova(fit7)
PW7 <- pairwise(fit7, groups = age_1[across2])
summary(PW7, stat.table = F)
##### shape variance comparison

disparity <- morphol.disparity(gpa_1$coords~1, groups =
interaction(families_1,age_1), iter=999)

#####Heatmaps for ontogenetic disparity

coords.adults <- gpa_1$coords[,,age_1== "adult"]
coords.hatchlings <- gpa_1$coords[,,age_1 == "hatchling"]

####### Distance matrices
Dist.Hatchs<-as.matrix(dist(two.d.array(coords.hatchlings)))
Dist.Adults<-as.matrix(dist(two.d.array(coords.adults)))

# Rename col and row names with Genus names

Names<- c("Anas", "Gallus","Dendrocygna", "Afropavo",
          "Pavo", "Aythya","Megapodius", "Cygnus",
          "Numida", "Malacorhynchus", "Anser",
          "Mergellus","Callonetta", "Rollulus","Oxyura",
          "Coturnix",  "Chauna", "Meleagris","Callipepla",
          "Lagopus","Aix","Alopochen","Tragopan", "Somateria",
          "Perdix", "Polyplectron","Mergus","Ortalis",
          "Spatula","Tadorna")
row.names(Dist.Hatchs)<- Names
colnames(Dist.Hatchs)<- Names

row.names(Dist.Adults)<- Names
```

```
colnames(Dist.Adults)<- Names

### Code for dendromgram
diag(Dist.Hatchs) <- 0 ## change the diagonal by 0 (if its not
the case)
diag(Dist.Adults) <- 0 ## change the diagonal by 0 (if its not
the case)
tree.galao <- read.tree("NeoTree.tree") # Open tree

#plot(tree.galao)
plot(tree.galao)

hc <- as.hclust(tree.galao) ## need to convert your tree file
into a cluster file for the plot
dend <- as.dendrogram(hc) ## convert it then to a dendogram
dd <- set(dend, "branches_lwd", 2) ## change the ldw of the tree
plot(dend, horiz=TRUE)

## Order the matrix as the phylogeny
mat_1<-as.matrix(Dist.Hatchs)
ord.mat_1 <- mat_1[tree.galao$tip.label,tree.galao$tip.label]
mat_2<-as.matrix(Dist.Adults)
ord.mat_2 <- mat_2[tree.galao$tip.label,tree.galao$tip.label]

##Make the heatmaps
coulb<-
colorRampPalette(c("gray66","#BDBDBD","#BDBDBD","#2B8CBE","#7BCC
C4","#CCEBC5","#FDBB84","#EF6548","#990000"))(25)
heatmap_1<- heatmap(ord.mat_1, Rowv=dd,
Colv=dd,scale="none",col=coulb)
heatmap_2<- heatmap(ord.mat_2, Rowv=dd,
Colv=dd,scale="none",col=coulb)

##make a legend
mid_1 <- (max(ord.mat_1)+min(ord.mat_2))/2
SpectrumLegend("bottomright", legend = c(max(ord.mat_1),mid_1,
min(ord.mat_1)),
               palette = coulb,
               inset = 0.1, # Inset from plot margin
               title = "Distances")

mid_2 <- (max(ord.mat_2)+min(ord.mat_2))/2
SpectrumLegend("bottomright", legend = c(max(ord.mat_2),mid_2,
min(ord.mat_2)),
               palette = coulb,
               inset = 0.1, # Inset from plot margin
               title = "Distances")
```

```
#############################################
#######Convergence/Divergence analysis########
##############################################

##### Using distances
##### 1. Distance matrix

Dist.Hatch.Ad<-as.matrix((dist(two.d.array(coords.hatchlings)))-
(dist(two.d.array(coords.adults))))

##### 2. Rename col and row names with Genus names
row.names(Dist.Hatch.Ad)<- c("Anas", "Gallus","Dendrocygna",
"Afropavo",
                             "Pavo", "Aythya","Megapodius",
"Cygnus",
                             "Numida", "Malacorhynchus",
"Anser",
                             "Mergellus","Callonetta",
"Rollulus","Oxyura",
                             "Coturnix",  "Chauna",
"Meleagris","Callipepla",

"Lagopus","Aix","Alopochen","Tragopan", "Somateria",
                             "Perdix",
"Polyplectron","Mergus","Ortalis",
                             "Spatula","Tadorna")

colnames(Dist.Hatch.Ad)<- c("Anas", "Gallus","Dendrocygna",
"Afropavo",
                            "Pavo", "Aythya","Megapodius",
"Cygnus",
                            "Numida", "Malacorhynchus", "Anser",
                            "Mergellus","Callonetta",
"Rollulus","Oxyura",
                            "Coturnix",  "Chauna",
"Meleagris","Callipepla",

"Lagopus","Aix","Alopochen","Tragopan", "Somateria",
                            "Perdix",
"Polyplectron","Mergus","Ortalis",
                            "Spatula","Tadorna")

####### 3. Extract the sign from the distance matrix

sign<-Dist.Hatch.Ad/abs(Dist.Hatch.Ad) # will be useful later to
```

```
mix distance sign with angle values

### Code for dendromgram
diag(Dist.Hatch.Ad) <- 0 ## change the diagonal by 0 (if its not
the case)

#plot(tree.galao)
plot(tree.galao)

hc <- as.hclust(tree.galao) ## need to convert your tree file
into a cluster file for the plot
dend <- as.dendrogram(hc) ## convert it then to a dendogram
dd <- set(dend, "branches_lwd", 2) ## change the ldw of the tree
plot(dend, horiz=TRUE)

## Order the matrix as the phylogeny
mat_3<-as.matrix(Dist.Hatch.Ad)
ord.mat_3 <- mat_3[tree.galao$tip.label,tree.galao$tip.label]

##Make the heatmaps
coulb<-
colorRampPalette(c("#1b5d80","#2478a3","#2B8CBE","#7BCCC4","#CCE
BC5","#BDBDBD","gray66","#f7bcb0","#e38b78"))(25)
heatmap_3<- heatmap(ord.mat_3, Rowv=dd,
Colv=dd,scale="none",col=coulb)

##make a legend
SpectrumLegend("bottomright", legend = c(max(ord.mat_3),'0',
min(ord.mat_3)),
               palette = coulb,
               inset = 0.001, # Inset from plot margin
               title = "Distances-based Convergence/Divergence")

##### Using angles
##### 1. Angle matrix

###Calculate angles

# Create matrix and empty matrix to be fill in the loop and
change the name of row and col

mat = matrix(nrow = 30, ncol = 30)

row.names(mat)<-c("Anas", "Gallus","Dendrocygna", "Afropavo",
                  "Pavo", "Aythya","Megapodius", "Cygnus",
                  "Numida", "Malacorhynchus", "Anser",
                  "Mergellus","Callonetta", "Rollulus","Oxyura",
```

```
                  "Coturnix",  "Chauna",
"Meleagris","Callipepla",
                  "Lagopus","Aix","Alopochen","Tragopan",
"Somateria",
                  "Perdix", "Polyplectron","Mergus","Ortalis",
                  "Spatula","Tadorna")
colnames(mat)<- c("Anas", "Gallus","Dendrocygna", "Afropavo",
                  "Pavo", "Aythya","Megapodius", "Cygnus",
                  "Numida", "Malacorhynchus", "Anser",
                  "Mergellus","Callonetta", "Rollulus","Oxyura",
                  "Coturnix",  "Chauna",
"Meleagris","Callipepla",
                  "Lagopus","Aix","Alopochen","Tragopan",
"Somateria",
                  "Perdix", "Polyplectron","Mergus","Ortalis",
                  "Spatula","Tadorna")

## Create a repetition that will used to do all the comparison
(Anas vs all the other then Gallus vs all the other etc....)
rep<-
c(rep("Anas",30),rep("Gallus",30),rep("Dendrocygna",30),rep("Afr
opavo",30),rep("Pavo",30),rep("Aythya",30),

rep("Megapodius",30),rep("Cygnus",30),rep("Numida",30),rep("Mala
corhynchus",30),rep("Anser",30),rep("Mergellus",30),

rep("Callonetta",30),rep("Rollulus",30),rep("Oxyura",30),rep("Co
turnix",30),rep("Chauna",30), rep("Meleagris",30),

rep("Callipepla",30),rep("Lagopus",30),rep("Aix",30),rep("Alopoc
hen",30),rep("Tragopan",30),rep("Somateria",30),

rep("Perdix",30),rep("Polyplectron",30),rep("Mergus",30),rep("Or
talis",30),rep("Spatula",30),rep("Tadorna",30))

df <- data.frame(row   = rep, col   = colnames(mat))
Angle_comparison2<-paste(df$row, df$col, sep="_")
comparisons2 <- as.vector(Angle_comparison2) # get the
comparisons we are going to need to match the other vectors
comparisons.split2 <- strsplit(comparisons2, "_") # this splits
the string of text in two items


# Vectors calculation and matrix filling


vec.angle <- function(v1,v2){
```

```
  acos((t(v1)%*%v2)/(sqrt(sum(v1^2))*sqrt(sum(v2^2))))*180/pi}
#function to calculate vector angles

coords.adults2 <- gpa_1$coords[,,age_1 == "adult"] ;
dimnames(coords.adults2) [[3]] <-
as.factor(row.names(Dist.Hatch.Ad))
coords.hatchlings2 <- gpa_1$coords[,,age_1 == "hatchling"] ;
dimnames(coords.hatchlings2) [[3]] <-
as.factor(row.names(Dist.Hatch.Ad))


Ang.mat = matrix(nrow = 30, ncol = 30) # Change the value in
function of the number of specimens (n-1 species)
row.names(Ang.mat)<- as.factor(row.names(Dist.Hatch.Ad))
colnames(Ang.mat)<- as.factor(row.names(Dist.Hatch.Ad))

y=1
x=1
z=1
for(z in 1:30){
  for(i in 1:30){
    taxon.1b <- comparisons.split2 [[x]] [[1]] # first part of
the string, first taxon
    taxon.2b <- comparisons.split2 [[x]] [[2]] # second part of
the string, second taxon

    x2<-coords.hatchlings2[,,taxon.1b]-
coords.adults2[,,taxon.1b]
    y2<-coords.hatchlings2[,,taxon.2b]-
coords.adults2[,,taxon.2b]

    X2<- as.vector(x2)/sqrt(sum(x^2))
    Y2<- as.vector(y2)/sqrt(sum(y^2))
    Ang.mat[i,z]<-vec.angle(X2,Y2)
    x=x+1

    if (i==30){
      i=1
      z=z+1

    }
    else{
      if (z==31) {
        print(Ang.mat[i,z])
      }
    }
  }
```

```
}

diag(Ang.mat) <- 0

## Order the matrix as the phylogeny
ord.mat_4 <- Ang.mat[tree.galao$tip.label,tree.galao$tip.label]

##Make the heatmaps
coulb<-
colorRampPalette(c("gray66","#BDBDBD","#BDBDBD","#2B8CBE","#7BCC
C4","#CCEBC5","#FDBB84","#EF6548","#990000"))(25)# Angle colors
mid <- (max(ord.mat_4)+min(ord.mat_4))/2
heatmap_4<- heatmap(ord.mat_4, Rowv=dd,
Colv=dd,scale="none",col=coulb)

##make a legend
SpectrumLegend("bottomright", legend = c(max(ord.mat_4),mid,
min(ord.mat_4)),
               palette = coulb,
               inset = 0.1, # Inset from plot margin
               title = "Angle-based Convergence/Divergence")

##### Using angles and signs

sign.mat <- Ang.mat*sign

diag(sign.mat) <- 0

## Order the matrix as the phylogeny
ord.mat_5 <- sign.mat[tree.galao$tip.label,tree.galao$tip.label]

##Make the heatmaps
coulb<-
colorRampPalette(c("#2B8CBE","#7BCCC4","#CCEBC5","#BDBDBD","gray
66","#BDBDBD","#EF6548","#990000"))(25)
heatmap_4<- heatmap(ord.mat_5, Rowv=dd,
Colv=dd,scale="none",col=coulb)

##make a legend
SpectrumLegend("bottomright", legend = c(max(ord.mat_5),'0',
min(ord.mat_5)),
               palette = coulb,
               inset = 0.1, # Inset from plot margin
               title = "Angle-based Convergence/Divergence")

########################################
########### Allometry analysis 2 #######
```

```
########################################

tree.temp <- tree.galao
#plots the scaled phylogeny

plot( tree.temp, direction = "u", cex=0.8, label.offset = 1,
edge.width = 3,
      no.margin=F, y.lim = c(-1,70), x.lim = c(-1, 30),
align.tip.label = T, srt = -45)

axisPhylo(side = 2)

#######Allometric tests functions

# Homogeneity of Slopes function code from Ollonen et al., 2022
to not log the centroid size, we do it afterwards
TestHOS <- function(shape, size, group, iter = 9, lm.fun) {

  hosgdf.gdf <- geomorph.data.frame(coords = shape, size = size,
spp = group)

  fit.unique <- lm.fun(coords ~ size * spp, iter = iter, data =
hosgdf.gdf, RRPP = FALSE)  # unique allometries by spp
  fit.common <- lm.fun(coords ~ size + spp, iter = iter, data =
hosgdf.gdf, RRPP = FALSE)  # common allometries across spp (NULL
HYPOTHESIS)
  pairwise(fit.unique, fit.common, groups = hosgdf.gdf$spp)

}
peram.test<-function(x,y,group,nperm=9999) {
  if (!is.factor(group)) stop("'group' must be a factor")
  if (!is.vector(x)) stop("'x' must be a vector")

  species<-as.numeric(group)
  fat_species<-group
  x<-x
  y<-as.matrix(y)

  dati=cbind(y,x,species)
  dati<-dati[order(group),]

  taglie=tapply(x,species,max)

  ### PERMUTATION PROCEDURE
  permute<-nperm
  PDiff_max_finale<-
matrix(0,nrow=max(species),ncol=max(species))
```

```
  Diff_obs_max<-matrix(0,nrow=max(species),ncol=max(species))
  x
  for(i in 1:(max(species)-1))
  {

    for(j in (i+1):max(species)){

      ################ per coppia di specie ###################

      dat=as.matrix(subset(as.data.frame(dati),species %in%
c(i,j)))

      ################ differenza intercetta osservata ########
      Y=dat[,1:ncol(y)]
      Csize=dat[,"x"]
      spe=dat[,"species"]

      model.fulli<-lm(Y[spe==i,]~Csize[spe==i])
      model.fullj<-lm(Y[spe==j,]~Csize[spe==j])

      vetti=c(1,taglie[i])
      vettj=c(1,taglie[j])

      full_maxi=as.numeric(crossprod(coef(model.fulli),vetti))
      full_maxj=as.numeric(crossprod(coef(model.fullj),vettj))

      obs.max=rbind(full_maxi,full_maxj)
      Dist.max.obs<-as.matrix((dist(obs.max)))

      ############## INIZIO PERMUTATION #####################


      PDiff_max<-array(1,dim=c(2,2))
      line<-nrow(Y)

      for(k in 1:permute){
        line.rand<-sample(line,replace=FALSE)
        res.temp<-cbind(line.rand,Y)
        z<-(order(line.rand))
        res.temp2<-as.matrix(res.temp[z,])
        Y.rand<-res.temp2[,-1]


        model.randi<-lm(Y.rand[spe==i,]~Csize[spe==i])
        model.randj<-lm(Y.rand[spe==j,]~Csize[spe==j])
```

```
        rand_maxi=as.numeric(crossprod(coef(model.randi),vetti))
        rand_maxj=as.numeric(crossprod(coef(model.randj),vettj))

        rand.max=rbind(rand_maxi,rand_maxj)

        Dist.max.rand<-as.matrix((dist(rand.max)))


        PDiff_max<-
ifelse(Dist.max.rand>=Dist.max.obs,PDiff_max+1,PDiff_max)


      }  #end permute

      PDiff_max<-PDiff_max/(permute+1)
      ############# FINE PERMUTATION ########################

      Diff_obs_max[i,j]<-Dist.max.obs[1,2]

      PDiff_max_finale[i,j]<-PDiff_max[1,2]

    }
    rownames(PDiff_max_finale)=colnames(PDiff_max_finale)<-
levels(group)
    rownames(Diff_obs_max)=colnames(Diff_obs_max)<-levels(group)

  }


  PG<-list("p.value"=PDiff_max_finale,"obs.diff"=Diff_obs_max)
  PG

} # peramorphosis test
int.test<-function(x,y,x.value=0,group,nperm=1000) {
  library(MASS)
  if (!is.factor(group)) stop("'group' must be a factor")
  if (!is.vector(x)) stop("'x' must be a vector")

  species<-as.numeric(group)
  fat_species<-group
  x<-x
  y<-as.matrix(y)

  dati=cbind(y,x,species)
  dati<-dati[order(group),]

  ### PERMUTATION PROCEDURE
```

```
  PDiff_finale<-matrix(0,nrow=max(species),ncol=max(species))
  Diff_obs<-matrix(0,nrow=max(species),ncol=max(species))

  vett=c(1,x.value)
  for(i in 1:(max(species)-1))
  {
    for(j in (i+1):max(species)){
      ################ pairwise ###################
      dat=as.matrix(subset(as.data.frame(dati),species %in%
c(i,j)))

      Y=as.matrix(dat[,1:ncol(y)])
      Csize=dat[,(ncol(y)+1):(ncol(dat)-1)]
      spe=dat[,ncol(dat)]

      model.fulli<-lm(Y[spe==i,]~Csize[spe==i])
      model.fullj<-lm(Y[spe==j,]~Csize[spe==j])

      full_inti=as.numeric(crossprod(coef(model.fulli),vett))
      full_intj=as.numeric(crossprod(coef(model.fullj),vett))

      obs.int=rbind(full_inti,full_intj)
      Dist.int.obs<-as.matrix((dist(obs.int)))
      ############## START PERMUTATION #####################
      PDiff<-array(1,dim=c(2,2))
      line<-nrow(Y)

      for(k in 1:nperm){
        line.rand<-sample(line,replace=FALSE)
        res.temp<-cbind(line.rand,Y)
        z<-(order(line.rand))
        res.temp2<-as.matrix(res.temp[z,])
        Y.rand<-as.matrix(res.temp2[,-1])

        model.randi<-lm(Y.rand[spe==i,]~Csize[spe==i])
        model.randj<-lm(Y.rand[spe==j,]~Csize[spe==j])

        rand_inti=as.numeric(crossprod(coef(model.randi),vett))
        rand_intj=as.numeric(crossprod(coef(model.randj),vett))

        rand.int=rbind(rand_inti,rand_intj)

        Dist.int.rand<-as.matrix((dist(rand.int)))

        PDiff<-ifelse(Dist.int.rand>=Dist.int.obs,PDiff+1,PDiff)
      }  #end permute
      PDiff<-PDiff/(nperm+1)
```

```
      ############# END PERMUTATION ########################
      Diff_obs[i,j]<-Dist.int.obs[1,2]
      PDiff_finale[i,j]<-PDiff[1,2]

    }
    rownames(PDiff_finale)=colnames(PDiff_finale)<-levels(group)
    rownames(Diff_obs)=colnames(Diff_obs)<-levels(group)
  }
  PG<-list("p.value"=PDiff_finale,"obs.diff"=Diff_obs)
  PG
} # intercept test

#### CLADE-LEVEL COMPARISONS ####

###### Analyses only between galliforms and anseriforms ######

#homogeneity of slopes

Test.HOS.temp <- TestHOS(gpa_1$coords, log10(gpa_1$Csize), group
= families_1, lm.fun = procD.lm)
Pairwise.Test.HOS.temp <- summary.pairwise(Test.HOS.temp,
test.type = "DL", confidence = 0.95)
Pairwise.Test.HOS.temp         #differences in slope (i.e., rate
of shape change per unit of size change)

Test.HOS.temp$means.length #allometric slopes per species

#same slopes between anseriforms and galliforms in general

#intercept test

intercept.results.species <- int.test(log10(gpa_1$Csize),
gpa_1$coords, -0.7, families_1, nperm=10000)
intercept.results.species

#same intercept between anseriforms and galliforms in general


#peramorphism test

peratest <- peram.test(log10(gpa_1$Csize),
two.d.array(gpa_1$coords), families_1)
peratest


#at the clade level it seems like Anseriformes show peramorphic
ontogenies with respect to Galliformes
```

```
#subset the taxa to compare Anatidae versus Phasianidae

outgroups <- c("Numida.hatchling", "Numida.adult",
               "Megapodius.hatchling", "Megapodius.adult",
               "Chauna.hatchling", "Chauna.adult",
               "Callipepla.hatchling", "Callipepla.adult",
               "Ortalis.hatchling", "Ortalis.adult")


coords.less.inclusive <- gpa_1$coords [,, !
dimnames(gpa_1$coords) [[3]]  %in% outgroups ]
csize.less.inclusive <- gpa_1$Csize [!names(gpa_1$Csize) %in%
outgroups]

#check if the lengths match

dim(coords.less.inclusive)
length(csize.less.inclusive)


#this is a new category that only has "Anatidae" and
"Phasianidae", do not include more categories.

families.less.inclusive <- as.factor(c("Anatidae", "Anatidae",
"Phasianidae", "Phasianidae",
                                       "Anatidae", "Anatidae",
"Phasianidae", "Phasianidae",
                                       "Phasianidae",
"Phasianidae", "Anatidae", "Anatidae",
                                       "Anatidae", "Anatidae",
"Anatidae", "Anatidae",
                                       "Anatidae", "Anatidae",
"Anatidae", "Anatidae",
                                       "Anatidae","Anatidae",
"Phasianidae", "Phasianidae",
                                       "Anatidae","Anatidae",
"Phasianidae", "Phasianidae",
                                       "Phasianidae",
"Phasianidae","Phasianidae","Phasianidae",

"Anatidae","Anatidae","Anatidae","Anatidae",
                                       "Phasianidae",
"Phasianidae", "Anatidae", "Anatidae",

"Phasianidae","Phasianidae","Phasianidae", "Phasianidae",
```

```
                                       "Anatidae","Anatidae",
"Anatidae","Anatidae",
                                       "Anatidae","Anatidae"))
#ages for less.inclusive

age_4<-
c("adult","hatchling","adult","hatchling","adult","hatchling","h
atchling","adult","hatchling","adult","hatchling","adult",

"hatchling","adult","hatchling","adult","hatchling","adult","hat
chling","adult","adult","hatchling","hatchling","adult",

"hatchling","adult","adult","hatchling","hatchling","adult","hat
chling","adult","hatchling","adult","hatchling","adult",

"hatchling","adult","hatchling","adult","hatchling","adult","hat
chling","adult","hatchling","adult","hatchling","adult",
          "hatchling","adult")

neudisparity <- morphol.disparity(coords.less.inclusive~1,
groups = interaction(families.less.inclusive,age_4), iter=999)
#check that the length matches

length(families.less.inclusive)

#homogeneity of slopes

Test.HOS.temp <- TestHOS(coords.less.inclusive,
log10(csize.less.inclusive), group = families.less.inclusive,
lm.fun = procD.lm)
Pairwise.Test.HOS.temp <- summary.pairwise(Test.HOS.temp,
test.type = "DL", confidence = 0.95)
Pairwise.Test.HOS.temp
#differences in slope (i.e., rate of shape change per unit of
size change)

#same slopes between anatids and phasianids in general


#intercept test

intercept.results.species <-
int.test(log10(csize.less.inclusive), coords.less.inclusive,
-0.7, families.less.inclusive, nperm=10000)
intercept.results.species

#same intercept between between anatids and phasianids in
```

```
general


#peramorphism test

peratest <- peram.test(log10(csize.less.inclusive),
two.d.array(coords.less.inclusive), families.less.inclusive)
peratest


#at the clade level it seems like anatids show peramorphic
ontogenies with respect to phasianids

############## SPECIES COMPARISONS ##############


# Homogeneity of slopes test # for all the species
# we do not have enough sample to trust the species-level tests
# instead we will characterise the species-level ontogenies in
various ways


## INDIVIDUAL VALUES FROM SPECIES-LEVEL ONTOGENIES ##

# First, we will extract individual metrics from all the
different species-level ontogenies
# and will summarise their variation as contMaps and PCA-plots

#get the species level slopes using HOS

Test.HOS.temp <- TestHOS(gpa_1$coords, log10(gpa_1$Csize), group
= taxons_1, lm.fun = procD.lm)
Pairwise.Test.HOS.temp <- summary.pairwise(Test.HOS.temp,
test.type = "DL", angle.type = "deg", confidence = 0.99)
#differences in slope (i.e., rate of shape change per unit of
size change)


species.allometric.slopes <-  Test.HOS.temp$means.length$obs
#allometric slopes per species



# Calculate total ontogenetic changes in size and shape

variable = two.d.array(gpa_1$coords)
variable = log10(gpa_1$Csize)
```

```
#log10(gpa$Csize) # use this for calculating changes in centroid
size

#distances among all pairs of taxa

distances <- as.matrix(dist(variable))

#subset the relevant comparisons
#the comparisons between hatchling and adult of the same taxon
are below the diagonal of the matrix of distances

distances.temp <- distances[1 + row(distances)==col(distances)];
names(distances.temp)<- row.names(distances)
[2:length(row.names(distances))]

#the appropriate comparisons are the odd numbers of the vector

numbers<-c(1:length(distances.temp))

odd_numbers <- numbers[numbers %% 2 != 0]

distances.temp.final <- distances.temp[odd_numbers ] ;
names(distances.temp.final) <- taxons_1[odd_numbers]

#Stores the values for species ontogenetic change in centroid
size (log10-transformed) and shape

species.shape.variation <- distances.temp.final
species.logCS.variation <- distances.temp.final

species.allometric.slopes.new <- species.shape.variation /
species.logCS.variation


# calculate c.size (log10-transformed) adults

species.logCS.hatchlings <- gpa_1$Csize [age_1 == "hatchling"];
names(species.logCS.hatchlings) <-
names(species.shape.variation)
species.logCS.adults <- gpa_1$Csize [age_1 == "adult"];
names(species.logCS.adults) <- names(species.shape.variation)
```

```
#combines all the species ontogenetic values into a matrix

#normalise all values to range 0-1

range01 <- function(x){(x-min(x))/(max(x)-min(x))} # normalises
ranges among all clades

n.species.allometric.slopes <-
range01(species.allometric.slopes.new)
n.species.logCS.variation <-
range01(species.logCS.variation[names(n.species.allometric.slope
s)])
n.species.shape.variation <-
range01(species.shape.variation[names(n.species.allometric.slope
s)])
n.species.logCS.hatchlings <-
range01(species.logCS.hatchlings[names(n.species.allometric.slop
es)])
n.species.logCS.adults <-
range01(species.logCS.adults[names(n.species.allometric.slopes)]
)


#calculates a PCA plot with all the species ontogeny metrics
(slopes, logCS variation, shape variation, logCS at hatching and
logCS as adult)

#make sure all the variables are ordered in the same way as
abind() would not check if that criterion is met

individual.onto.traits <-
cbind( n.species.allometric.slopes,n.species.logCS.variation,
n.species.shape.variation,
                                 n.species.logCS.hatchlings,
n.species.logCS.adults)


individual.onto.traits<-as.data.frame(individual.onto.traits)
[tree.temp$tip.label,]

species.onto.space <- prcomp(individual.onto.traits)
x<-summary(species.onto.space)

print(row.names(individual.onto.traits))

colours.phylo.onto.space <- c("#4EB773",
"#4EB773","#4EB773","#4EB773","#4EB773",
```

```
"#4EB773","#4EB773","#4EB773","#4EB773","#4EB773",

"#4EB773","#4EB773","#4EB773","#4EB773","#4E99B7",
                              "#4E99B7", "#4E99B7", "#4E99B7",
"#4E99B7", "#4E99B7",
                              "#4E99B7", "#4E99B7", "#4E99B7",
"#4E99B7", "#4E99B7",
                              "#4E99B7", "#4E99B7", "#4E99B7",
"#4E99B7", "#4E99B7")
#  "#4E99B7", "#4E99B7", "#4EB773", "#4E99B7",
# "#4E99B7", "#4EB773", "#4E99B7", "#4E99B7",
#"#4EB773", "#4EB773", "#4E99B7", "#4EB773",
#"#4EB773", "#4E99B7", "#4EB773", "#4E99B7",
#"#4EB773", "#4EB773", "#4EB773", "#4EB773",
#"#4E99B7", "#4EB773"




plot(species.onto.space$x[,1], species.onto.space$x[,2],
     xlab = paste("PC1", "(", round(100*x$importance[2,1],
digits = 2), "%", ")"),
     ylab = paste("PC2", "(", round(100*x$importance[2,2],
digits = 2), "%", ")") )

phylomorphospace(tree.temp,species.onto.space$x[,
c(1,2)],ftype="off", add=TRUE, node.size=0)
points(species.onto.space$x[,1], species.onto.space$x[,2],
col=colours.phylo.onto.space, pch=16, cex=3)

text(species.onto.space$x[,1], species.onto.space$x[,2], labels
= rownames(individual.onto.traits),
     cex = 0.5, offset = 5)

biplot(species.onto.space,choices = c(1,2), col=c("transparent",
'red'), cex = 0.5)

#plots individual ontogenetic traits: the evolution of total
shape change, total size change and allometric slopes in the
phylogeny

#species.shape.variation
#species.logCS.variation
#species.allometric.slopes.new
#species.logCS.hatchlings
```

```
#species.logCS.adults

##############################################
#test differences in ontogenetic shape and size change

families1 <- c("Anseriformes", "Anseriformes", "Galliformes",
"Galliformes", "Anseriformes",
               "Anseriformes", "Galliformes", "Galliformes",
"Galliformes", "Galliformes",
               "Anseriformes", "Anseriformes", "Galliformes",
"Galliformes", "Anseriformes",
               "Anseriformes", "Galliformes",
"Galliformes","Anseriformes","Anseriformes",
               "Anseriformes", "Anseriformes", "Anseriformes",
"Anseriformes", "Anseriformes",
               "Anseriformes", "Galliformes", "Galliformes",
"Anseriformes","Anseriformes",
               "Galliformes",
"Galliformes","Anseriformes","Anseriformes", "Galliformes",
               "Galliformes", "Galliformes", "Galliformes",
"Galliformes", "Galliformes",

"Anseriformes","Anseriformes","Anseriformes","Anseriformes",
"Galliformes",
               "Galliformes","Anseriformes","Anseriformes",
"Galliformes", "Galliformes",
               "Galliformes",
"Galliformes","Anseriformes","Anseriformes", "Galliformes",

"Galliformes","Anseriformes","Anseriformes","Anseriformes","Anse
riformes")
families1 <- families1[-
c(2,4,6,8,10,12,14,16,18,20,22,24,26,28,30,32,34,36,38,40,42,44,
46,48,50,52,54,56,58,60)]

mean(species.shape.variation[families1=="Anseriformes"])
##calcuate the mean shape vector length for Anseriformes
mean(species.shape.variation[families1=="Galliformes"])
##calcuate the mean shape vector length for Galliformes

gdf5 <- geomorph.data.frame(data = species.shape.variation,
families = families1 )
fit5<-procD.lm( data~ families, data = gdf5, iter = 9999,
print.progress = F)
anova(fit5)
PW5 <- pairwise(fit5, groups = families1)
summary(PW5, stat.table = F)
```

```
morphol.disparity(species.shape.variation~1, groups =
families1 , iter=999) ## calculate the variance in shape vector
length in both clades

mean(species.logCS.variation[families1=="Anseriformes"])
##calcuate the mean quantity of ontogenetic size change for
Anseriformes
mean(species.logCS.variation[families1=="Galliformes"])
##calcuate the mean quantity of ontogenetic size change for
Galliformes

gdf6 <- geomorph.data.frame(data = species.logCS.variation,
families = families1 )
fit6<-procD.lm( data~ families, data = gdf6, iter = 9999,
print.progress = F)
anova(fit6)
PW6 <- pairwise(fit6, groups = families1)
summary(PW6, stat.table = F)

morphol.disparity(species.logCS.variation~1, groups =
families1 , iter=999) ## calculate the variance in ontogenetic
size change in both clades
################################

variable <- species.shape.variation

tree <- tree.temp
x<-variable
names(x)<-names(variable)
x <- x[tree1$tip.label]

drop.taxa <- tree$tip.label[ ! tree$tip.label %in% names(x) ]
drop.taxa # check the taxa are the expected to be drawn out


phylo.IREmod<-contMap(tree,
                      x,
                      res=100,
                      lwd=4,
                      legend = 40,
                      outline=FALSE,
                      sig=3,
                      type="phylogram",
                      direction="upwards",
                      plot=F)
```

```
obj<-setMap(phylo.IREmod,colors=c("#5BACE8", "#A3CCE5",
"#E5C69E", "#E8B15B", "#E5765C"))

plot(obj, fsize=0.8,lwd = 5, outline = FALSE, direction =
"upwards",
     no.margin=T, ylim = c(-5,65), xlim = c(1, 30),
align.tip.label = T, mar=c(5.1, 4.1, 4.1, 2.1))

axisPhylo(side = 2)


#############trees with tips only
x<-gpa_1$Csize[age_1=="hatchling"]
unique(taxons_1)[2]
"Gallus" <-


  ggtree(tree1)+geom_tippoint(aes(data = x,colour = x))
+scale_colour_gradient(low='#3399FF', high='#CC3333')
+geom_tiplab()+theme(legend.position = "bottom")

## PAIRWISE DIFFERENCES AMONG SPECIES-LEVEL ONTOGENIES ##

# New pairwise differences in vector slopes

species.allometric.slopes.new <- species.allometric.slopes.new
[taxons.alphabetical]

delta.slope.vector.lengths <-
as.matrix(dist(species.allometric.slopes.new, upper = F))



delta <- c(1:length(comparisons))
names(delta) <- comparisons
delta


comparisons <- names(delta.slope.vector.angles) # get the
comparisons we are going to need to match the other vectors

comparisons.split <- strsplit(comparisons, ":") # this splits
the string of text in two items


for( i in 1:length( delta ) ) {
```

```
  taxon.1 <- comparisons.split [[i]] [[1]] # first part of the
string, first taxon
  taxon.2 <- comparisons.split [[i]] [[2]] # second part of the
string, second taxon

  taxon.1
  taxon.2

  delta [[i]] <- delta.slope.vector.lengths [taxon.1,taxon.2] #
subsets the appropriate comparison among taxa




}


delta
delta.slope.vector.lengths <- delta



delta.size.stage ["Anas","Coturnix"]  # check it worked
comparing original values


write.csv(delta.size.stage.final,
"delta.slope.vector.lengths.csv", row.names = T)



# Pairwise differences in vector slope angles


delta.slope.vector.angles <-
Pairwise.Test.HOS.temp$summary.table[, 2] ; names
(delta.slope.vector.angles) <-
row.names(Pairwise.Test.HOS.temp$summary.table)

delta.slope.vector.angles



#Pairwise differences in shape at hatchling stage
```

}

```
shape.hatchlings <- two.d.array(gpa$coords[,,age ==
"hatchling"]) ; rownames(shape.hatchlings) <-
taxons[odd_numbers]

taxons.alphabetical <- as.character(sort(taxons_1[odd_numbers]))

shape.hatchlings <- shape.hatchlings [taxons.alphabetical,]

rownames(shape.hatchlings)

delta.shape.hatchlings <- as.matrix(dist(shape.hatchlings, upper
= F))



delta <- c(1:length(comparisons))
names(delta) <- comparisons
delta


comparisons <- names(delta.slope.vector.angles) # get the
comparisons we are going to need to match the other vectors

comparisons.split <- strsplit(comparisons, ":") # this splits
the string of text in two items


for( i in 1:length( delta ) ) {



  taxon.1 <- comparisons.split [[i]] [[1]] # first part of the
string, first taxon
  taxon.2 <- comparisons.split [[i]] [[2]] # second part of the
string, second taxon

  taxon.1
  taxon.2

  delta [[i]] <- delta.shape.hatchlings [taxon.1,taxon.2] #
subsets the appropriate comparison among taxa





}
```

```
delta
delta.shape.hatchlings.final <- delta


delta.shape.hatchlings ["Rollulus","Tragopan"]  # check it
worked comparing original values


write.csv(delta.shape.hatchlings.final,
"delta.shape.hatchlings.final.csv", row.names = T)


#Pairwise differences in shape at adult stage


shape.adults <- two.d.array(gpa$coords[,,age == "adult"]) ;
rownames(shape.adults) <- taxons[odd_numbers]

taxons.alphabetical <- as.character(sort(taxons[odd_numbers]))

shape.adults <- shape.adults [taxons.alphabetical,]

rownames(shape.adults)

delta.shape.adults <- as.matrix(dist(shape.adults, upper = F))



delta <- c(1:length(comparisons))
names(delta) <- comparisons
delta


comparisons <- names(delta.slope.vector.angles) # get the
comparisons we are going to need to match the other vectors

comparisons.split <- strsplit(comparisons, ":") # this splits
the string of text in two items


for( i in 1:length( delta ) ) {



  taxon.1 <- comparisons.split [[i]] [[1]] # first part of the
string, first taxon
```

```
  taxon.2 <- comparisons.split [[i]] [[2]] # second part of the
string, second taxon

  taxon.1
  taxon.2

  delta [[i]] <- delta.shape.adults [taxon.1,taxon.2] # subsets
the appropriate comparison among taxa




}


delta
delta.shape.adults.final <- delta


delta.shape.adults ["Rollulus","Tragopan"]  # check it worked
comparing original values


write.csv(delta.shape.adults.final,
"delta.shape.adults.final.csv", row.names = T)


#Pairwise differences in size at adults and hatchlings

ontogenetic.stage = "hatchling"


size.stage <- log10(gpa$Csize[age == ontogenetic.stage]) ;
names(size.stage) <- taxons[odd_numbers]

taxons.alphabetical <- as.character(sort(taxons[odd_numbers]))

size.stage <- size.stage [taxons.alphabetical]

names(size.stage)

delta.size.stage <- as.matrix(dist(size.stage, upper = F))



delta <- c(1:length(comparisons))
names(delta) <- comparisons
```

```
delta


comparisons <- names(delta.slope.vector.angles) # get the
comparisons we are going to need to match the other vectors

comparisons.split <- strsplit(comparisons, ":") # this splits
the string of text in two items


for( i in 1:length( delta ) ) {



  taxon.1 <- comparisons.split [[i]] [[1]] # first part of the
string, first taxon
  taxon.2 <- comparisons.split [[i]] [[2]] # second part of the
string, second taxon

  taxon.1
  taxon.2

  delta [[i]] <- delta.size.stage [taxon.1,taxon.2] # subsets
the appropriate comparison among taxa




}


delta
delta.size.stage.final <- delta



delta.size.stage ["Rollulus","Tragopan"]  # check it worked
comparing original values


write.csv(delta.size.stage.final,
"delta.size.hatchling.final.csv", row.names = T)




#Read these two final lines to store the size data
```

```
delta.size.hatchlings.final<- delta.size.stage.final

#delta.size.adults.final<- delta.size.stage.final





## Differences in angle between shape trajectories among species
##


delta <- c(1:length(comparisons))
names(delta) <- comparisons
delta

comparisons <- names(delta.slope.vector.angles) # get the
comparisons we are going to need to match the other vectors

comparisons.split <- strsplit(comparisons, ":") # this splits
the string of text in two items



vec.angle <- function(v1,v2){
  acos((t(v1)%*%v2)/(sqrt(sum(v1^2))*sqrt(sum(v2^2))))*180/pi}
#function to calculate vector angles





for( i in 1:length( delta ) ) {



  taxon.1 <- comparisons.split [[i]] [[1]] # first part of the
string, first taxon
  taxon.2 <- comparisons.split [[i]] [[2]] # second part of the
string, second taxon

  taxon.1
  taxon.2

  coords.adults <- gpa$coords[,,age == "adult"] ;
```

```
dimnames(coords.adults) [[3]] <- taxons[odd_numbers]
  coords.hatchlings <- gpa$coords[,,age == "hatchling"] ;
dimnames(coords.hatchlings) [[3]] <- taxons[odd_numbers]


  x<-coords.hatchlings[,,taxon.1]-coords.adults[,,taxon.1]
  y<-coords.hatchlings[,,taxon.2]-coords.adults[,,taxon.2]

  X <- as.vector(x)/sqrt(sum(x^2))
  Y <- as.vector(y)/sqrt(sum(y^2))



  angle.temp <- vec.angle(X,Y)
  angle.temp


  delta [[i]] <- angle.temp # subsets the appropriate comparison
among taxa



}


delta.shape.angles <- delta



#### ONTO-PCA FOR DIFFERENCES BETWEEN PAIRS OF TAXA #####


# colours for comparisons

colours.comparisons <-
read.csv("categories.comparisons.final.csv", row.names = 1) #
this is a list that was done outside of R and imported
indicating which is a inter and intraclade comparison

colours.1 <- as.character(colours.comparisons [,
"comparisons.taxonomy"])

colours.1.1 <- gsub("1", "#B78A4E", colours.1)
colours.1.2 <- gsub( "2", "#4E99B7", colours.1.1)
colours.1.3 <- gsub("3", "#4EB773", colours.1.2)

colours.1.3
```

```
colours.2 <- as.character(colours.comparisons [,
"comparisons.less.inclusive"])

colours.2.1 <- gsub("1", "#B78A4E", colours.2)
colours.2.2 <- gsub( "2", "#4E99B7", colours.2.1)
colours.2.3 <- gsub("3", "#4EB773", colours.2.2)

colours.2.4 <- gsub("%&", "#926FBF", colours.2.3) # chauna

colours.2.4


#bind all the comparisons


n.delta.slope.vector.lengths <-
range01(delta.slope.vector.lengths)
n.delta.slope.vector.angles <-
range01(delta.slope.vector.angles)
n.delta.shape.adults.final <- range01(delta.shape.adults.final)
n.delta.shape.hatchlings.final <-
range01(delta.shape.hatchlings.final)
n.delta.size.adults.final <- range01(delta.size.adults.final)
n.delta.size.hatchlings.final <-
range01(delta.size.hatchlings.final)
n.delta.shape.angles <- range01(delta.shape.angles)



delta.onto.traits <- cbind(n.delta.slope.vector.lengths,
n.delta.slope.vector.angles,
                           n.delta.shape.adults.final,
n.delta.shape.hatchlings.final,
                           n.delta.size.adults.final,
n.delta.size.hatchlings.final,
                           n.delta.shape.angles)

delta.onto.traits.no.chauna <- delta.onto.traits [!colours.2 ==
"%&",] #take out Chauna

colours.2.4 <- colours.2.4 [!colours.2 == "%&"] # colours for
subset without Chauna
```

```
delta.onto.space <- prcomp(delta.onto.traits)
x<-summary(delta.onto.space)

biplot(delta.onto.space,choices = c(1,3), col=c("transparent",
'red'), cex = 0.5)

plot(delta.onto.space$x[,1], delta.onto.space$x[,2], cex = 2,
col = "transparent", pch = 21, bg = colours.1.3, frame.plot = T,
xlab = paste("PC1", "(", round(100*x$importance[2,1], digits =
2), "%", ")"), ylab = paste("PC2", "(",
round(100*x$importance[2,2], digits = 2), "%", ")") )
plot(delta.onto.space$x[,1], delta.onto.space$x[,3], cex = 2,
col = "transparent", pch = 21, bg = colours.1.3, frame.plot = T,
xlab = paste("PC1", "(", round(100*x$importance[2,1], digits =
2), "%", ")"), ylab = paste("PC3", "(",
round(100*x$importance[2,3], digits = 2), "%", ")") )
ggplot(data=delta.onto.space$x, aes(x=delta.onto.space$x[,1], y=
delta.onto.space$x[,3], legend = F))+
  geom_point(size = 5, stroke= 0.1,color =colours.1.3, alpha =
0.6)+scale_y_continuous(trans = "reverse")+
  theme(text = element_text(family = "Helvetica"))
+theme(axis.title.x = element_blank(),axis.text.x =
element_blank(),axis.text.y = element_blank(),axis.title.y =
element_blank(),axis.line = element_line(colour =
"black"),panel.grid.major = element_blank(),panel.background =
element_blank(),panel.grid.minor = element_blank())

text(delta.onto.space$x[,1], delta.onto.space$x[,3], labels =
rownames(delta.onto.traits), cex = 0.5)


######### ANCESTRAL STATE RECONSTRUCTION #######
######## OF HATCHLING SHAPE AND SIZE FOR #######
######## ASTERIORNIS AND PRESBYORNIS ##########

## Needs to read landmarks for expanded dataset and 'eroded'
landmarking scheme ##


# subset the adults or hatchlings from the shape data - i.e.,
expanded dataset eroded landmarking scheme

# age variable for expanded dataset

# ten items per row so we can keep track easily
```

```
age <- age_3


coords.temp <- gpa_3$coords[,, age_3 == "hatchling"] # subsets
coordinates
coords.temp.adult <- gpa_3$coords[,, age_3 == "adult"] # subsets
coordinates

cs.temp <- gpa_3$Csize[age_3 == "hatchling"] # subsets csize
cs.temp.adult <- gpa_3$Csize[age_3 == "adult"] # subsets csize


# change rownames to taxa so they match phylogeny

names <- as.character(dimnames(coords.temp)[[3]])
names.adults <- as.character(dimnames(coords.temp.adult)[[3]])

taxa <- strsplit(names, ".hatchling")
taxa.adults <- strsplit(names.adults, ".adult")

dimnames(coords.temp)[[3]] <- taxa
names(cs.temp) <- taxa

dimnames(coords.temp.adult)[[3]] <- taxa.adults
names(cs.temp.adult) <- taxa.adults


# next step is getting rid of taxa we don't want in the analyses

# first, get rid of non-hatchling malacorhynchus. -----fixed

plot3d(coords.temp[,,12], type = 's', size = 1 ,decorate =
FALSE)
aspect3d("iso")

coords.final <- coords.temp [,, c(1:10, 12:dim(coords.temp)
[[3]])]
cs.final <- cs.temp[c(1:10, 12:length(cs.temp))]


# get rid of Porphyrio too

outgroups <- c("Porphyrio","Presbyornis","Presbyornis
2","Asteriornis","Asteriornis Recon")

coords.less.inclusive <- coords.temp [,, !dimnames(coords.temp)
[[3]]  %in% outgroups ]
```

```
csize.less.inclusive <- cs.temp[!names(cs.temp) %in% outgroups]

coords.adults.less.inclusive <- coords.temp.adult [,, !
dimnames(coords.temp.adult) [[3]]  %in% outgroups ]
csize.adults.less.inclusive <- cs.temp.adult[!
names(cs.temp.adult) %in% outgroups]
# read new phylogeny that includes Anseranas, and the extant
outgroups

# Reads phylogeny for expanded dataset


tree.temp <- read.tree("new-larger.tre")

tree.temp$tip.label <- gsub("'", "", tree.temp$tip.label)

# check tree

plot( tree.temp, direction = "u", cex=0.5, label.offset = 0.5,
edge.width = 3,
      no.margin=F, y.lim = c(-1,90), x.lim = c(-1, 40),
align.tip.label = T, srt = -45)

axisPhylo(side = 2)
nodelabels() # to inspect the node numbers in the tree that we
want to use later on.


# check taxa in data and tree match

drop.taxa <- tree.temp$tip.label[ ! tree.temp$tip.label %in%
dimnames(coords.less.inclusive)[[3]] ]
drop.taxa # check the taxa are the expected to be drawn out

tree.final<- drop.tip(tree.temp, drop.taxa)

#check tree again

plot( tree.final, direction = "u", cex=0.5, label.offset = 0.5,
edge.width = 3,
      no.margin=F, y.lim = c(-1,90), x.lim = c(-1, 40),
align.tip.label = T, srt = -45)

axisPhylo(side = 2)
nodelabels() # to inspect the node numbers in the tree that we
want to use later on.
```

```
# ANCESTRAL SHAPES RECONSTRUCTION #

pca.with.ancestral.shapes <- gm.prcomp(coords.less.inclusive,
tree.final, GLS = T, transform = T)
pca.with.ancestral.shapes.adults <-
gm.prcomp(coords.adults.less.inclusive, tree.final, GLS = T,
transform = T)

ancestral.hatchlings <-
arrayspecs(pca.with.ancestral.shapes$ancestors, 100, 3)
ancestral.adults <-
arrayspecs(pca.with.ancestral.shapes.adults$ancestors, 100, 3)

# we are interested in the following nodes:

# 36: Neornithes
# 37: Neognathae
# 39: Galloanserae
# 40: Galliformes
# 53: Anseriformes
# 54: Anseriformes minus Chauna


hatchling.neornithes <- ancestral.hatchlings[,,"36"]
hatchling.neognathae <- ancestral.hatchlings[,,"37"]
hatchling.galloanserae<- ancestral.hatchlings[,, "39"]
hatchling.anseriformes <- ancestral.hatchlings [,, "53"]
hatchling.anseriformes.nochauna <- ancestral.hatchlings [,,
"54"]
hatchling.galliformes <- ancestral.hatchlings [,, "40"]

hatchling.palaeognathae <- ancestral.hatchlings[,, "69"]

hatchling.anatidae <- ancestral.hatchlings [,, "55"]

adult.neornithes <- ancestral.adults[,,"36"]
adult.neognathae <- ancestral.adults[,,"37"]
adult.galloanserae<- ancestral.adults[,, "39"]
adult.anseriformes <- ancestral.adults [,, "53"]
adult.anseriformes.nochauna <- ancestral.adults [,, "54"]
adult.galliformes <- ancestral.adults [,, "40"]

adult.palaeognathae <- ancestral.adults[,, "69"]

adult.anatidae <- ancestral.adults [,, "55"]
```

```
# check ancestral shapes


plot3d(adult.anatidae, type = 's', size = 1 ,decorate = FALSE)
aspect3d("iso")

plot3d(hatchling.galliformes, type = 's', size = 1 ,decorate =
FALSE)
aspect3d("iso")

open3d()
par3d(windowRect = c(0,0,700,250))
Sys.sleep(1)
mfrow3d(nr = 1, nc = 2, byrow = TRUE, sharedMouse = TRUE)
plot3d(hatchling.anseriformes, type = 's', size = 1 ,decorate =
FALSE)
aspect3d("iso")

next3d()
plot3d(hatchling.anatidae, type = 's', size = 1 ,decorate =
FALSE)
aspect3d("iso")

plotRefToTarget(adult.anatidae,hatchling.anatidae,method =
"points")####check adult and hatchling shapes



# NEXT STEPS: PRODUCING OTHER ESTIMATIONS USING OTHER MODELS?
EXPLORE AT LEAST S

gdf.final <- list(shape=two.d.array(coords.less.inclusive),
centroid.size=csize.less.inclusive)
gdf.final.adults <-
list(shape=two.d.array(coords.adults.less.inclusive),
centroid.size=csize.adults.less.inclusive)


#fit for models to hatchling shape evolution

fit1 <- mvgls(shape ~ 1, data = gdf.final,  tree = tree.final,
model="BM", penalty="RidgeArch", method="H&L")
fit2 <- mvgls(shape ~ 1, data = gdf.final,  tree =
tree.final,model="OU", penalty="RidgeArch", method="H&L")
fit3 <- mvgls(shape ~ 1, data = gdf.final,  tree = tree.final,
model="EB", penalty="RidgeArch", method="H&L")
fit4 <- mvgls(shape ~ 1, data = gdf.final,  tree = tree.final,
```

```
model="lambda", penalty="RidgeArch", method="H&L")

fit1.adults <- mvgls(shape ~ 1, data = gdf.final.adults,  tree =
tree.final, model="BM", penalty="RidgeArch", method="H&L")
fit2.adults <- mvgls(shape ~ 1, data = gdf.final.adults,  tree =
tree.final,model="OU", penalty="RidgeArch", method="H&L")
fit3.adults <- mvgls(shape ~ 1, data = gdf.final.adults,  tree =
tree.final, model="EB", penalty="RidgeArch", method="H&L")
fit4.adults <- mvgls(shape ~ 1, data = gdf.final.adults,  tree =
tree.final, model="lambda", penalty="RidgeArch", method="H&L")

GIC(fit1); GIC(fit2); GIC(fit3); GIC(fit4)  # compares fit with
a penalised likelihood based information criterion
GIC(fit1.adults); GIC(fit2.adults); GIC(fit3.adults);
GIC(fit4.adults)
#lambda is the preferred model

#we do the ancestral shape reconstruction with the preferred
model: lambda

ancestors.mvmorph <- ancestral(fit4)
ancestors.mvmorph.adults <- ancestral(fit4.adults)

ancestors.mvmorph <- arrayspecs(ancestors.mvmorph, 100, 3)
ancestors.mvmorph.adults <- arrayspecs(ancestors.mvmorph.adults,
100, 3)



# this block is to check the differences in ancestral node
shapes between the four models, mostly for internal control

dist.OU <- dist(two.d.array(ancestors.mvmorph))

dist.EB <- dist(two.d.array(ancestors.mvmorph))

dist.BM <- dist(two.d.array(ancestral.hatchlings))

dist.lambda<- dist(two.d.array(ancestors.mvmorph))

dist.lambda.adults<- dist(two.d.array(ancestors.mvmorph.adults))


# plot these differences

plot(dist.OU, cex = 3, pch = 21, bg = "#8A6A91", col =
"transparent")
```

```
points(dist.BM, cex = 3, pch = 21, bg = "#DBB76E", col =
"transparent")

points(dist.EB, cex = 1, pch = 21, bg = "#CE5782", col =
"transparent")

points(dist.lambda, cex = 3, pch = 21, bg = "#9EDB9E", col =
"transparent")

points(dist.lambda.adults, cex = 3, pch = 21, bg = "#9EDB9E",
col = "transparent")



#compare shapes between OU, EB and lambda models(mvMORPH) and BM
model (geomorph), equivalent computationally to mvmorph's BM.

open3d()
par3d(windowRect = c(0,0,700,250))
Sys.sleep(1)
mfrow3d(nr = 1, nc = 2, byrow = TRUE, sharedMouse = TRUE)
plot3d(hatchling.galloanserae, type = 's', size = 1 ,decorate =
FALSE)
aspect3d("iso")

next3d()
plot3d(ancestors.mvmorph.final[,,"node_39"], type = 's', size =
1 ,decorate = FALSE)
aspect3d("iso")

####################################
#########Estimate fossil hatchling shapes
####################################

onto.galloanserae<- ancestors.mvmorph.adults[,,4]-
ancestors.mvmorph[,,4]
onto.galliformes<- ancestors.mvmorph.adults[,,5]-
ancestors.mvmorph[,,5]
onto.anseriformes<- ancestors.mvmorph.adults[,,18]-
ancestors.mvmorph[,,18]
onto.anatidae <- ancestors.mvmorph.adults[,,19]-
ancestors.mvmorph[,,19]

#####Asteriornis hatchling

###using galloanserae ontogeny
```

```
Aster.hatch.shape.1 <- gpa_3$coords[,,69]-onto.galloanserae


plot3d(gpa_3$coords[,,69], type = 's', size = 1.5 ,col =
"grey",decorate = FALSE)
aspect3d("iso")

rgl.snapshot( "Asteriornis adult.png" )

plot3d(Aster.hatch.shape.1, type = 's', size = 1.5 ,col =
"grey",decorate = FALSE)
aspect3d("iso")

rgl.snapshot( "Asteriornis hatchling-1.png" )

Aster.adult.mesh <- read.ply("/Users/basselarnaout/Desktop/PhD/
Posthatching series/Landmarks/Asteriornis Recon/Skull
restored.ply", ShowSpecimen = FALSE, addNormals = TRUE)
Aster.hatch.1 <- tps3d(Aster.adult.mesh, refmat =
AllAsteriornis_Reconlandmarks, tarmat = Aster.hatch.shape.1)

plotRefToTarget(Aster.hatch.shape.1,gpa_3$coords[,,69], method =
"points")

shade3d(Aster.hatch.1, col= 8, alpha = 1)

###using galliform ontogeny
Aster.hatch.shape.2 <- gpa_3$coords[,,69]-onto.galliformes

plot3d(Aster.hatch.shape.2, type = 's', size = 1.5 ,col =
"grey",decorate = FALSE)
aspect3d("iso")

rgl.snapshot( "Asteriornis hatchling-2.png" )

Aster.hatch.2 <- tps3d(Aster.adult.mesh, refmat =
AllAsteriornis_Reconlandmarks, tarmat = Aster.hatch.shape.2)

plotRefToTarget(Aster.hatch.shape.2,gpa_3$coords[,,69], method =
"points")

shade3d(Aster.hatch.2, col= 8, alpha = 1)
#####presbyornis hatchling

###using galloanserae ontogeny
Presby.hatch.shape.1 <- gpa_3$coords[,,67]-onto.galloanserae
```

```
plot3d(gpa_3$coords[,,67], type = 's', size = 1.5 ,col =
"grey",decorate = FALSE)
aspect3d("iso")

rgl.snapshot( "Presbyornis adult.png" )

plot3d(Presby.hatch.shape.1, type = 's', size = 1.5 ,col =
"grey",decorate = FALSE)
aspect3d("iso")

rgl.snapshot( "Presbyornis hatchling-1.png" )

Presby.adult.mesh <- read.ply("/Users/basselarnaout/Desktop/PhD/
Posthatching series/Landmarks/Presbyornis 2/Model.ply",
ShowSpecimen = FALSE, addNormals = TRUE)
Presby.hatch.1 <- tps3d(Presby.adult.mesh, refmat =
AllPresby2landmarks, tarmat = Presby.hatch.shape.1)

plotRefToTarget(Presby.hatch.shape.1,gpa$coords[,,67], method =
"points")

shade3d(Presby.hatch.1, col= 8, alpha = 1)

###using anseriform ontogeny
Presby.hatch.shape.2 <- gpa_3$coords[,,67]-onto.anseriformes

plot3d(Presby.hatch.shape.2, type = 's', size = 1.5 ,col =
"grey",decorate = FALSE)
aspect3d("iso")

rgl.snapshot( "Presbyornis hatchling-2.png" )

Presby.hatch.2 <- tps3d(Presby.adult.mesh, refmat =
AllPresby2landmarks, tarmat = Presby.hatch.shape.2)

plotRefToTarget(Presby.hatch.shape.2,gpa$coords[,,67], method =
"points")

shade3d(Presby.hatch.2, col= 8, alpha = 1)

###using anatidae ontogeny
Presby.hatch.shape.3 <- gpa_3$coords[,,67]-onto.anatidae

plot3d(Presby.hatch.shape.3, type = 's', size = 1.5 ,col =
"grey",decorate = FALSE)
aspect3d("iso")
```

```
rgl.snapshot( "Presbyornis hatchling-3.png" )

Presby.hatch.3 <- tps3d(Presby.adult.mesh, refmat =
AllPresby2landmarks, tarmat = Presby.hatch.shape.3)

plotRefToTarget(Presby.hatch.shape.3,gpa$coords[,,67], method =
"points")

shade3d(Presby.hatch.3, col= 8, alpha = 1)

fossil.hatchlings <-
array(c(Aster.hatch.shape.1,Aster.hatch.shape.2,Presby.hatch.sha
pe.1,Presby.hatch.shape.2,Presby.hatch.shape.3), c(100,3,5))
##########################

# NEXT STEPS: PROJECT SHAPE INTO PCA USING THE ANCESTRAL
HATCHLING SHAPES FROM THE BEST MODEL

# make an object for the final ancestral shapes with the
preferred model


ancestors.mvmorph.final <- ancestral(fit4)
ancestors.mvmorph.final <- arrayspecs(ancestors.mvmorph.final,
100, 3)

ancestors.mvmorph.final.adults <- ancestral(fit4)
ancestors.mvmorph.final.adults <-
arrayspecs(ancestors.mvmorph.final.adults, 100, 3)

plot3d(ancestors.mvmorph.final[,,19], type = 's', size =
1.5 ,col = "grey",decorate = FALSE)
aspect3d("iso")

rgl.snapshot( "Anatidae hatchling.png" )

# First, we need to make the PCA with the full eroded dataset

# read the full 'eroded' dataset

eroded.coords <- gpa_3$coords

# take out porphyrio

eroded.coords <- eroded.coords [,, -c(72,73)]
dimnames(eroded.coords) # check label names
```

```
PCA.all<- gm.prcomp(eroded.coords)
x<-summary(PCA.all)


plot3d(eroded.coords[,,"Afropavo.hatchling"], type = 's', size =
1 ,decorate = FALSE)
aspect3d("iso")


#lets try to calculate now the pcscores for an ancestral shape

ancestors.final <- two.d.array(ancestors.mvmorph.final)
ancestors.final.adults <- two.d.array(ancestors.mvmorph.adults)
fossil.hatchling.shapes <- two.d.array(fossil.hatchlings)

ancestral.pcscores <- matrix(data= NA, nrow =
dim(ancestors.final)[1], ncol = 3)
row.names(ancestral.pcscores) <- row.names(ancestors.final)

ancestral.pcscores.adults <- matrix(data= NA, nrow =
dim(ancestors.final.adults)[1], ncol = 3)
row.names(ancestral.pcscores.adults) <-
row.names(ancestors.final.adults)

fossil.hatchling.pcscores <- matrix(data= NA, nrow =
dim(fossil.hatchling.shapes)[1], ncol = 3)
#row.names(fossil.hatchling.pcscores) <-
row.names(fossil.hatchling.shapes)

for( i in 1:dim(ancestors.final)[1] ) {


  ancestral.hatchling.shape <- ancestors.final[i,]

  PC1.score <- sum(ancestral.hatchling.shape * PCA.all$rotation
[,1])

  ancestral.pcscores[i,1] <- PC1.score

  PC2.score <- sum(ancestral.hatchling.shape * PCA.all$rotation
[,2])

  ancestral.pcscores[i,2]<-PC2.score

  PC3.score <- sum(ancestral.hatchling.shape * PCA.all$rotation
[,3])
```

```
  ancestral.pcscores[i,3]<-PC3.score



}

ancestral.pcscores

for( i in 1:dim(ancestors.final.adults)[1] ) {


  ancestral.adult.shape <- ancestors.final.adults[i,]

  PC1.score <- sum(ancestral.adult.shape * PCA.all$rotation
[,1])

  ancestral.pcscores.adults[i,1] <- PC1.score

  PC2.score <- sum(ancestral.adult.shape * PCA.all$rotation [,
2])

  ancestral.pcscores.adults[i,2]<-PC2.score

  PC3.score <- sum(ancestral.adult.shape * PCA.all$rotation [,
3])

  ancestral.pcscores.adults[i,3]<-PC3.score




}

ancestral.pcscores.adults



for( i in 1:dim(fossil.hatchling.shapes)[1] ) {


  fossil.hatchlings.shape.2 <- fossil.hatchling.shapes[i,]

  PC1.score <- sum(fossil.hatchlings.shape.2 * PCA.all$rotation
[,1])
```

```
  fossil.hatchling.pcscores[i,1] <- PC1.score

  PC2.score <- sum(fossil.hatchlings.shape.2 * PCA.all$rotation
[,2])

  fossil.hatchling.pcscores[i,2]<-PC2.score

  PC3.score <- sum(fossil.hatchlings.shape.2 * PCA.all$rotation
[,3])

  fossil.hatchling.pcscores[i,3]<-PC3.score




}
fossil.hatchling.pcscores


#final plot


plot(PCA.all$x[,1], PCA.all$x[,2], pch = 16,col = c("#5E96B4",
"#5E96B4", "#54B473", "#54B473", "#5E96B4",
                                                    "#5E96B4",
"#54B473", "#54B473", "#54B473", "#54B473",
                                                    "#5E96B4",
"#5E96B4", "#54B473", "#54B473", "#5E96B4",
                                                    "#5E96B4",
"#54B473", "#54B473","#5E96B4", "#5E96B4",

                                                    "#5E96B4",
"#5E96B4", "#5E96B4", "#5E96B4", "#5E96B4",
                                                    "#5E96B4",
"#54B473", "#54B473", "#5E96B4",
                                                    "#5E96B4",
"#54B473", "#54B473","#5E96B4","#5E96B4",
                                                    "#54B473",
"#54B473", "#54B473", "#54B473", "#54B473",

                                                    "#54B473",
"#5E96B4","#5E96B4","#5E96B4","#5E96B4",
                                                    "#54B473",
"#54B473","#5E96B4","#5E96B4", "#54B473",
                                                    "#54B473",
"#54B473", "#54B473","#5E96B4","#5E96B4",
```

```
                                                              "#54B473",
"#54B473","#5E96B4","#5E96B4","#5E96B4",

                                                              "#5E96B4",
"#E5CA7F", "#E5CA7F","#D84C71", "#D84C71",
                                                              "#855BC1",
"#855BC1", "dark blue", "dark blue", "black",
                                                              "#5E96B4",
"#5E96B4","pink","pink"), cex = 2,
     xlab = paste("PC1", "(", round(100*x$PC.summary[2,1],
digits = 2), "%", ")"),
     ylab = paste("PC2", "(", round(100*x$PC.summary[2,2],
digits = 2), "%", ")") )

text(PCA.all$x[,1], PCA.all$x[,2], cex= 0.5, labels =
names(PCA.all$x[,2]), pos = 3)


points(ancestral.pcscores[c(4,5,18,19),1],
ancestral.pcscores[c(4,5,18,19),3])
points(ancestral.pcscores.adults[c(4,5,18,19),1],ancestral.pcsco
res.adults[c(4,5,18,19),3])
text(ancestral.pcscores[c(4,5,18,19),1],
ancestral.pcscores[c(4,5,18,19),2], labels =
row.names(ancestral.pcscores)[c(4,5,18,19)], cex = 0.5)
text(ancestral.pcscores.adults[c(4,5,18,19),1],
ancestral.pcscores.adults[c(4,5,18,19),2], labels =
row.names(ancestral.pcscores.adults)[c(4,5,18,19)], cex = 0.5)
points(fossil.hatchling.pcscores[,1],fossil.hatchling.pcscores[,
3],pch=2)
text(fossil.hatchling.pcscores[,1],
fossil.hatchling.pcscores[,3], labels = c("1","2","3","4","5"),
cex = 0.5)

###ggplot
specimen.colors.3 <- c("#5E96B4", "#5E96B4", "#54B473",
"#54B473", "#5E96B4",
                       "#5E96B4", "#54B473", "#54B473",
"#54B473", "#54B473",
                       "#5E96B4", "#5E96B4", "#54B473",
"#54B473", "#5E96B4",
                       "#5E96B4", "#54B473",
"#54B473","#5E96B4", "#5E96B4",

                       "#5E96B4", "#5E96B4", "#5E96B4",
"#5E96B4", "#5E96B4",
                       "#5E96B4", "#54B473", "#54B473",
```

```
"#5E96B4",
                       "#5E96B4", "#54B473",
"#54B473","#5E96B4","#5E96B4",
                       "#54B473", "#54B473", "#54B473",
"#54B473", "#54B473",

                       "#54B473",
"#5E96B4","#5E96B4","#5E96B4","#5E96B4",
                       "#54B473", "#54B473","#5E96B4","#5E96B4",
"#54B473",
                       "#54B473", "#54B473",
"#54B473","#5E96B4","#5E96B4",
                       "#54B473",
"#54B473","#5E96B4","#5E96B4","#5E96B4",

                       "#5E96B4", "#E5CA7F",
"#E5CA7F","#D84C71", "#D84C71",
                       "#855BC1", "#855BC1", "dark blue", "dark
blue", "black",
                       "#5E96B4", "#5E96B4","pink","pink")
shapes.3 <- c(23, 21, 23, 21, 23, 21, 21, 23, 21,23,
              21, 23, 21,23, 21, 23, 21, 23, 21,
              23, 21, 23, 21, 23, 23, 21, 21, 23,
              21,23,23, 21, 21,23, 21,23, 21,23,
              21,23, 21,23, 21,23, 21,23, 21,23,
              21,23, 21,23, 21,23, 21,23, 21,23,
              21,23,21,23,21,23,21,23,23,21,23,23,21,21,23)

ggplot(data=PCA.all$x[,1:2], aes(x=PCA.all$x[,1], y=
PCA.all$x[,2], color = specimen.colors.3, legend = F))
+theme(text = element_text(family = "Helvetica"))+
  geom_point(shape = shapes.3,size = 4, stroke= 0.2,color =
"black",fill =specimen.colors.3, alpha = 10)+ theme(axis.title.x
= element_blank(),axis.text.x = element_blank(),axis.text.y =
element_blank(),axis.title.y = element_blank(),axis.line =
element_line(colour = "black"),

panel.grid.major = element_blank(),panel.background =
element_blank(),panel.grid.minor = element_blank())+
  geom_segment(aes(x = PCA.all$x[2], y = PCA.all$x[2,2], xend =
PCA.all$x[1], yend = PCA.all$x[1,2]), linewidth = 1,color
='#4E99B7',alpha = 0.01)+
  geom_segment(aes(x = PCA.all$x[4], y = PCA.all$x[4,2], xend =
PCA.all$x[3], yend = PCA.all$x[3,2]), linewidth = 1,color
='#5EB16B',alpha = 0.01)+
  geom_segment(aes(x = PCA.all$x[6], y = PCA.all$x[6,2], xend =
PCA.all$x[5], yend = PCA.all$x[5,2]), linewidth = 1,color
```

```
='#4E99B7',alpha = 0.01)+
  geom_segment(aes(x = PCA.all$x[7], y = PCA.all$x[7,2], xend =
PCA.all$x[8], yend = PCA.all$x[8,2]), linewidth = 1,color
='#5EB16B',alpha = 0.01)+
  geom_segment(aes(x = PCA.all$x[9], y = PCA.all$x[9,2], xend =
PCA.all$x[10], yend = PCA.all$x[10,2]), linewidth = 1,color
='#5EB16B',alpha = 0.01)+
  geom_segment(aes(x = PCA.all$x[11], y = PCA.all$x[11,2], xend
= PCA.all$x[12], yend = PCA.all$x[12,2]), linewidth = 1,color
='#4E99B7',alpha = 0.01)+
  geom_segment(aes(x = PCA.all$x[13], y = PCA.all$x[13,2], xend
= PCA.all$x[14], yend = PCA.all$x[14,2]), linewidth = 1,color
='#5EB16B',alpha = 0.01)+
  geom_segment(aes(x = PCA.all$x[15], y = PCA.all$x[15,2], xend
= PCA.all$x[16], yend = PCA.all$x[16,2]), linewidth = 1,color
='#4E99B7',alpha = 0.01)+
  geom_segment(aes(x = PCA.all$x[17], y = PCA.all$x[17,2], xend
= PCA.all$x[18], yend = PCA.all$x[18,2]), linewidth = 1,color
='#5EB16B',alpha = 0.01)+
  geom_segment(aes(x = PCA.all$x[19], y = PCA.all$x[19,2], xend
= PCA.all$x[20], yend = PCA.all$x[20,2]), linewidth = 1,color
='#4E99B7',alpha = 0.01)+
  geom_segment(aes(x = PCA.all$x[21], y = PCA.all$x[21,2], xend
= PCA.all$x[22], yend = PCA.all$x[22,2]), linewidth = 1,color
='#4E99B7',alpha = 0.01)+
  geom_segment(aes(x = PCA.all$x[23], y = PCA.all$x[23,2], xend
= PCA.all$x[24], yend = PCA.all$x[24,2]), linewidth = 1,color
='#4E99B7',alpha = 0.01)+
  geom_segment(aes(x = PCA.all$x[25], y = PCA.all$x[25,2], xend
= PCA.all$x[26], yend = PCA.all$x[26,2]), linewidth = 1,color
='#4E99B7',alpha = 0.01)+
  geom_segment(aes(x = PCA.all$x[27], y = PCA.all$x[27,2], xend
= PCA.all$x[28], yend = PCA.all$x[28,2]), linewidth = 1,color
='#5EB16B',alpha = 0.01)+
  geom_segment(aes(x = PCA.all$x[29], y = PCA.all$x[29,2], xend
= PCA.all$x[30], yend = PCA.all$x[30,2]), linewidth = 1,color
='#4E99B7',alpha = 0.01)+
  geom_segment(aes(x = PCA.all$x[31], y = PCA.all$x[31,2], xend
= PCA.all$x[32], yend = PCA.all$x[32,2]), linewidth = 1,color
='#5EB16B',alpha = 0.01)+
  geom_segment(aes(x = PCA.all$x[33], y = PCA.all$x[33,2], xend
= PCA.all$x[34], yend = PCA.all$x[34,2]), linewidth = 1,color
='#4E99B7',alpha = 0.01)+
  geom_segment(aes(x = PCA.all$x[35], y = PCA.all$x[35,2], xend
= PCA.all$x[36], yend = PCA.all$x[36,2]), linewidth = 1,color
='#5EB16B',alpha = 0.01)+
  geom_segment(aes(x = PCA.all$x[37], y = PCA.all$x[37,2], xend
```

```
= PCA.all$x[38], yend = PCA.all$x[38,2]), linewidth = 1,color
='#5EB16B',alpha = 0.01)+
  geom_segment(aes(x = PCA.all$x[39], y = PCA.all$x[39,2], xend
= PCA.all$x[40], yend = PCA.all$x[40,2]), linewidth = 1,color
='#5EB16B',alpha = 0.01)+
  geom_segment(aes(x = PCA.all$x[41], y = PCA.all$x[41,2], xend
= PCA.all$x[42], yend = PCA.all$x[42,2]), linewidth = 1,color
='#4E99B7',alpha = 0.01)+
  geom_segment(aes(x = PCA.all$x[43], y = PCA.all$x[43,2], xend
= PCA.all$x[44], yend = PCA.all$x[44,2]), linewidth = 1,color
='#4E99B7',alpha = 0.01)+
  geom_segment(aes(x = PCA.all$x[45], y = PCA.all$x[45,2], xend
= PCA.all$x[46], yend = PCA.all$x[46,2]), linewidth = 1,color
='#5EB16B',alpha = 0.01)+
  geom_segment(aes(x = PCA.all$x[47], y = PCA.all$x[47,2], xend
= PCA.all$x[48], yend = PCA.all$x[48,2]), linewidth = 1,color
='#4E99B7',alpha = 0.01)+
  geom_segment(aes(x = PCA.all$x[49], y = PCA.all$x[49,2], xend
= PCA.all$x[50], yend = PCA.all$x[50,2]), linewidth = 1,color
='#5EB16B',alpha = 0.01)+
  geom_segment(aes(x = PCA.all$x[51], y = PCA.all$x[51,2], xend
= PCA.all$x[52], yend = PCA.all$x[52,2]), linewidth = 1,color
='#5EB16B',alpha = 0.01)+
  geom_segment(aes(x = PCA.all$x[53], y = PCA.all$x[53,2], xend
= PCA.all$x[54], yend = PCA.all$x[54,2]), linewidth = 1,color
='#4E99B7',alpha = 0.01)+
  geom_segment(aes(x = PCA.all$x[55], y = PCA.all$x[55,2], xend
= PCA.all$x[56], yend = PCA.all$x[56,2]), linewidth = 1,color
='#5EB16B',alpha = 0.01)+
  geom_segment(aes(x = PCA.all$x[57], y = PCA.all$x[57,2], xend
= PCA.all$x[58], yend = PCA.all$x[58,2]), linewidth = 1,color
='#4E99B7',alpha = 0.01)+
  geom_segment(aes(x = PCA.all$x[59], y = PCA.all$x[59,2], xend
= PCA.all$x[60], yend = PCA.all$x[60,2]), linewidth = 1,color
='#4E99B7',alpha = 0.01)+
  geom_segment(aes(x = PCA.all$x[61], y = PCA.all$x[61,2], xend
= PCA.all$x[62], yend = PCA.all$x[62,2]), linewidth = 1,color
='#E5CA7F',alpha = 0.01)+
  geom_segment(aes(x = PCA.all$x[63], y = PCA.all$x[63,2], xend
= PCA.all$x[64], yend = PCA.all$x[64,2]), linewidth = 1,color
='#D84C71',alpha = 0.01)+
  geom_segment(aes(x = PCA.all$x[65], y = PCA.all$x[65,2], xend
= PCA.all$x[66], yend = PCA.all$x[66,2]), linewidth = 1,color
='#855BC1',alpha = 0.01)+
  geom_segment(aes(x = PCA.all$x[67], y = PCA.all$x[67,2], xend
= PCA.all$x[68], yend = PCA.all$x[68,2]), linewidth = 1,color
='darkblue',alpha = 0.01)+
```

```
  geom_segment(aes(x = PCA.all$x[70], y = PCA.all$x[70,2], xend
= PCA.all$x[71], yend = PCA.all$x[71,2]), linewidth = 1,color
='#5E96B4',alpha = 0.01)+
  geom_segment(aes(x = PCA.all$x[72], y = PCA.all$x[72,2], xend
= PCA.all$x[73], yend = PCA.all$x[73,2]), linewidth = 1,color
='pink',alpha = 0.01)

#Extract extreme shapes from PCA

open3d()
par3d(windowRect = c(0,0,700,250))
Sys.sleep(1)
mfrow3d(nr = 1, nc = 2, byrow = TRUE, sharedMouse = TRUE)
plot3d(preds$pred1, type = 's', size = 1 ,decorate = FALSE)
aspect3d("iso")

next3d()
plot3d(preds$pred2, type = 's', size = 1 ,decorate = FALSE)
aspect3d("iso")


PC=3
PC.temp<-PCA.all$x[,PC]

preds<-shape.predictor(eroded.coords, x =
as.numeric( PCA.all$x[,PC] ), Intercept = F,
                       pred1 = quantile(PC.temp, probs = 0.5),
pred2 = quantile(PC.temp, probs = 0.97))


#take snapshot in each position
rgl.snapshot( "PC3 lateral.png" )


##########################
########## 3D plots ######
##########################

##PCA 3d plot
array.2<- data.frame(PCA.all$x[,1],PCA.all$x[,2],PCA.all$x[,3])

families_3<- families_3[-c(72,73)]
age_3<- age_3[-c(72,73)]
subset <- c(which(families_3 == "Galliformes"),which(families_3
== "Anseriformes"))
families.subset<- as.vector(families_3[subset])
age.subset <- as.vector(age_3[subset])
```

```
####with shadows 3d plot
plot_ly(array.2,x=
~PCA.all$x[subset,1],y=~PCA.all$x[subset,2],z=~PCA.all$x[subset,
3], marker = list(opacity =1,line=list(width=1,color='black')),
        type = 'scatter3d',mode = 'markers',color =
as.factor(interaction(families.subset,age.subset)),opacity =
0.5,
        colors=c("#4E99B7","#4EB773","#a6ccdb","#a6dbb9"))%>%
  layout(scene = list(xaxis = list(title = 'PC1 (48.9%)',
showbackground = TRUE, backgroundcolor = "rgba(232,232,232,1)"),
                      yaxis = list(title = 'PC2 (16.01%)',
showbackground = TRUE, backgroundcolor = "rgba(211,211,211,1)"),
                      zaxis = list(title = 'PC3 (8.82%)')))%>%
  add_trace(x = ~PCA.all$x[33:34,1], y = ~PCA.all$x[33:34,2], z
= ~PCA.all$x[33:34,3], inherit = F, mode = "lines", type =
"scatter3d", line = list( color = "#4E99B7"), opacity =
0.5,showlegend = F)%>%
  add_trace(x = ~PCA.all$x[61:62,1], y = ~PCA.all$x[61:62,2], z
= ~PCA.all$x[61:62,3], inherit = F, mode = "markers", type =
"scatter3d", marker =
list( line=list(width=1,color='black'),color = "#E5CA7F"),
opacity = 1,showlegend = F)%>%
  add_trace(x = ~PCA.all$x[61:62,1], y = ~PCA.all$x[61:62,2], z
= ~min(PCA.all$x[,3]-0.01), inherit = F, mode = "markers", type
= "scatter3d", marker = list( color = "#848688"), opacity =
0.3,showlegend = F)%>%
  add_trace(x = ~PCA.all$x[61:62,1], y = ~PCA.all$x[61:62,2], z
= ~PCA.all$x[61:62,3], inherit = F, mode = "lines", type =
"scatter3d", line = list( color = "#E5CA7F"), opacity =
1,showlegend = F)%>%
  add_trace(x = c(PCA.all$x[61,1],PCA.all$x[61,1]), y =
c(list(PCA.all$x[61,2]),list(PCA.all$x[61,2])), z =
c(list(PCA.all$x[61,3]),rep(list(-0.118561),2)), inherit = F,
mode = "lines", type = "scatter3d",line = list(color =
'#E5CA7F'),showlegend = F, opacity = 1)%>%
  add_trace(x = c(PCA.all$x[62,1],PCA.all$x[62,1]), y =
c(list(PCA.all$x[62,2]),list(PCA.all$x[62,2])), z =
c(list(PCA.all$x[62,3]),rep(list(-0.118561),2)), inherit = F,
mode = "lines", type = "scatter3d",line = list(color =
'#E5CA7F'),showlegend = F, opacity = 1)%>%
  add_trace(x = ~PCA.all$x[72:73,1], y = ~PCA.all$x[72:73,2], z
= ~PCA.all$x[72:73,3], inherit = F, mode = "markers", type =
"scatter3d", marker =
list( line=list(width=1,color='black'),color = "pink"), opacity
= 1,showlegend = F)%>%
  add_trace(x = c(PCA.all$x[72,1],PCA.all$x[72,1]), y =
```

```
c(list(PCA.all$x[72,2]),list(PCA.all$x[72,2])), z =
c(list(PCA.all$x[72,3]),rep(list(-0.118561),2)), inherit = F,
mode = "lines", type = "scatter3d",line = list(color =
'pink'),showlegend = F, opacity = 1)%>%
  add_trace(x = c(PCA.all$x[73,1],PCA.all$x[73,1]), y =
c(list(PCA.all$x[73,2]),list(PCA.all$x[73,2])), z =
c(list(PCA.all$x[73,3]),rep(list(-0.118561),2)), inherit = F,
mode = "lines", type = "scatter3d",line = list(color =
'pink'),showlegend = F, opacity = 1)%>%
  add_trace(x = ~PCA.all$x[72:73,1], y = ~PCA.all$x[72:73,2], z
= ~PCA.all$x[72:73,3], inherit = F, mode = "lines", type =
"scatter3d", line = list( color = "pink"), opacity =
1,showlegend = F)%>%
  add_trace(x = ~PCA.all$x[72:73,1], y = ~PCA.all$x[72:73,2], z
= ~min(PCA.all$x[,3]-0.01), inherit = F, mode = "markers", type
= "scatter3d", marker = list( color = "#848688"), opacity =
0.3,showlegend = F)%>%
  add_trace(x = ~PCA.all$x[63:64,1], y = ~PCA.all$x[63:64,2], z
= ~PCA.all$x[63:64,3], inherit = F, mode = "markers", type =
"scatter3d", marker =
list( line=list(width=1,color='black'),color = "#D84C71"),
opacity = 1,showlegend = F)%>%
  add_trace(x = ~PCA.all$x[63:64,1], y = ~PCA.all$x[63:64,2], z
= ~PCA.all$x[63:64,3], inherit = F, mode = "lines", type =
"scatter3d", line = list( color = "#D84C71"), opacity =
1,showlegend = F)%>%
  add_trace(x = ~PCA.all$x[63,1], y = ~PCA.all$x[63,2], z =
~min(PCA.all$x[,3]-0.01), inherit = F, mode = "markers", type =
"scatter3d", marker = list( color = "#848688"), opacity =
0.3,showlegend = F)%>%
  add_trace(x = ~PCA.all$x[64,1], y = ~PCA.all$x[64,2], z =
~min(PCA.all$x[,3]-0.01), inherit = F, mode = "markers", type =
"scatter3d", marker = list( color = "#848688"), opacity =
0.3,showlegend = F)%>%
  add_trace(x = c(PCA.all$x[63,1],PCA.all$x[63,1]), y =
c(list(PCA.all$x[63,2]),list(PCA.all$x[63,2])), z =
c(list(PCA.all$x[63,3]),rep(list(-0.118561),2)), inherit = F,
mode = "lines", type = "scatter3d",line = list(color =
'#D84C71'),showlegend = F, opacity = 0.5)%>%
  add_trace(x = c(PCA.all$x[64,1],PCA.all$x[64,1]), y =
c(list(PCA.all$x[64,2]),list(PCA.all$x[64,2])), z =
c(list(PCA.all$x[64,3]),rep(list(-0.118561),2)), inherit = F,
mode = "lines", type = "scatter3d",line = list(color =
'#D84C71'),showlegend = F, opacity = 0.5)%>%
  add_trace(x = ~PCA.all$x[65:66,1], y = ~PCA.all$x[65:66,2], z
= ~PCA.all$x[65:66,3], inherit = F, mode = "markers", type =
"scatter3d", marker =
```

```
list( line=list(width=1,color='black'),color = "#855BC1"),
opacity = 1,showlegend = F)%>%
  add_trace(x = ~PCA.all$x[65:66,1], y = ~PCA.all$x[65:66,2], z
= ~PCA.all$x[65:66,3], inherit = F, mode = "lines", type =
"scatter3d", line = list( color = "#855BC1"), opacity =
1,showlegend = F)%>%
  add_trace(x = ~PCA.all$x[65,1], y = ~PCA.all$x[65,2], z =
~min(PCA.all$x[,3]-0.01), inherit = F, mode = "markers", type =
"scatter3d", marker = list( color = "#848688"), opacity =
0.3,showlegend = F)%>%
  add_trace(x = ~PCA.all$x[66,1], y = ~PCA.all$x[66,2], z =
~min(PCA.all$x[,3]-0.01), inherit = F, mode = "markers", type =
"scatter3d", marker = list( color = "#848688"), opacity =
0.3,showlegend = F)%>%
  add_trace(x = c(PCA.all$x[65,1],PCA.all$x[65,1]), y =
c(list(PCA.all$x[65,2]),list(PCA.all$x[65,2])), z =
c(list(PCA.all$x[65,3]),rep(list(-0.118561),2)), inherit = F,
mode = "lines", type = "scatter3d",line = list(color =
'#855BC1'),showlegend = F, opacity = 0.5)%>%
  add_trace(x = c(PCA.all$x[66,1],PCA.all$x[66,1]), y =
c(list(PCA.all$x[66,2]),list(PCA.all$x[66,2])), z =
c(list(PCA.all$x[66,3]),rep(list(-0.118561),2)), inherit = F,
mode = "lines", type = "scatter3d",line = list(color =
'#855BC1'),showlegend = F, opacity = 0.5)%>%
  add_trace(x = ~PCA.all$x[69,1], y = ~PCA.all$x[69,2], z =
~PCA.all$x[69,3], inherit = F, mode = "markers", type =
"scatter3d", marker =
list( line=list(width=1,color='black'),color = "black"), opacity
= 1,showlegend = F)%>%
  add_trace(x = ~PCA.all$x[69,1], y = ~PCA.all$x[69,2], z =
~min(PCA.all$x[,3]-0.01), inherit = F, mode = "markers", type =
"scatter3d", marker = list( line=list(color='#848688'),color =
"#848688"), opacity = 1,showlegend = F)%>%
  add_trace(x = c(PCA.all$x[69,1],PCA.all$x[69,1]), y =
c(list(PCA.all$x[69,2]),list(PCA.all$x[69,2])), z =
c(list(PCA.all$x[69,3]),rep(list(-0.118561),2)), inherit = F,
mode = "lines", type = "scatter3d",line = list(color =
'black'),showlegend = F, opacity = 0.5)%>%
  add_trace(x =
~c(PCA.all$x[69,1],fossil.hatchling.pcscores[1,1]), y =
~c(PCA.all$x[69,2],fossil.hatchling.pcscores[1,2]), z =
~c(PCA.all$x[69,3],fossil.hatchling.pcscores[1,3]), inherit = F,
mode = "markers", type = "scatter3d", marker =
list( line=list(width=1,color='black'),color = "black"), opacity
= 1,showlegend = F)%>%
  add_trace(x = ~fossil.hatchling.pcscores[1,1], y =
~fossil.hatchling.pcscores[1,2], z = ~min(PCA.all$x[,3]-0.01),
```

```
inherit = F, mode = "markers", type = "scatter3d", marker =
list( line=list(color='#848688'),color = "#848688"), opacity =
0.3,showlegend = F)%>% ###projection point
  add_trace(x =
c(fossil.hatchling.pcscores[1,1],fossil.hatchling.pcscores[1,1])
, y =
c(list(fossil.hatchling.pcscores[1,2]),list(fossil.hatchling.pcs
cores[1,2])), z =
c(list(fossil.hatchling.pcscores[1,3]),rep(list(-0.118561),2)),
inherit = F, mode = "lines", type = "scatter3d",line =
list(color = 'black'),showlegend = F, opacity = 0.5)%>%
###projection line
  add_trace(x = ~fossil.hatchling.pcscores[2,1], y =
~fossil.hatchling.pcscores[2,2], z = ~min(PCA.all$x[,3]-0.01),
inherit = F, mode = "markers", type = "scatter3d", marker =
list( line=list(color='#848688'),color = "#848688"), opacity =
0.3,showlegend = F)%>%
  add_trace(x =
c(fossil.hatchling.pcscores[2,1],fossil.hatchling.pcscores[2,1])
, y =
c(list(fossil.hatchling.pcscores[2,2]),list(fossil.hatchling.pcs
cores[2,2])), z =
c(list(fossil.hatchling.pcscores[2,3]),rep(list(-0.118561),2)),
inherit = F, mode = "lines", type = "scatter3d",line =
list(color = 'black'),showlegend = F, opacity = 0.5)%>%
###projection line
  add_trace(x =
~c(PCA.all$x[69,1],fossil.hatchling.pcscores[1,1]), y =
~c(PCA.all$x[69,2],fossil.hatchling.pcscores[1,2]), z =
~c(PCA.all$x[69,3],fossil.hatchling.pcscores[1,3]), inherit = F,
mode = "lines", type = "scatter3d", line =
list( line=list(width=1,color='black'),color = "black",dash =
'longdashdot'), opacity = 1,showlegend = F)%>%
  add_trace(x =
~c(PCA.all$x[69,1],fossil.hatchling.pcscores[2,1]), y =
~c(PCA.all$x[69,2],fossil.hatchling.pcscores[2,2]), z =
~c(PCA.all$x[69,3],fossil.hatchling.pcscores[2,3]), inherit = F,
mode = "markers", type = "scatter3d", marker =
list( line=list(width=1,color='black'),color = "black"), opacity
= 1,showlegend = F)%>%
  add_trace(x =
~c(PCA.all$x[69,1],fossil.hatchling.pcscores[2,1]), y =
~c(PCA.all$x[69,2],fossil.hatchling.pcscores[2,2]), z =
~c(PCA.all$x[69,3],fossil.hatchling.pcscores[2,3]), inherit = F,
mode = "lines", type = "scatter3d", line =
list( line=list(color='black'),color = "black",dash =
'longdashdot'), opacity = 1,showlegend = F)%>%
```

```
  add_trace(x =
~c(PCA.all$x[67,1],fossil.hatchling.pcscores[3,1]), y =
~c(PCA.all$x[67,2],fossil.hatchling.pcscores[3,2]), z =
~c(PCA.all$x[67,3],fossil.hatchling.pcscores[3,3]), inherit = F,
mode = "markers", type = "scatter3d", marker =
list( line=list(width=1,color='darkblue'),color = "darkblue"),
opacity = 1,showlegend = F)%>%
  add_trace(x = ~fossil.hatchling.pcscores[3,1], y =
~fossil.hatchling.pcscores[3,2], z = ~min(PCA.all$x[,3]-0.01),
inherit = F, mode = "markers", type = "scatter3d", marker =
list( line=list(color='#848688'),color = "#848688"), opacity =
0.3,showlegend = F)%>% ###projection point
  add_trace(x =
c(fossil.hatchling.pcscores[3,1],fossil.hatchling.pcscores[3,1])
, y =
c(list(fossil.hatchling.pcscores[3,2]),list(fossil.hatchling.pcs
cores[3,2])), z =
c(list(fossil.hatchling.pcscores[3,3]),rep(list(-0.118561),2)),
inherit = F, mode = "lines", type = "scatter3d",line =
list(color = 'darkblue'),showlegend = F, opacity = 0.5)%>%
###projection line
  add_trace(x =
~c(PCA.all$x[67,1],fossil.hatchling.pcscores[3,1]), y =
~c(PCA.all$x[67,2],fossil.hatchling.pcscores[3,2]), z =
~c(PCA.all$x[67,3],fossil.hatchling.pcscores[3,3]), inherit = F,
mode = "lines", type = "scatter3d", line =
list( line=list(width=1,color='darkblue'),color =
"darkblue",dash = 'longdashdot'), opacity = 1,showlegend = F)%>%
  add_trace(x =
~c(PCA.all$x[67,1],fossil.hatchling.pcscores[4,1]), y =
~c(PCA.all$x[67,2],fossil.hatchling.pcscores[4,2]), z =
~c(PCA.all$x[67,3],fossil.hatchling.pcscores[4,3]), inherit = F,
mode = "markers", type = "scatter3d", marker =
list( line=list(width=1,color='darkblue'),color = "darkblue"),
opacity = 1,showlegend = F)%>%
  add_trace(x = ~fossil.hatchling.pcscores[4,1], y =
~fossil.hatchling.pcscores[4,2], z = ~min(PCA.all$x[,3]-0.01),
inherit = F, mode = "markers", type = "scatter3d", marker =
list( line=list(color='#848688'),color = "#848688"), opacity =
0.3,showlegend = F)%>% ###projection point
  add_trace(x =
c(fossil.hatchling.pcscores[4,1],fossil.hatchling.pcscores[4,1])
, y =
c(list(fossil.hatchling.pcscores[4,2]),list(fossil.hatchling.pcs
cores[4,2])), z =
c(list(fossil.hatchling.pcscores[4,3]),rep(list(-0.118561),2)),
inherit = F, mode = "lines", type = "scatter3d",line =
```

```
list(color = 'darkblue'),showlegend = F, opacity = 0.5)%>%
###projection line
  add_trace(x =
~c(PCA.all$x[67,1],fossil.hatchling.pcscores[4,1]), y =
~c(PCA.all$x[67,2],fossil.hatchling.pcscores[4,2]), z =
~c(PCA.all$x[67,3],fossil.hatchling.pcscores[4,3]), inherit = F,
mode = "lines", type = "scatter3d", line =
list( line=list(width=1,color='darkblue'),color =
"darkblue",dash = 'longdashdot'), opacity = 1,showlegend = F)%>%
  add_trace(x =
~c(PCA.all$x[67,1],fossil.hatchling.pcscores[5,1]), y =
~c(PCA.all$x[67,2],fossil.hatchling.pcscores[5,2]), z =
~c(PCA.all$x[67,3],fossil.hatchling.pcscores[5,3]), inherit = F,
mode = "markers", type = "scatter3d", marker =
list( line=list(width=1,color='darkblue'),color = "darkblue"),
opacity = 1,showlegend = F)%>%
  add_trace(x = ~fossil.hatchling.pcscores[5,1], y =
~fossil.hatchling.pcscores[5,2], z = ~min(PCA.all$x[,3]-0.01),
inherit = F, mode = "markers", type = "scatter3d", marker =
list( line=list(color='#848688'),color = "#848688"), opacity =
0.3,showlegend = F)%>% ###projection point
  add_trace(x =
c(fossil.hatchling.pcscores[5,1],fossil.hatchling.pcscores[5,1])
, y =
c(list(fossil.hatchling.pcscores[5,2]),list(fossil.hatchling.pcs
cores[5,2])), z =
c(list(fossil.hatchling.pcscores[5,3]),rep(list(-0.118561),2)),
inherit = F, mode = "lines", type = "scatter3d",line =
list(color = 'darkblue'),showlegend = F, opacity = 0.5)%>%
###projection line
  add_trace(x =
~c(PCA.all$x[67,1],fossil.hatchling.pcscores[5,1]), y =
~c(PCA.all$x[67,2],fossil.hatchling.pcscores[5,2]), z =
~c(PCA.all$x[67,3],fossil.hatchling.pcscores[5,3]), inherit = F,
mode = "lines", type = "scatter3d", line =
list( line=list(width=1,color='darkblue'),color =
"darkblue",dash = 'longdashdot'), opacity = 1,showlegend = F)%>%
  add_trace(x = ~PCA.all$x[67:68,1], y = ~PCA.all$x[67:68,2], z
= ~PCA.all$x[67:68,3], inherit = F, mode = "markers", type =
"scatter3d", marker =
list( line=list(width=1,color='black'),color = "darkblue"),
opacity = 1,showlegend = F)%>%
  add_trace(x = ~PCA.all$x[67,1], y = ~PCA.all$x[67,2], z =
~min(PCA.all$x[,3]-0.01), inherit = F, mode = "markers", type =
"scatter3d", marker = list( line=list(color='#848688'),color =
"#848688"), opacity = 0.3,showlegend = F)%>% ###projection point
  add_trace(x = ~PCA.all$x[68,1], y = ~PCA.all$x[68,2], z =
```

```
~min(PCA.all$x[,3]-0.01), inherit = F, mode = "markers", type =
"scatter3d", marker = list( line=list(color='#848688'),color =
"#848688"), opacity = 0.3,showlegend = F)%>% ###projection point
  add_trace(x = c(PCA.all$x[67,1],PCA.all$x[67,1]), y =
c(list(PCA.all$x[67,2]),list(PCA.all$x[67,2])), z =
c(list(PCA.all$x[67,3]),rep(list(-0.118561),2)), inherit = F,
mode = "lines", type = "scatter3d",line = list(color =
'darkblue'),showlegend = F, opacity = 0.5)%>% ###projection line
  add_trace(x = c(PCA.all$x[68,1],PCA.all$x[68,1]), y =
c(list(PCA.all$x[68,2]),list(PCA.all$x[68,2])), z =
c(list(PCA.all$x[68,3]),rep(list(-0.118561),2)), inherit = F,
mode = "lines", type = "scatter3d",line = list(color =
'darkblue'),showlegend = F, opacity = 0.5)%>% ###projection line
  add_trace(x =
~c(ancestral.pcscores[4,1],ancestral.pcscores.adults[4,1]), y =
~c(ancestral.pcscores[4,2],ancestral.pcscores.adults[4,2]), z =
~c(ancestral.pcscores[4,3],ancestral.pcscores.adults[4,3]),
inherit = F, mode = "markers", type = "scatter3d", marker =
list( line=list(width=1,dash =
'longdashdot',color='black'),color = "white"), opacity =
1,showlegend = F)%>%
  add_trace(x = ~ancestral.pcscores[4,1], y =
~ancestral.pcscores[4,2], z = ~min(PCA.all$x[,3]-0.01), inherit
= F, mode = "markers", type = "scatter3d", marker =
list( line=list(color='#848688'),color = "#848688"), opacity =
0.3,showlegend = F)%>% ###projection point
  add_trace(x =
c(ancestral.pcscores[4,1],ancestral.pcscores[4,1]), y =
c(list(ancestral.pcscores[4,2]),list(ancestral.pcscores[4,2])),
z = c(list(ancestral.pcscores[4,3]),rep(list(-0.118561),2)),
inherit = F, mode = "lines", type = "scatter3d",line =
list(color = 'black'),showlegend = F, opacity = 0.5)%>%
###projection line
  add_trace(x = ~ancestral.pcscores.adults[4,1], y =
~ancestral.pcscores.adults[4,2], z = ~min(PCA.all$x[,3]-0.01),
inherit = F, mode = "markers", type = "scatter3d", marker =
list( line=list(color='#848688'),color = "#848688"), opacity =
0.3,showlegend = F)%>% ###projection point
  add_trace(x =
c(ancestral.pcscores.adults[4,1],ancestral.pcscores.adults[4,1])
, y =
c(list(ancestral.pcscores.adults[4,2]),list(ancestral.pcscores.a
dults[4,2])), z =
c(list(ancestral.pcscores.adults[4,3]),rep(list(-0.118561),2)),
inherit = F, mode = "lines", type = "scatter3d",line =
list(color = 'black'),showlegend = F, opacity = 0.5)%>%
###projection line
```

```
  add_trace(x =
~c(ancestral.pcscores[4,1],ancestral.pcscores.adults[4,1]), y =
~c(ancestral.pcscores[4,2],ancestral.pcscores.adults[4,2]), z =
~c(ancestral.pcscores[4,3],ancestral.pcscores.adults[4,3]),
inherit = F, mode = "lines", type = "scatter3d", line =
list( line=list(width=1,color='black'),color = "black",dash =
'longdashdot'), opacity = 1,showlegend = F)%>%
  add_trace(x =
~c(ancestral.pcscores[5,1],ancestral.pcscores.adults[5,1]), y =
~c(ancestral.pcscores[5,2],ancestral.pcscores.adults[5,2]), z =
~c(ancestral.pcscores[5,3],ancestral.pcscores.adults[5,3]),
inherit = F, mode = "markers", type = "scatter3d", marker =
list( line=list(width=1,dash =
'longdashdot',color='black'),color = "white"), opacity =
1,showlegend = F)%>%
  add_trace(x =
c(ancestral.pcscores[5,1],ancestral.pcscores[5,1]), y =
c(list(ancestral.pcscores[5,2]),list(ancestral.pcscores[5,2])),
z = c(list(ancestral.pcscores[5,3]),rep(list(-0.118561),2)),
inherit = F, mode = "lines", type = "scatter3d",line =
list(color = 'black'),showlegend = F, opacity = 0.5)%>%
###projection line
  add_trace(x =
c(ancestral.pcscores.adults[5,1],ancestral.pcscores.adults[5,1])
, y =
c(list(ancestral.pcscores.adults[5,2]),list(ancestral.pcscores.a
dults[5,2])), z =
c(list(ancestral.pcscores.adults[5,3]),rep(list(-0.118561),2)),
inherit = F, mode = "lines", type = "scatter3d",line =
list(color = 'black'),showlegend = F, opacity = 0.5)%>%
###projection line
  add_trace(x = ~ancestral.pcscores[5,1], y =
~ancestral.pcscores[5,2], z = ~min(PCA.all$x[,3]-0.01), inherit
= F, mode = "markers", type = "scatter3d", marker =
list( line=list(color='#848688'),color = "#848688"), opacity =
0.3,showlegend = F)%>% ###projection point
  add_trace(x = ~ancestral.pcscores.adults[5,1], y =
~ancestral.pcscores.adults[5,2], z = ~min(PCA.all$x[,3]-0.01),
inherit = F, mode = "markers", type = "scatter3d", marker =
list( line=list(color='#848688'),color = "#848688"), opacity =
0.3,showlegend = F)%>% ###projection point
  add_trace(x =
~c(ancestral.pcscores[5,1],ancestral.pcscores.adults[5,1]), y =
~c(ancestral.pcscores[5,2],ancestral.pcscores.adults[5,2]), z =
~c(ancestral.pcscores[5,3],ancestral.pcscores.adults[5,3]),
inherit = F, mode = "lines", type = "scatter3d", line =
list( line=list(width=1,color='black'),color = "black",dash =
```

```
'longdashdot'), opacity = 1,showlegend = F)%>%
  add_trace(x =
~c(ancestral.pcscores[18,1],ancestral.pcscores.adults[18,1]), y
= ~c(ancestral.pcscores[18,2],ancestral.pcscores.adults[18,2]),
z =
~c(ancestral.pcscores[18,3],ancestral.pcscores.adults[18,3]),
inherit = F, mode = "markers", type = "scatter3d", marker =
list( line=list(width=1,dash =
'longdashdot',color='black'),color = "white"), opacity =
1,showlegend = F)%>%
  add_trace(x = ~ancestral.pcscores[18,1], y =
~ancestral.pcscores[18,2], z = ~min(PCA.all$x[,3]-0.01), inherit
= F, mode = "markers", type = "scatter3d", marker =
list( line=list(color='#848688'),color = "#848688"), opacity =
0.3,showlegend = F)%>% ###projection point
  add_trace(x = ~ancestral.pcscores.adults[18,1], y =
~ancestral.pcscores.adults[18,2], z = ~min(PCA.all$x[,3]-0.01),
inherit = F, mode = "markers", type = "scatter3d", marker =
list( line=list(color='#848688'),color = "#848688"), opacity =
0.3,showlegend = F)%>% ###projection point
  add_trace(x =
c(ancestral.pcscores[18,1],ancestral.pcscores[18,1]), y =
c(list(ancestral.pcscores[18,2]),list(ancestral.pcscores[18,2]))
, z = c(list(ancestral.pcscores[18,3]),rep(list(-0.118561),2)),
inherit = F, mode = "lines", type = "scatter3d",line =
list(color = 'black'),showlegend = F, opacity = 0.5)%>%
###projection line
  add_trace(x =
c(ancestral.pcscores.adults[18,1],ancestral.pcscores.adults[18,1
]), y =
c(list(ancestral.pcscores.adults[18,2]),list(ancestral.pcscores.
adults[18,2])), z =
c(list(ancestral.pcscores.adults[18,3]),rep(list(-0.118561),2)),
inherit = F, mode = "lines", type = "scatter3d",line =
list(color = 'black'),showlegend = F, opacity = 0.5)%>%
###projection line
  add_trace(x =
~c(ancestral.pcscores[18,1],ancestral.pcscores.adults[18,1]), y
= ~c(ancestral.pcscores[18,2],ancestral.pcscores.adults[18,2]),
z =
~c(ancestral.pcscores[18,3],ancestral.pcscores.adults[18,3]),
inherit = F, mode = "lines", type = "scatter3d", line =
list( line=list(width=1,color='black'),color = "black",dash =
'longdashdot'), opacity = 1,showlegend = F)%>%
  add_trace(x =
~c(ancestral.pcscores[19,1],ancestral.pcscores.adults[19,1]), y
= ~c(ancestral.pcscores[19,2],ancestral.pcscores.adults[19,2]),
```

```
z =
~c(ancestral.pcscores[19,3],ancestral.pcscores.adults[19,3]),
inherit = F, mode = "markers", type = "scatter3d", marker =
list( line=list(width=1,dash =
'longdashdot',color='black'),color = "white"), opacity =
1,showlegend = F)%>%
  add_trace(x = ~ancestral.pcscores[19,1], y =
~ancestral.pcscores[19,2], z = ~min(PCA.all$x[,3]-0.01), inherit
= F, mode = "markers", type = "scatter3d", marker =
list( line=list(color='#848688'),color = "#848688"), opacity =
0.3,showlegend = F)%>% ###projection point
  add_trace(x = ~ancestral.pcscores.adults[19,1], y =
~ancestral.pcscores.adults[19,2], z = ~min(PCA.all$x[,3]-0.01),
inherit = F, mode = "markers", type = "scatter3d", marker =
list( line=list(color='#848688'),color = "#848688"), opacity =
0.3,showlegend = F)%>% ###projection point
  add_trace(x =
c(ancestral.pcscores[19,1],ancestral.pcscores[19,1]), y =
c(list(ancestral.pcscores[19,2]),list(ancestral.pcscores[19,2]))
, z = c(list(ancestral.pcscores[19,3]),rep(list(-0.118561),2)),
inherit = F, mode = "lines", type = "scatter3d",line =
list(color = 'black'),showlegend = F, opacity = 0.5)%>%
###projection line
  add_trace(x =
c(ancestral.pcscores.adults[19,1],ancestral.pcscores.adults[19,1
]), y =
c(list(ancestral.pcscores.adults[19,2]),list(ancestral.pcscores.
adults[19,2])), z =
c(list(ancestral.pcscores.adults[19,3]),rep(list(-0.118561),2)),
inherit = F, mode = "lines", type = "scatter3d",line =
list(color = 'black'),showlegend = F, opacity = 0.5)%>%
###projection line
  add_trace(x =
~c(ancestral.pcscores[19,1],ancestral.pcscores.adults[19,1]), y
= ~c(ancestral.pcscores[19,2],ancestral.pcscores.adults[19,2]),
z =
~c(ancestral.pcscores[19,3],ancestral.pcscores.adults[19,3]),
inherit = F, mode = "lines", type = "scatter3d", line =
list( line=list(width=1,color='black'),color = "black",dash =
'longdashdot'), opacity = 1,showlegend = F)

####without shadows 3d plot
plot_ly(array.2,x=
~PCA.all$x[subset,1],y=~PCA.all$x[subset,2],z=~PCA.all$x[subset,
3], marker = list(opacity =1,line=list(width=1,color='black')),
        type = 'scatter3d',mode = 'markers',color =
as.factor(interaction(families.subset,age.subset)),opacity =
```

```
0.5,
        colors=c("#4E99B7","#4EB773","#a6ccdb","#a6dbb9"))%>%
  layout(scene = list(xaxis = list(title = 'PC1 (48.9%)',
showbackground = TRUE, backgroundcolor = "rgba(232,232,232,1)"),
                      yaxis = list(title = 'PC2 (16.01%)',
showbackground = TRUE, backgroundcolor = "rgba(211,211,211,1)"),
                      zaxis = list(title = 'PC3 (8.82%)')))%>%
  add_trace(x = ~PCA.all$x[33:34,1], y = ~PCA.all$x[33:34,2], z
= ~PCA.all$x[33:34,3], inherit = F, mode = "lines", type =
"scatter3d", line = list( color = "#4E99B7"), opacity =
0.5,showlegend = F)%>%
  add_trace(x = ~PCA.all$x[61:62,1], y = ~PCA.all$x[61:62,2], z
= ~PCA.all$x[61:62,3], inherit = F, mode = "markers", type =
"scatter3d", marker =
list( line=list(width=1,color='black'),color = "#E5CA7F"),
opacity = 1,showlegend = F)%>%
  add_trace(x = ~PCA.all$x[61:62,1], y = ~PCA.all$x[61:62,2], z
= ~PCA.all$x[61:62,3], inherit = F, mode = "lines", type =
"scatter3d", line = list( color = "#E5CA7F"), opacity =
1,showlegend = F)%>%
  add_trace(x = ~PCA.all$x[72:73,1], y = ~PCA.all$x[72:73,2], z
= ~PCA.all$x[72:73,3], inherit = F, mode = "markers", type =
"scatter3d", marker =
list( line=list(width=1,color='black'),color = "pink"), opacity
= 1,showlegend = F)%>%
  add_trace(x = ~PCA.all$x[72:73,1], y = ~PCA.all$x[72:73,2], z
= ~PCA.all$x[72:73,3], inherit = F, mode = "lines", type =
"scatter3d", line = list( color = "pink"), opacity =
1,showlegend = F)%>%
  add_trace(x = ~PCA.all$x[63:64,1], y = ~PCA.all$x[63:64,2], z
= ~PCA.all$x[63:64,3], inherit = F, mode = "markers", type =
"scatter3d", marker =
list( line=list(width=1,color='black'),color = "#D84C71"),
opacity = 1,showlegend = F)%>%
  add_trace(x = ~PCA.all$x[63:64,1], y = ~PCA.all$x[63:64,2], z
= ~PCA.all$x[63:64,3], inherit = F, mode = "lines", type =
"scatter3d", line = list( color = "#D84C71"), opacity =
1,showlegend = F)%>%
  add_trace(x = ~PCA.all$x[65:66,1], y = ~PCA.all$x[65:66,2], z
= ~PCA.all$x[65:66,3], inherit = F, mode = "markers", type =
"scatter3d", marker =
list( line=list(width=1,color='black'),color = "#855BC1"),
opacity = 1,showlegend = F)%>%
  add_trace(x = ~PCA.all$x[65:66,1], y = ~PCA.all$x[65:66,2], z
= ~PCA.all$x[65:66,3], inherit = F, mode = "lines", type =
"scatter3d", line = list( color = "#855BC1"), opacity =
1,showlegend = F)%>%
```

```
  add_trace(x =
~c(PCA.all$x[69,1],fossil.hatchling.pcscores[1,1]), y =
~c(PCA.all$x[69,2],fossil.hatchling.pcscores[1,2]), z =
~c(PCA.all$x[69,3],fossil.hatchling.pcscores[1,3]), inherit = F,
mode = "markers", type = "scatter3d", marker =
list( line=list(width=1,color='black'),color = "black"), opacity
= 1,showlegend = F)%>%
  add_trace(x =
~c(PCA.all$x[69,1],fossil.hatchling.pcscores[1,1]), y =
~c(PCA.all$x[69,2],fossil.hatchling.pcscores[1,2]), z =
~c(PCA.all$x[69,3],fossil.hatchling.pcscores[1,3]), inherit = F,
mode = "lines", type = "scatter3d", line =
list( line=list(width=1,color='black'),color = "black",dash =
'longdashdot'), opacity = 1,showlegend = F)%>%
  add_trace(x =
~c(PCA.all$x[69,1],fossil.hatchling.pcscores[2,1]), y =
~c(PCA.all$x[69,2],fossil.hatchling.pcscores[2,2]), z =
~c(PCA.all$x[69,3],fossil.hatchling.pcscores[2,3]), inherit = F,
mode = "markers", type = "scatter3d", marker =
list( line=list(width=1,color='black'),color = "black"), opacity
= 1,showlegend = F)%>%
  add_trace(x =
~c(PCA.all$x[69,1],fossil.hatchling.pcscores[2,1]), y =
~c(PCA.all$x[69,2],fossil.hatchling.pcscores[2,2]), z =
~c(PCA.all$x[69,3],fossil.hatchling.pcscores[2,3]), inherit = F,
mode = "lines", type = "scatter3d", line =
list( line=list(color='black'),color = "black",dash =
'longdashdot'), opacity = 1,showlegend = F)%>%
  add_trace(x =
~c(PCA.all$x[67,1],fossil.hatchling.pcscores[3,1]), y =
~c(PCA.all$x[67,2],fossil.hatchling.pcscores[3,2]), z =
~c(PCA.all$x[67,3],fossil.hatchling.pcscores[3,3]), inherit = F,
mode = "markers", type = "scatter3d", marker =
list( line=list(width=1,color='darkblue'),color = "darkblue"),
opacity = 1,showlegend = F)%>%
  add_trace(x =
~c(PCA.all$x[67,1],fossil.hatchling.pcscores[3,1]), y =
~c(PCA.all$x[67,2],fossil.hatchling.pcscores[3,2]), z =
~c(PCA.all$x[67,3],fossil.hatchling.pcscores[3,3]), inherit = F,
mode = "lines", type = "scatter3d", line =
list( line=list(width=1,color='darkblue'),color =
"darkblue",dash = 'longdashdot'), opacity = 1,showlegend = F)%>%
  add_trace(x =
~c(PCA.all$x[67,1],fossil.hatchling.pcscores[4,1]), y =
~c(PCA.all$x[67,2],fossil.hatchling.pcscores[4,2]), z =
~c(PCA.all$x[67,3],fossil.hatchling.pcscores[4,3]), inherit = F,
mode = "markers", type = "scatter3d", marker =
```

```
list( line=list(width=1,color='darkblue'),color = "darkblue"),
opacity = 1,showlegend = F)%>%
  add_trace(x =
~c(PCA.all$x[67,1],fossil.hatchling.pcscores[4,1]), y =
~c(PCA.all$x[67,2],fossil.hatchling.pcscores[4,2]), z =
~c(PCA.all$x[67,3],fossil.hatchling.pcscores[4,3]), inherit = F,
mode = "lines", type = "scatter3d", line =
list( line=list(width=1,color='darkblue'),color =
"darkblue",dash = 'longdashdot'), opacity = 1,showlegend = F)%>%
  add_trace(x =
~c(PCA.all$x[67,1],fossil.hatchling.pcscores[5,1]), y =
~c(PCA.all$x[67,2],fossil.hatchling.pcscores[5,2]), z =
~c(PCA.all$x[67,3],fossil.hatchling.pcscores[5,3]), inherit = F,
mode = "markers", type = "scatter3d", marker =
list( line=list(width=1,color='darkblue'),color = "darkblue"),
opacity = 1,showlegend = F)%>%
  add_trace(x =
~c(PCA.all$x[67,1],fossil.hatchling.pcscores[5,1]), y =
~c(PCA.all$x[67,2],fossil.hatchling.pcscores[5,2]), z =
~c(PCA.all$x[67,3],fossil.hatchling.pcscores[5,3]), inherit = F,
mode = "lines", type = "scatter3d", line =
list( line=list(width=1,color='darkblue'),color =
"darkblue",dash = 'longdashdot'), opacity = 1,showlegend = F)%>%
  add_trace(x = ~PCA.all$x[67:68,1], y = ~PCA.all$x[67:68,2], z
= ~PCA.all$x[67:68,3], inherit = F, mode = "markers", type =
"scatter3d", marker =
list( line=list(width=1,color='black'),color = "darkblue"),
opacity = 1,showlegend = F)%>%
  add_trace(x =
~c(ancestral.pcscores[4,1],ancestral.pcscores.adults[4,1]), y =
~c(ancestral.pcscores[4,2],ancestral.pcscores.adults[4,2]), z =
~c(ancestral.pcscores[4,3],ancestral.pcscores.adults[4,3]),
inherit = F, mode = "markers", type = "scatter3d", marker =
list( line=list(width=1,dash =
'longdashdot',color='black'),color = "white"), opacity =
1,showlegend = F)%>%
  add_trace(x =
~c(ancestral.pcscores[4,1],ancestral.pcscores.adults[4,1]), y =
~c(ancestral.pcscores[4,2],ancestral.pcscores.adults[4,2]), z =
~c(ancestral.pcscores[4,3],ancestral.pcscores.adults[4,3]),
inherit = F, mode = "lines", type = "scatter3d", line =
list( line=list(width=1,color='black'),color = "black",dash =
'longdashdot'), opacity = 1,showlegend = F)%>%
  add_trace(x =
~c(ancestral.pcscores[5,1],ancestral.pcscores.adults[5,1]), y =
~c(ancestral.pcscores[5,2],ancestral.pcscores.adults[5,2]), z =
~c(ancestral.pcscores[5,3],ancestral.pcscores.adults[5,3]),
```

```
inherit = F, mode = "markers", type = "scatter3d", marker =
list( line=list(width=1,dash =
'longdashdot',color='black'),color = "white"), opacity =
1,showlegend = F)%>%
  add_trace(x =
~c(ancestral.pcscores[5,1],ancestral.pcscores.adults[5,1]), y =
~c(ancestral.pcscores[5,2],ancestral.pcscores.adults[5,2]), z =
~c(ancestral.pcscores[5,3],ancestral.pcscores.adults[5,3]),
inherit = F, mode = "lines", type = "scatter3d", line =
list( line=list(width=1,color='black'),color = "black",dash =
'longdashdot'), opacity = 1,showlegend = F)%>%
  add_trace(x =
~c(ancestral.pcscores[18,1],ancestral.pcscores.adults[18,1]), y
= ~c(ancestral.pcscores[18,2],ancestral.pcscores.adults[18,2]),
z =
~c(ancestral.pcscores[18,3],ancestral.pcscores.adults[18,3]),
inherit = F, mode = "markers", type = "scatter3d", marker =
list( line=list(width=1,dash =
'longdashdot',color='black'),color = "white"), opacity =
1,showlegend = F)%>%
  add_trace(x =
~c(ancestral.pcscores[18,1],ancestral.pcscores.adults[18,1]), y
= ~c(ancestral.pcscores[18,2],ancestral.pcscores.adults[18,2]),
z =
~c(ancestral.pcscores[18,3],ancestral.pcscores.adults[18,3]),
inherit = F, mode = "lines", type = "scatter3d", line =
list( line=list(width=1,color='black'),color = "black",dash =
'longdashdot'), opacity = 1,showlegend = F)%>%
  add_trace(x =
~c(ancestral.pcscores[19,1],ancestral.pcscores.adults[19,1]), y
= ~c(ancestral.pcscores[19,2],ancestral.pcscores.adults[19,2]),
z =
~c(ancestral.pcscores[19,3],ancestral.pcscores.adults[19,3]),
inherit = F, mode = "markers", type = "scatter3d", marker =
list( line=list(width=1,dash =
'longdashdot',color='black'),color = "white"), opacity =
1,showlegend = F)%>%
  add_trace(x =
~c(ancestral.pcscores[19,1],ancestral.pcscores.adults[19,1]), y
= ~c(ancestral.pcscores[19,2],ancestral.pcscores.adults[19,2]),
z =
~c(ancestral.pcscores[19,3],ancestral.pcscores.adults[19,3]),
inherit = F, mode = "lines", type = "scatter3d", line =
list( line=list(width=1,color='black'),color = "black",dash =
'longdashdot'), opacity = 1,showlegend = F)
```
